# An Autonomous Implantable Device for the Prevention of Death from Opioid Overdose

**DOI:** 10.1101/2024.06.27.600919

**Authors:** Joanna L. Ciatti, Abraham Vazquez-Guardado, Victoria E. Brings, Jihun Park, Brian Ruyle, Rebecca A. Ober, Alicia J. McLuckie, Michael R. Talcott, Emily A. Carter, Amy R. Burrell, Rebecca A. Sponenburg, Jacob Trueb, Prashant Gupta, Joohee Kim, Raudel Avila, Minho Seong, Richard A. Slivicki, Melanie A. Kaplan, Bryan Villalpando-Hernandez, Nicolas Massaly, Michael C. Montana, Mitchell Pet, Yonggang Huang, Jose A. Morón, Robert W. Gereau, John A. Rogers

## Abstract

Opioid overdose accounts for nearly 75,000 deaths per year in the United States, representing a leading cause of mortality amongst the prime working age population (25-54 years). At overdose levels, opioid-induced respiratory depression becomes fatal without timely administration of the rescue drug naloxone. Currently, overdose survival relies entirely on bystander intervention, requiring a nearby person to discover and identify the overdosed individual, and have immediate access to naloxone to administer. Government efforts have focused on providing naloxone in abundance but do not address the equally critical component for overdose rescue: a willing and informed bystander. To address this unmet need, we developed the Naloximeter: a class of life-saving implantable devices that autonomously detect and treat overdose, with the ability to simultaneously contact first-responders. We present three Naloximeter platforms, for both fundamental research and clinical translation, all equipped with optical sensors, drug delivery mechanisms, and a supporting ecosystem of technology to counteract opioid-induced respiratory depression. In small and large animal studies, the Naloximeter rescues from otherwise fatal opioid overdose within minutes. This work introduces life-changing, clinically translatable technologies that broadly benefit a susceptible population recovering from opioid use disorder.

## Introduction

Opioids have been used for medicinal purposes for centuries and remain the primary analgesic to treat moderate to severe acute pain (*1*, *2*). Opioid use disorder (OUD), characterized by persistent opioid usage despite its adverse consequences, is a chronic relapse disorder and is one of the major contributors to global disease burden, with over 40 million people dependent on opioids worldwide in 2017 (*3–5*). A rise in drug-involved overdose deaths in the United States has coincided with the widespread availability of synthetic opioids, e.g. fentanyl, which accounted for approximately 75,000 deaths in 2023, roughly 70% of all drug overdose deaths (*6–9*). The age-adjusted drug overdose mortality rate has nearly tripled over the last ten years (*10*). Medically supervised withdrawal, or detoxification, is the standard of care for OUD treatment (*4*). This population has, however, a risk of fatal overdose that is elevated by 10 – 16 times within the first several months following a period of sobriety (*11–15*). Further, 5–10% of patients hospitalized for a drug overdose will die within one year of discharge (*16*, *17*). Therefore, high-risk individuals recovering from OUD require solutions to prevent fatalities in the event of relapse.

Opioid-induced respiratory depression (OIRD) is a dose-dependent phenomenon that can progress to complete apnea without intervention. This can lead to overdose fatality, due to neuronal damage caused by disruption of oxygen supply to the brain and other vital organs (*18–21*). Thus, the timely detection of overdose is critical to minimizing the risk of death. Current treatment for overdose involves administration of the rescue drug naloxone (NLX), an opioid receptor antagonist that temporarily reverses the effect of opioids and restores breathing (*19*, *22*). Overdose reversal with NLX is highly effective; however, deployment to an acutely overdosing patient requires timely identification of overdose, the immediate availability of NLX, and proper training of the bystander to administer (*23*, *24*). Given that overdoses often occur when the patient is alone, hypoxic brain injury and death are the most common outcomes (*25*, *26*). Public-health solutions including supervised injection facilities (*27*, *28*), virtual overdose monitoring systems (*29–33*), or fixed-location overdose detection devices (*34–36*) are of growing interest, but these solutions are slow to implement and costly, and most importantly still require human intervention to deliver NLX.

To address this challenge, we envisioned a closed-loop rescue device for the rapid detection and autonomous treatment of opioid overdose, which obviates the need for bystander intervention. A few devices designed to detect overdose and deliver NLX have been proposed recently, but their bulky form-factors, impracticality, and lack of *in-vivo* validation studies suggest limited real-world relevance (*37–39*). Wearable devices for overdose detection are also of interest (*40–43*), but as mentioned, they lack autonomous rescue capabilities. We introduce a device platform that includes physiological sensors with several options for automated drug delivery, power supply and wireless communications. An accompanying software system supports real-time data analytics and closed-loop control. The technology, referred to in the following as a Naloximeter, addresses both research and clinical applications, as demonstrated in benchtop investigations and *in-vivo* demonstrations of automatic overdose rescue in rodent and porcine models. The design choices emphasize engineering simplicity and capability for clinical translation, where large animal demonstrations highlight its potential life-saving benefits for humans.

## Results

### Implantable device for automated closed-loop rescue from opioid overdose

The Naloximeter is a subcutaneously implanted device that utilizes (1) a dual-wavelength optical sensor for measurements of local tissue oxygenation (StO_2_) to detect physiological signs of an overdose and (2) an automated drug delivery system to execute a closed-loop, rapid rescue response (Fig. 1A). The optical sensor operates at red (660 nm) and near-infrared (880 nm) wavelengths, to yield time-series data that is analyzed with a multivariable algorithm for overdose detection including calculations of StO_2_, based on established methods of near-infrared spectroscopy (*44*), along with other features in the data. The system microcontroller communicates via Bluetooth Low Energy (BLE) protocols to a cellular-enabled device (phone or tablet) with a custom mobile application to control the overall operation.

**Fig. 1.**
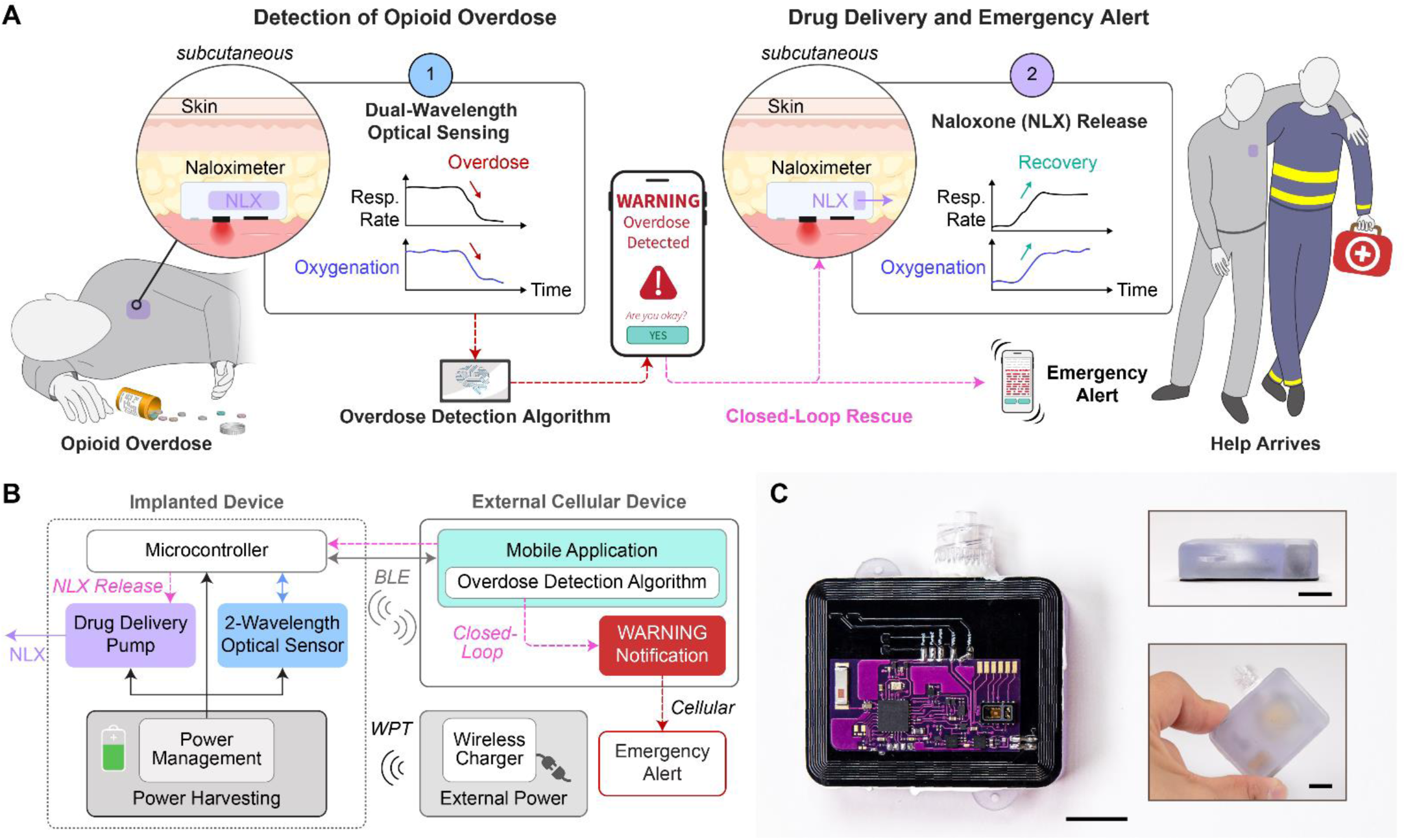
Autonomous operational scheme of Naloximeter for continuous sensing, triggered drug delivery and rescue from opioid overdose. (**A**) Implanted device (Naloximeter) detects respiratory depression leading to hypoxia, a signature of opioid overdose, delivers the rescue drug naloxone (NLX) and sends an emergency alert to first responders. (**B**) Operational diagram including transdermal wireless power transfer, bidirectional communication via Bluetooth Low Energy (BLE) protocols, and cellular communication that delivers an emergency alert. (**C**) Optical images of a Naloximeter, *inset:* side (top) and top view (bottom) of the device. Scale bars, 1 cm.

The closed-loop functionality is a response cascade that begins with the continuous analysis of data from the optical sensor to monitor for signatures of an overdose, followed by rapid NLX delivery and transmission of an emergency alert to first responders. A warning notification on the cellular device allows the user to abort the rescue process as a fail-safe mechanism in the case of a falsely detected overdose event. In an embodiment designed for large animals and ultimately humans, a lithium polymer (LiPo) battery capable of wireless recharging at the medical frequency band (13.56 MHz) via magnetic induction serves as the power supply (Fig. 1B). A battery-free, near-field communication (NFC) version designed for research with small animal models exploits magnetic induction for wireless communication and continuous power delivery. Additional details on the electronics, software, and related characterizations are in the supplementary text, Fig. S1 - S6, and Table S1. Figure 1C presents images of a Naloximeter device for large animals, the areal footprint (42 × 34 × 13 mm^3^) is approximately 75% the size of a commercial pacemaker (Fig. S7).

We developed two schemes for drug administration, both designed to rapidly deliver NLX to the bloodstream. The intravenous version (Fig. 2A) includes a printed circuit board (PCB) with interdigitated gold electrodes, drug and electrolyte reservoirs separated by an elastomeric membrane, in an enclosure formed in a biocompatible polymer with a catheter connection. Upon activation, electrolytic water splitting generates H_2_ and O_2_ gas in the electrolyte chamber. The resulting pressure deforms the membrane (polystyrene-*b*-polyisoprene-*b*-polystyrene; *E* = 4.3 MPa) and forces NLX from the drug reservoir into the vasculature via the intravenous catheter (Fig. 2B). Parametric modeling of the mechanics of this deformation process, supported by benchtop experimental studies, guides selection of optimized design parameters (Fig. S8 and S9), including a membrane diameter of 24 mm and thickness of 70 μm. The device accommodates a drug volume of 3 mL, capable of complete release of NLX directly into the bloodstream in less than 6 min (Fig. 2C, Supplementary Video 1). The dead volume of the catheter (0.2 mL in Fig. 2C) accounts for a portion of the drug payload that is not infused. The doses quoted in the following correspond to the total volume loaded in the device. Degradation studies of the electrodes show adequate corrosion resistance to ensure stable operation (Supplementary Fig. S10).

**Fig. 2.**
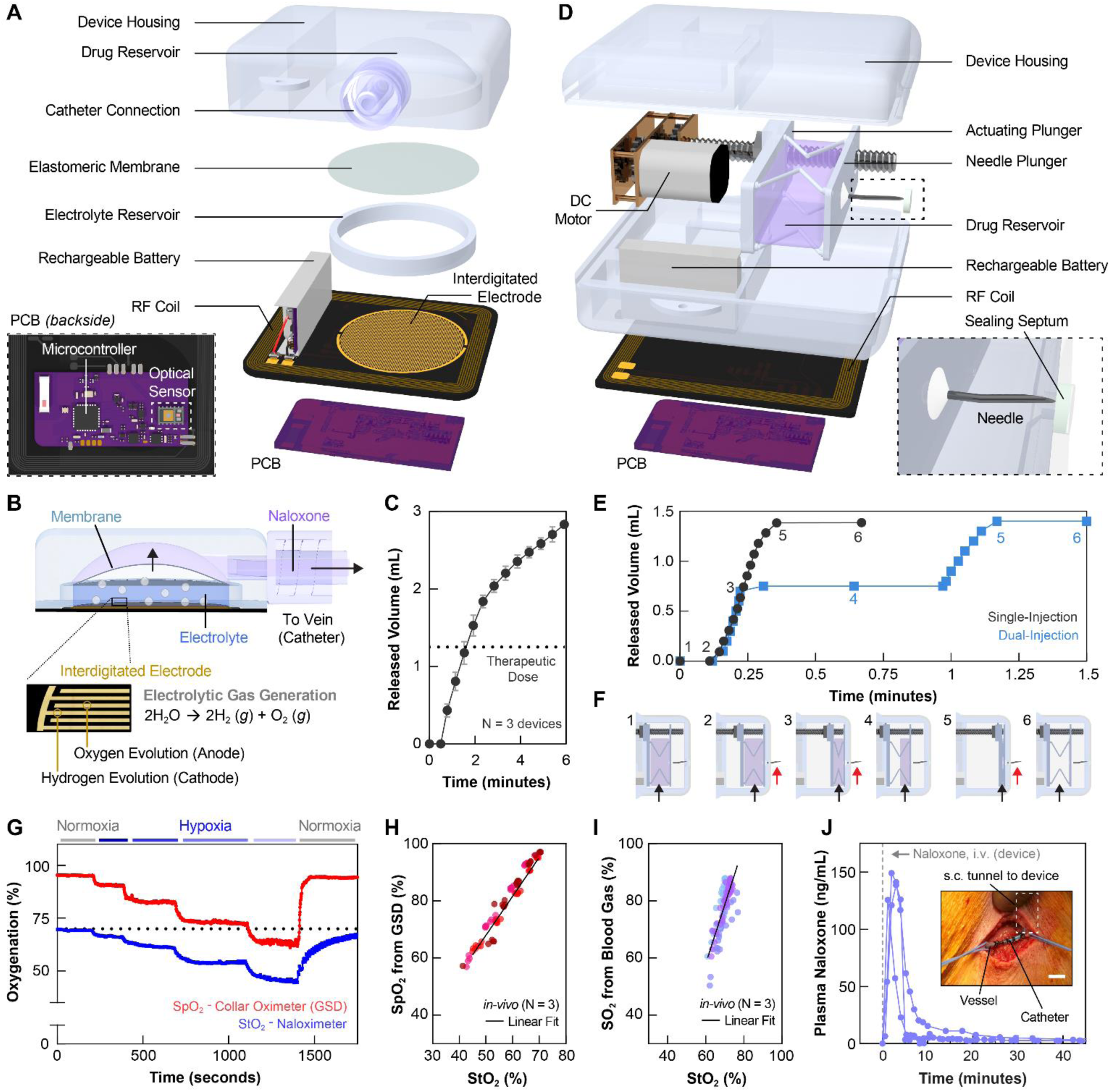
Device design and *in-vivo* validation studies. (**A**) Exploded view illustration of a Naloximeter for intravenous drug delivery via catheter and electrolytic pump, *inset:* printed-circuit board (PCB) layout with critical components labeled including the dual-wavelength optical sensor, mounted on the backside of the assembled pump device. (**B**) Cross sectional illustration showing deformation of a flexible membrane under pressure from gas generated via electrolytic water splitting. This process drives naloxone (NLX) through a catheter port to a vein. (**C**) Drug release from the electrolytic device, *N* = 3 devices, error bars are SD. (**D**) Exploded view illustration of a Naloximeter for drug injection via a needle and motor-driven actuator, *inset:* Huber needle and PTFE septum at the outlet of the housing. (**E**) Drug release from the injector device in single- or dual-injection mode (*N* = 3 devices). (**F**) Schematic illustrations of dual-injection operation, namely: needle deployment (*1–2*), delivery of first dose (*3*), retraction (*4*), delivery of second dose (*5*), and final retraction once empty (*6*). (**G**) Validation of the dual-wavelength optical sensor for determining StO_2_ via sequential exposure to hypoxic gas mixtures (21, 14, 12, 10, 8, 21% O_2_, denoted as colored bars) in a rodent model; SpO_2_ data collected using a gold standard device (GSD; MouseOx collar). (**H**) Correlation plot for SpO_2_ (GSD) and StO_2_ (Naloximeter) resulting from rodent hypoxia validation studies (*R^2^* = 0.95). Colors signify individual animals. (**I**) Correlation plot for SO_2_ from blood gases and StO_2_ from the dual-wavelength optical sensor validated in porcine hypoxia studies (*R^2^* = 0.65). Colors signify individual devices. (**J**) Pharmacokinetics of naloxone (NLX) delivered using the intravenous device in a porcine model, dose: 3 mL (2.7 mg NLX), *N* = 3 animals. *Inset:* Optical image of a catheter tunneled subcutaneously from a device and secured in the jugular vein, scale bar is 1 cm.

The injector version of the Naloximeter (Fig. 2D) includes an actuating plunger, a compressible drug reservoir made of ethylene vinyl alcohol and a Huber needle. A small-scale DC motor drives injection of NLX into the adjacent subcutaneous tissue, puncturing any fibrotic capsule that may form around the device as a foreign body response (*45*). A polytetrafluoroethylene/silicone septum seals the chamber at the needle outlet to prevent flow of biofluid into the enclosure, formed in a biocompatible polymer identical to the intravenous version (Fig. 2D, inset). The force required to deploy and retract the needle through the sealing septum is less than 1.3 N, far below the 10 N force delivered by the motor (Fig. S11 - S12). The capacity of the drug reservoir is 1.5 mL. Complete delivery occurs within 25 s as a single dose, with a loss volume of < 0.1 mL (Fig. 2E). The device can also be operated in dual-injection mode, with two 0.75 mL payloads delivered sequentially, as in Fig. 2E, 2F and Supplementary Video 2. The potency of fentanyl and its analogs often requires multiple naloxone administrations to revive during an overdose event, thus warranting the dual-injection capability of the device (*22*, *23*). Results of additional characterization studies are in Fig. S13. Further details on the device architecture and fabrication appear in the methods section and Fig. S14 - S15.

The electronic hardware and firmware are similar for these two device embodiments; the optical sensor is the same (Fig. 2A inset, Fig. S2). Hypoxia studies in rodent and porcine models with stepwise modulation of inspired oxygen, serve as the basis for validating measurements of StO_2_. In rodents, the calculated StO_2_ values correlate with SpO_2_ determined by a gold standard device (GSD), for SpO_2_ in the range of 60–100% across multiple animals (*N* = 3; Fig. 2G, 2H and S16). Furthermore, comparisons of StO_2_ measurements against oxygen saturation (SO_2_) determined by blood gas analysis in hypoxic porcine models produce similar results (*N* = 3; Fig. 2I and S17 - S18). Pharmacokinetic data for NLX delivered with the intravenous device in porcine models indicate swift delivery into the bloodstream. NLX appears in the plasma within one minute of pump activation, and the peak concentration of 138 ± 10 ng/mL occurs at 2 ± 1 min after activation (*N* = 3; Fig. 2J). The variations can be attributed to differences in catheter length, diameter, and blood circulation between animals. The implanted configuration of the device and catheter appear in Fig. 2J, inset, and Fig. S20. Additional experiments employed a gadolinium (Gd) tracer element found in MRI contrast agents as a proxy for NLX (supplementary materials and Fig. S19).

### Characterization of opioid overdose in large animal models

The physiological manifestation of opioid overdose consists of the gradual reduction, including possible cessation, of oxygen uptake in the lungs (Fig. 3A), driven by respiratory depression (Fig. 3B) and accompanied by an increment in end-tidal carbon dioxide (EtCO_2_, Fig. 3C). Respiratory depression often precipitates cardiac depression, observed as bradycardia (Fig. 3D) and low blood pressure (Fig. 3E). The circulation of excess deoxygenated blood produces systemic oxygen deprivation, as observed with clinical measurements (arterial blood gas: ABG) and optical sensors (Naloximeter: StO_2_, standard pulse oximeter: SpO_2_) in Fig. 3A. This oxygen desaturation occurs on the timescale of several seconds to one minute after fentanyl administration. The minimum dosage of fentanyl that will produce fatal overdose in humans is difficult to predict because the respiratory response depends on the health and opioid tolerance of the individual subject. After fatal overdose, often the only data available are the fentanyl concentrations in plasma, not the dosage that precipitated the overdose (*46*). Nevertheless, the drug enforcement administration (DEA) estimates a lethal dose of approximately 30 µg/kg (2 mg) in humans, and recommends ventilatory support for doses above 10 µg/kg (*47*, *48*). Experiments at five dosages of intravenous fentanyl in the range of 2.5 to 100 µg/kg in porcine models indicates that the oxygenation response is similar for doses above 10 µg/kg, producing a desaturation event with oxygen decline at a rate of ∼ -10.4% StO_2_ per minute (Fig. 3F), as confirmed with ABG analysis (∼ -13.9% SO_2_ per minute, Fig. 3G). The observed desaturation rates produced by these dosages yield severe hypoxia (< 70% SO_2_) within 3 min, with risk of global ischemia and brain injury if sustained over several minutes (*49*). Smaller fentanyl doses lead to reduced desaturation rates (Fig. 3G and S21), but equally fatal outcomes.

**Fig. 3.**
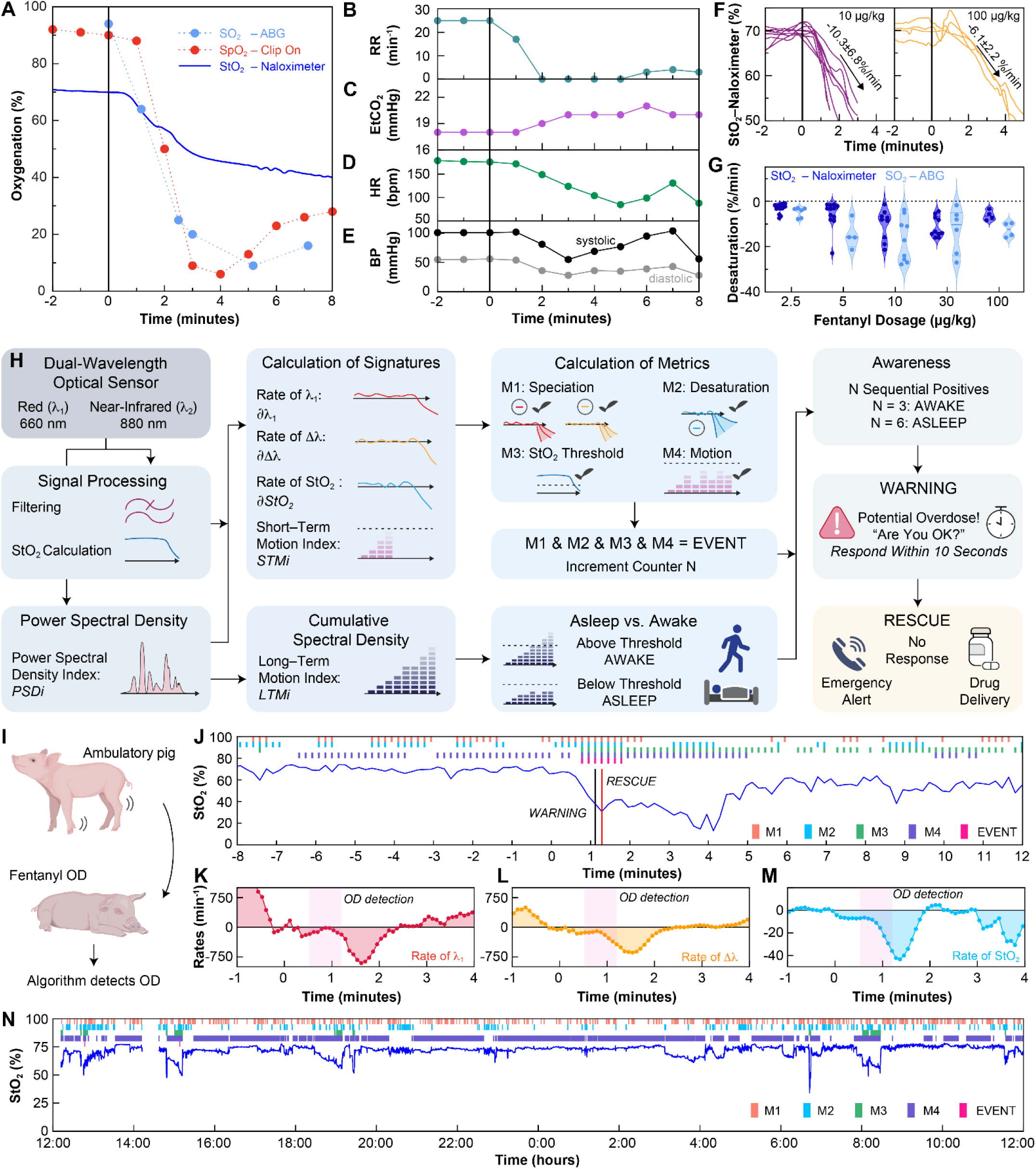
Physiological signs of opioid overdose and an algorithm for overdose detection in freely moving subjects. (**A**) StO_2_ recorded with a Naloximeter during a fentanyl overdose (OD; 10 µg/kg) in an anesthetized pig and its comparison with SO_2_ measured by arterial blood gas (ABG) analysis and SpO_2_ measured by a pulse oximeter (ear clip-on). (**B**) Decline of respiration rate, (**C**) elevation of end-tidal carbon dioxide (EtCO_2_), (**D**) bradycardia, and (**D**) reduction in blood pressure after fentanyl injection (at time = 0) represent clinical signs of overdose. (**F**) StO_2_ recorded from Naloximeters during fentanyl overdose quantify the rate of desaturation: (left) 10 µg/kg, -10.3 ± 6.8 %/min, n = 9 devices; and (right) 100 µg/kg, -6.1 ± 2.2 %/min, n = 4 devices. (**G**) Desaturation rates following fentanyl administration, estimated with data from Naloximeters and ABG analysis at different dosages. N = 29 subjects and n = 52 devices; 100 µg/kg [N, n] = [4, 4], 30 µg/kg [N, n] = [6, 10], 10 µg/kg [N, n] = [9, 9], 5 µg/kg [N, n] = [4, 12], and 2.5 µg/kg [N, n] = [6, 17]. (**H**) Schematic illustration of the analytical approach for overdose detection based on data from the Naloximeter dual-wavelength optical sensor. Various metrics associated to optical absorption and its rate of change (M1), desaturation rate (M2), tissue oxygenation levels (M3), and signal power density (M4) serve as the basis for a multivariable algorithm that discriminates physiological signs of opioid overdose from confounding events associated with physical activity, postural changes, and sleep apnea events. (**I**) Operation of the algorithm on data from a Naloximeter implanted in an ambulatory pig, including (**J**) StO_2_ during opioid overdose (30 µg/kg), and the rates of change of (**K**) the red optical signal (*δλ*_1_), (**L**) the difference between the red and infrared optical signals (*∂*Δ*λ*) of the dual-wavelength optical sensor, and (**M**) the tissue oxygenation (*∂StO*_2_). Fentanyl was administered at time = 0. (**O**) Application of the overdose detection algorithm in a healthy, normally ambulating pig for 24 hours. Although several instances of synchronous positive metrics, i.e. EVENTs (EVENT = M1 & M2 & M3 & M4) were registered, none of them led to a WARNING. Colored bars in **J** and **O** indicate time stamps where individual metrics (M1, M2, M3, and M4) or compound logical metrics (EVENT) are true.

Detecting an overdose with high specificity and selectivity requires accurate interpretation of physiological signatures, while discriminating against false positives induced by motions and other artifacts. At the onset of OIRD, the transition to hypoxia produces changes in tissue optical absorption due to the speciation of deoxy- and oxygenated hemoglobin (*50*). An increase in deoxygenated hemoglobin produces an increase in optical absorption at the red wavelength and a corresponding reduction in the red optical signal at the sensor, with an opposite effect for the near-infrared signal (Fig. S16 and S22). These changes differ from motion-induced artifacts, where rapid fluctuations in both signals produce short-lived variations in the calculated values of StO_2_. Power spectral density analysis of data from humans performing specific movements shows that the reduction of spectral content in the 1-5 Hz range correlates with the reduction of gross body motions, thus serving as an indicator of the intoxicated overdose state (Fig. S23). The overdose detection algorithm (ODA) uses multivariable analysis of data from the optical sensor to operate reliably and quickly (within 90 s of fentanyl administration) to detect overdose and reject false positives during normal ambulation, as illustrated in Fig. 3H. The ODA calculates the StO_2_ and an index of motion (*STMi*, cumulative over 60 s), and it determines the rates of change of the StO_2_ (*∂StO*_2_), the magnitude of the red signal (*δλ*_1_), and the difference between the red and infrared signals (*∂*Δ*λ*). Comparisons of these parameters against threshold values produce four logical metrics every 10 s. The mutually inclusive condition of all four metrics suggests a potential overdose episode (an “EVENT”). A second motion index (*LTMi*, cumulative over 10 min) determines if the subject is awake or asleep (Fig. S24). An uninterrupted successive series of EVENTs detected within 30 s leads to a warning flag (a “WARNING”) that produces a push notification on the mobile app to prompt user engagement. When the subject is asleep, the ODA adjusts the triggering time to 60 s to account for confounding desaturation produced during obstructive sleep apnea. If the user fails to acknowledge this notification within 10 s, the system launches the closed-loop rescue cascade: activating the drug delivery mechanism and relaying an emergency alert. Further details regarding the algorithm are in the supplemental text.

Figure 3I depicts tests of the ODA, performed through multiple fentanyl overdoses with ambulatory porcine models. A dose of 30 µg/kg fentanyl leads to cyanosis, respiratory depression, stiff posture, and muscle tremors in these animals. In one animal, the ODA detected the early signs of overdose (first EVENT) within 37 s after fentanyl administration, as the onset of a steady decrease in StO_2_ (Fig. 3J) accompanied by sequential positive metric conditions that confirm the overdose (Fig. 3K - 3M, S25). A WARNING at 67 s preceded the rescue cascade at 77 s (Fig. 3J). A 24-hour period of continuous monitoring in a freely moving pig serves as the basis for further validation of the ODA (Fig. 3N; with additional datasets, *N* = 3 total, in Fig. S26). This example shows multiple episodes where the StO_2_ approaches clinically dangerous levels, but the absence of sequential occurrences of positive metrics (Fig. S27) avoids false indications of an overdose.

### Device rescues freely moving animals from opioid overdose

Experiments in rodent models validate the closed-loop rescue concept with a miniaturized, battery-free version of the electrolytic Naloximeter designed for research purposes, implemented without the catheter (Fig. 4A, inset). This device operates via NFC protocols and wireless power transfer, as described in the methods section and Fig. S28 and S29. As with pigs, administration of a sufficient dose of fentanyl in rats leads to a rapid desaturation event. The NFC Naloximeter detects signs of an overdose within 1 min, and then triggers a rescue cascade as described previously, with an emergency alert received at 70 s after fentanyl administration. The animals show signs of recovery within 4 min and full recovery (90% SpO_2_) after 4.5 min (*N* = 3; Fig. 4B). Comparisons of overdose recovery between groups rescued by the Naloximeter, manual subcutaneous NLX injection, or without administration of NLX (self-recovery) highlight the rescue capabilities of the device (Fig. 4C). Overdose recovery time is highly significant (*P* < 0.0001) between the Naloximeter and self-recovery (*N* = 3 and 4; recovery times = 4.5 ± 0.3 and 23 ± 4 min, respectively). The same is true between manual injection (*N* = 4; recovery time = 3.7 ± 0.6 min) and self-recovery (*P* < 0.0001).

**Fig. 4.**
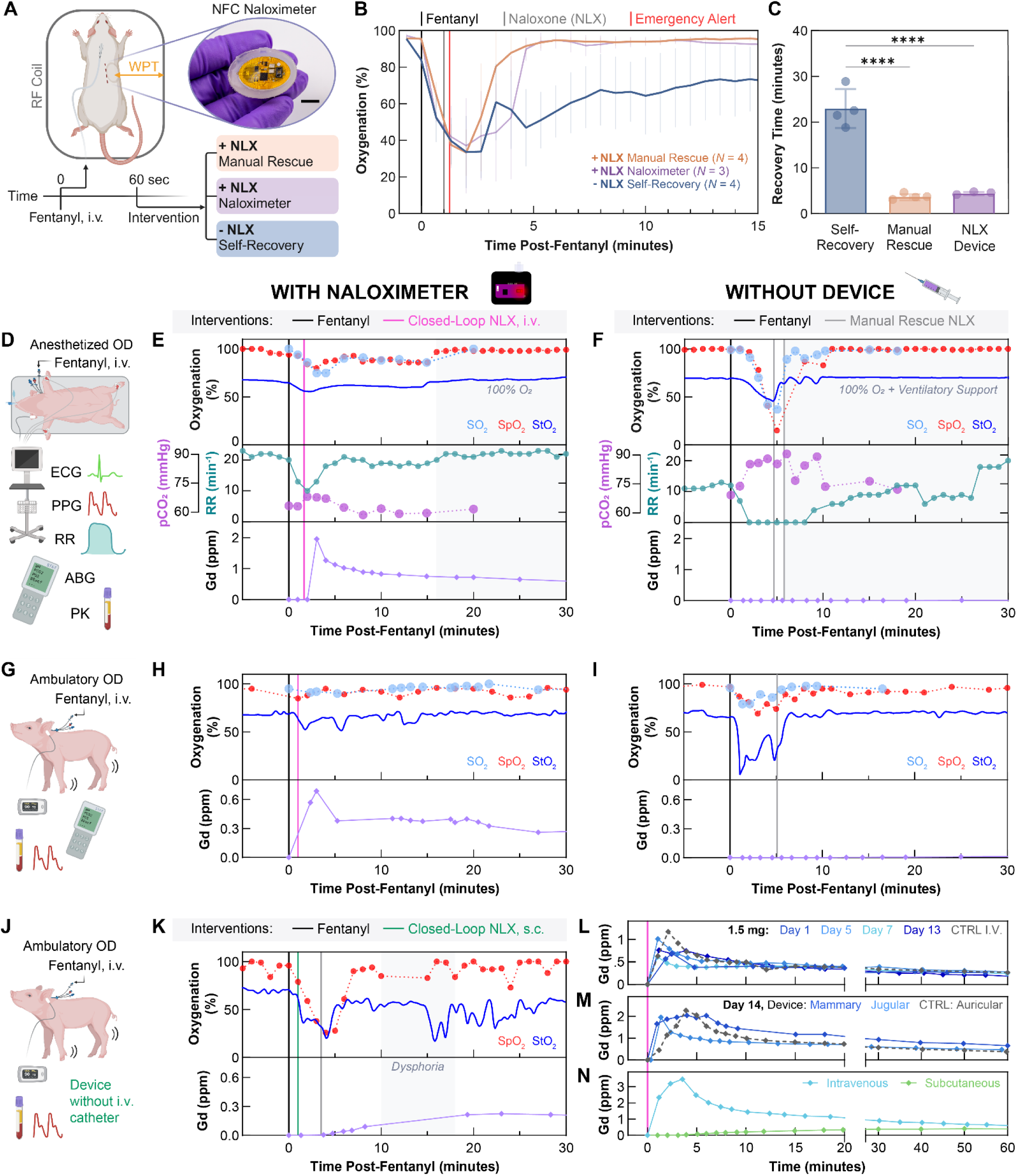
Demonstration of autonomous rescue from life-threatening opioid overdose in rodents and freely moving porcine models. (**A**) Subcutaneous implant location for a miniaturized, battery-free wireless NFC Naloximeter designed for use in rodents and corresponding fentanyl overdose experiment grouping by naloxone (NLX) treatments, *inset:* optical image of the device, scale bar is 1 cm. (**B**) Oxygenation in a rodent model following fentanyl overdose and manual subcutaneous administration of NLX (manual rescue), autonomous rescue enabled by administration of NLX with a Naloximeter, or self-recovery without administration of NLX. Error bars depict SD at each point. Sample sizes are in the legend. (**C**) Comparison of recovery time between self-recovery and NLX treatments, *P*<0.0001 for manual and closed-loop rescue. Data points correspond to individual animals. The bars depict the mean and the error bars are SD. (**D**) Experimental setup for demonstrations of overdose (OD) rescue in an anesthetized pig model (**E**) with and (**F**) without a Naloximeter, the latter of which required manually administered NLX to rescue. Oxygenation (top), respiratory vitals (center), and pharmacokinetic data (bottom) for each fentanyl overdose. (**G**) Experimental setup for demonstrations of OD rescue in ambulatory pigs (**H**) with and (**I**) without a Naloximeter. (**J**) Demonstration of subcutaneous drug delivery from (**K**) a Naloximeter without venous access in a freely moving pig, insufficient to recover from OD, requiring manual administration of NLX to rescue. Panels (**H**, **I**, and **K**) show oxygenation (top) and pharmacokinetic (bottom) data for each experiment. The oxygenation in **E**, **F**, **H**, **I** and **K** was measured with blood gases (SO_2_), a commercial pulse oximeter (SpO_2_), and the Naloximeter (StO_2_). (**L**) Longitudinal pharmacokinetic studies with an intravenous Naloximeter at 1 to 13 days post-implantation, and comparison to delivery of NLX with a continuous infusion pump as a control (CTRL I.V.). Dose is 1.5 mL. (**M**) Gadolinium pharmacokinetics when delivered via a Naloximeter (Device) at 14 days post-implantation in the jugular or mammary vein, or continuous infusion pump (CTRL) in the auricular vein. Dose is 2 mL. (**N**) Gadolinium pharmacokinetic profile when delivered by a Naloximeter intravenously (jugular vein) or subcutaneously. Dose is 3 mL.

Large animal rescue is more complex because, as with humans, the response to fentanyl varies between individuals. Survival studies in pigs simulate real-world uses of the intravenous Naloximeter for humans. Details of the surgical approach, experiment timeline, and device biointegration are in the methods section and Fig. S20 and S30 - S32. Initial experiments validate the closed-loop rescue capability using animals anesthetized with the hypnotic agent propofol and monitored using clinical systems (Fig. 4D). In one example, the pig was administered a dose of fentanyl (2.5 µg/kg) after a post-operative healing period of 14 days. The onset of respiratory depression occurred within 1 min, causing oxygen desaturation and hypercapnia observed by the optical sensor in the Naloximeter (StO_2_) and clinical systems (SO_2_ and SpO_2_) between 1-1.5 min (Fig. 4E, top and center). The Naloximeter triggered closed-loop drug delivery at 1.67 min after fentanyl administration and Gd (NLX proxy) appeared in the bloodstream by 3 min (Fig. 4E, bottom). The animal recovered to baseline respiratory rate within 5 min of closed-loop rescue, and oxygenation remained unchanged until the inspired oxygen returns to 100% at 15 min after administration. A control experiment involved delivery of the same dose to the same animal without activation of the Naloximeter. As before, respiratory depression onset occurred at 1 min, but without the Naloximeter rescue, apnea occurred at 2 min, with significant hypoxia and hypercapnia (Fig. 4F) until the manual administration of two doses of NLX and rescue breaths. Delivery of excess inspired oxygen (100%) and mechanical ventilation supported the animal to recovery. Additional Naloximeter rescue demonstrations (*N* = 4 total) and further vital signs are in Fig. S33 - S35.

Tests of fentanyl overdose in conscious, freely moving pigs validated the ODA and rescue capabilities of the Naloximeter for eventual applications with humans (Fig. 4G). In this case, the ODA detected the overdose at 55 s after fentanyl administration (30 µg/kg), and triggered the rescue cascade at 65 s. The initial desaturation event was short-lived due to the reversal effect of NLX, and the pig recovered to baseline levels of oxygenation within 10 min of the overdose (Fig. 4H, top). Pharmacokinetic data show the NLX proxy in the bloodstream within 1 min of pump activation (Fig. 4H, bottom). The control case led to a severe desaturation event with signs of possible reoxygenation at 3 min followed by an immediate sharp desaturation before imminent death averted by manual rescue (Fig. 4I). An important additional observation is that drug delivery with the same electrolytic pumping Naloximeter performed without the catheter (Fig. 4J), is insufficient to rescue from fentanyl overdose (30 µg/kg), as illustrated by a deep desaturation event (Fig. 4K, top) that is similar to the control case. Specifically, despite closed-loop triggering of the rescue cascade at 1 min, pharmacokinetic data shows very low levels of drug in the bloodstream even after 5 min in this delivery modality (Fig. 4K, bottom). These findings likely result from frustrated diffusion of NLX through fibrotic tissue that forms around the device. Intravenous Naloximeters with catheters implanted for several weeks in both peripheral and central vasculature show similar pharmacokinetic profiles to the control case (infusion pump, Fig. 4L and 4M). When compared directly to subcutaneous drug delivery without the catheter (Fig. 4N), the rapid delivery of NLX to the bloodstream is clearly critical to overdose rescue. Additional pharmacokinetic studies (*N* = 11) in Fig. S36 substantiate this trend.

## Discussion

This work introduces three Naloximeter device platforms for autonomous rescue from opioid overdose, enabling fundamental studies in rodent models and providing two options suitable for clinical translation to humans. Survival experiments in large and small animals demonstrate the capabilities for automatically detecting and quickly responding to overdose through the combined use of a dual-wavelength optical sensor and a drug delivery system. In general, we observe that overdoses treated with the Naloximeter are much less severe, with minimal lasting effects to overall health. Rapid delivery of NLX to the bloodstream is key to rescue, a design feature in both Naloximeter embodiments for clinical translation.

The envisioned patient population includes those in recovery from OUD, due to their high risk of fatal overdose, where device implantation can occur just before their release from treatment. Knock-on effects of non-fatal overdose are a part of OUD that is often overlooked but contribute significantly to the morbidity observed in those who have survived an overdose (*16*, *17*). For implantation in humans, we envision outpatient procedures that align with traditional subcutaneous pacemakers and vascular access ports, surgeries encountered by millions worldwide (*51–54*). While there is a risk of thrombus and complications with indwelling catheters, we observed no significant difference in a biomarker for cardiac damage between samples drawn pre- and post-Naloximeter activation (*N* = 12, Fig. S37). Relative to the injector embodiment, the intravenous device optimizes speed of NLX to the bloodstream over risk of potential complications. The availability of both technologies provides valuable options for the patient and their physician. Future work will include detailed comparative studies, with specific focus on the injector version.

The simplicity of the optical sensor and associated algorithms and their ability to function reliably in porcine models represent attractive aspects of the engineering approaches described here. Further study of overdose detection and false positive rejection in humans would be essential for clinical translation. Alternative or additional sensors for motion or heart rate may be considered for enhanced robustness, at the expense of engineering complexity. Though demonstrated specifically in the context of opioid overdose, the Naloximeter platform is not limited to this disease paradigm and can be adapted to treat other emergency rescue scenarios with rapid pharmacokinetic demands such as anaphylaxis or epilepsy.

## Materials and Methods

### Description of the Electronics Module

The electronics consisted of two printed circuit boards (PCBs; 1 oz. Cu on FR-4, 0.4 mm), one active and one passive. The controlling PCB (active), 15 × 31 mm^2^, contained all electronic components while a supporting PCB (passive), 34 × 42 mm^2^ for the intravenous device and 31 × 45 mm^2^ for the injector device, contained the planar coil for wireless recharging and interdigitated electrodes for electrolytic gas generation (intravenous device). Electroless plating (ENIG) formed a conformal gold layer (75 nm) on the Cu interdigitated electrodes. The controlling electronics comprised a low power Arm Cortex M3 microcontroller unit (MCU, CC2640R2F, 5 × 5 mm^2^, Texas Instruments) that supports Bluetooth low energy (BLE) 5.1 communication protocols. This MCU controlled all aspects of device operation and wireless communication. An integrated optical sensor (MAX30101, Maxim Integrated Products, Inc.) acted as a dual-wavelength optical sensor equipped with light emitting diodes (LEDs, red = 660 nm and near-infrared = 880 nm). This component interfaced with the MCU for recording of photocurrent from the embedded silicon photodetector (PD) at 50 samples per second and 16-bit resolution. Two general purpose input/output (GPIO) pins in the MCU served as logical controls for actuation of drug delivery. In the intravenous device configuration, an n-MOSFET served as a switch for electrolytic gas generation. In the injector configuration, an H-bridge driver chip (DRV8837C, Texas Instruments) provided power to the DC motor in both forward and backward directions. Magnetic induction was used to wirelessly charge a 75 mAh lithium polymer (LiPo) battery. During wireless charging, a transmitter antenna operating at the industrial, scientific, and medical (ISM) band of 13.56 MHz, coupled with the planar coil on the supporting PCB, enabled wireless power transfer (WPT). The induced alternating voltage was rectified and regulated to 5 V to power the battery management chip (LTC4065EDC, Analog Devices Inc.) at a charging current of ∼30 mA. The system design also included a remote reset mechanism for the MCU in the event of a failure in the process of starting the device. A negative temperature coefficient (NTC) sensor interfaced with the MCU to provide measurements of temperature. Figure S1 and S2 show an electronic circuit diagram and functional block diagram of the device, respectively. Heat generated by the optical sensor and device electronics during operation is negligible, as measured by an infrared camera (FLIR A655sc, Teledyne FLIR) shown in Fig. S5. Wireless recharging of devices generated minimal thermal load, as measured by the NTC on the device, and plotted in Fig. S6B and S6C for the implanted and non-implanted cases, respectively. The temperature of devices during charging (T_max_ = 41.7°C) is below the threshold for burn injury, which can be 43°C for up to 8 hours (*55*).

### Fabrication of Naloximeters for Porcine Model Studies

The fabrication of the implantable Naloximeter devices consisted of three parts: electronics module, drug delivery pump (two types), and encapsulation.

#### Electronics Module

The designs of the PCBs made use of a commercial computer aided design utility (Autodesk Fusion 360, Autodesk Inc.) and an outsourcing vendor (PCBWay, Inc.) for fabrication. Low-temperature soldering bonded the electronic components to the PCBs. Half-cut castellated holes served as a means for mounting the controlling PCB to the supporting PCB. After connecting a battery to the supporting PCB, marine epoxy (Marine Epoxy, Loctite) covered the exposed soldering connections.

#### Drug Delivery Pump – Intravenous

Pump assembly began with the fabrication of flexible membranes. The block copolymer polystyrene-*b*-polyisoprene-*b-*polystyrene at 22 wt. % styrene (SIS22; Sigma Aldrich) was dispersed in toluene (≥ 99.5% purity, Sigma Aldrich) at 30 wt. % concentration using a combination of ultrasonication and stirring at 40 °C. Spin-coating this solution on a silicon wafer treated with a hydrophobic coating (Rain-X Inc.) formed the membrane film, thickness: ∼70 μm. Membranes remained at room temperature in a fume hood for at least 1 hour following spin-coating to ensure evaporation of the volatile solvent.

Figure S14 illustrates the methods for assembling the intravenous pump. Briefly, a stereolithographic (SLA) 3D printer (Form 3B, Formlabs Inc.) fabricated the device housing (drug reservoir) and electrolyte reservoir in a biocompatible resin (BioMed Clear V1, Formlabs). After removal of printing supports, sanding and sonication produced finished parts with low surface roughness. Waterproof polyurethane sealant (3M Marine Adhesive Sealant Fast-Cure 5200, 3M Inc.) bonded the electrolyte reservoir to the membrane. After curing, the membrane/electrolyte reservoir was released from the substrate and inserted into the device housing with the same sealant to form the drug reservoir with a water-tight bond. A male Luer lock fitting (716281, QOSINA Corp.) was secured within the device housing with superglue (Gel Control superglue, Loctite) and additional waterproof sealant. After inserting the battery into the device chamber, the supporting PCB containing the electrode bonded to the electrolyte reservoir and device housing with additional waterproof sealant.

#### Drug Delivery Pump – Injector

Fabrication of pumps for the injector device began with SLA 3D printing of a two-piece housing, plungers, and flexible bridges in the same biocompatible resin described above. The drug reservoir consisted of 50 μm-thick ethylene vinyl alcohol (EVOH) film (Non-Catch, Kyodoprinting) manually cut and heat-sealed to the desired shapes. Bonding the EVOH drug reservoir and Huber needle (22G, SAI Infusion) with waterproof polyurethane sealant and marine epoxy completed the fabrication of the needle plunger structure. Biomedical epoxy (EA M-31CL, Loctite) secured the plungers, a 316 stainless steel nut, and four flexible bridges together to form a plunger sub-assembly. After inserting the battery, motor, and the plunger sub-assembly into the lower half of the device housing, waterproof polyurethane sealant bonded the supporting PCB to the lower portion of the device housing. Additional waterproof sealant secured the upper device housing with the lower portion, to complete the pump assembly for the injector device.

#### Encapsulation

Figure S15 illustrates the encapsulation process, beginning with the controlling PCB in the electronics module. A liquid-phase adhesion-promoter treatment used 3- (trimethoxysilyl)propyl methacrylate (Millipore Sigma) in a 1:50:50 volumetric mixture with isopropanol and deionized water, and was applied immediately before depositing a conformal coating of parylene-C (10-13 μm, dichloro [2,2] paracyclophane, SCS Labcoater II, Specialty Coating Systems Inc.). Thereafter, the controlling PCB was soldered to the supporting PCB and assembled pump. Biomedical epoxy then covered both PCBs as a secondary biofluid barrier. A molded encapsulation layer of poly(dimethylsiloxane) (PDMS; Sylgard 184, Dow Corning) provided a soft and biocompatible tissue interface. In the intravenous device, a filling port (0.6 mm diameter) was drilled at the electrolyte reservoir level after encapsulation. An aqueous solution of potassium hydroxide (50 mM, Fisher Chemical) served as the electrolyte and the filling port was sealed with fast-curing silicone adhesive (Kwik-Sil, World Precision Instruments). In the case of the injector device, tape sealed an outlet hole for the needle, prior to forming the PDMS encapsulation. Once cured, a biopsy punch removed the tape and PDMS at the outlet to expose the interior of the chamber. A polytetrafluoroethylene (PTFE)/silicone sealing septum (Sigma Aldrich), secured with Kwik-Sil, covered the opened outlet. The dimensions of the intravenous and injector devices were 46 × 36 × 16.5 mm^3^ and 55 × 46.5 × 18.5 mm^3^, respectively.

### Fabrication of Miniaturized Naloximeters for Rodent Model Studies

Fabrication of the battery-free electrolytic pumps designed for use in rodents followed the same procedures as described above, but on a smaller scale. Two-sided flexible PCBs (f-PCBs, PCBWay, Inc) supported interdigitated gold-plated electrodes and electronic components on opposite sides. Low temperature solder bonded surface-mount components to the bottom side of the f-PCB. The top side (containing electrodes) was covered with a laser-cut release liner to protect the active metal surface during the deposition of parylene-C (10-13 μm) on the f-PCB, as described above. After removing the release liner, the f-PCB was attached to the electrolyte reservoirs using a layer of waterproof polyurethane sealant. The devices were encapsulated in a similar molding process as above, with a layer of PDMS over the electronics, and an overcoat of a soft, biocompatible silicone (Ecoflex 00-30, Smooth-On, Inc.) to form the upper aspect of the encapsulation. The drug outlet at the top of the device was sealed prior to encapsulation and reopened after de-molding, at which point a microfluidic channel layer (Ecoflex 00-30) was secured at the outlet with fast-curing silicone (Ecoflex 00-35 FAST, Smooth-On, Inc.). The circuit and logic diagrams and exploded view of the device are shown in Fig. S28 and S29.

### Studies of Release Rate from the Intravenous Device

A 10 mM aqueous solution of Rhodamine B (Acros Organics) filled the drug reservoir of the electrolytic pump. Deionized water (70-90 g) was added to a beaker with a magnetic stir bar and placed onto a stirring plate. A Groshong catheter (7 French) was filled with water from the beaker and connected to the device, with the free end of the catheter secured in the beaker of water. The electrolytic pump was activated via command from the mobile application and aliquots from the cup were collected every 30-60 seconds for 10 minutes. A benchtop plate reader (Synergy NEO2, BioTek Instruments, Inc.) analyzed the aliquots in duplicate by measuring the absorbance spectrum between 450 and 650 nm. A calibration curve with reference concentration samples was included in each plate. The absorbance value at 553 nm was used to calculate the time-resolved dye concentration in the beaker (accounting for volume loss due to aliquot sampling) based on the Beer-Lambert Law. The volume of dye released by the pump was then computed based on a simple dilution factor.

### Calculation of Tissue Oxygenation (StO_2_)

A differential spectroscopy method estimated the regional tissue oxygenation (StO_2_) using the two LED colors (*λ*_1_= 660 nm and *λ*_2_= 880 nm) and the integrated silicon PD of the biometric optical sensor (MAX30101, Maxim Integrated). This method used the modified Beer-Lambert law of optical absorption in scattering media (*56–58*) to calculate changes in oxygenated (HbO_2_) and deoxygenated (Hb) hemoglobin. At *λ*_1_= 660 nm the molar absorption coefficients are ε_Hb-λ1_ = 344.2 mm^-1^M^-1^ and ε_HbO2-λ1_ = 44.5 mm^-1^M^-1^; whereas for *λ*_2_= 880 nm they are ε_Hb-λ2_ = 83.57 mm^-1^M^-1^ and ε_HbO2-λ2_ = 119.9 mm^-1^M^-1^ (*50*). Solving the linear equation, Eq. 1, resulting from the differential optical losses at the two wavelengths yielded the changes in the concentration of Hb (Δ*cHb*) and HbO_2_ (Δ*cHbO*_2_) with respect to baseline.

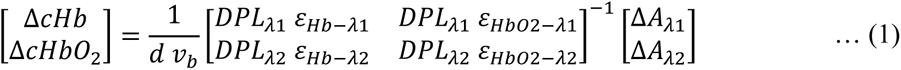

Here, *d* is the LED-PD separation distance (3.25 mm), *v*_*b*_ is the blood volume fraction of the tissue, *DPL*_*λ*1,2_are the differential path lengths at the two wavelengths (*λ*_1_and *λ*_2_). The time-dependent difference in the optical signal (Δ*A*_*λ*1,2_) with respect to baseline, measured at the two wavelengths, is Δ*A*_*λ*1,2_(*t*) = − log _10_(*I*_*λ*1,2_(*t*)⁄*I*_*λ*1,2_(*t*_0_)), where *I*_*λ*1,2_(*t*) is the digitalized photocurrent data at time *t* and *I*_*λ*1,2_(*t*_0_) is the photocurrent recorded at baseline (*t*_0_). The ratio of oxygenated hemoglobin (*cHbO*_2_) to total hemoglobin (*cHbO*_2_ + *cHb*) on a concentration basis defines the tissue oxygenation *StO*_2_(*t*), Eq. 2.

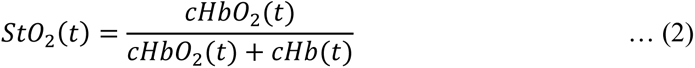

Where *cHbO*_2_(*t*) = *cHbO*_2_(*t*_0_) + Δ*cHbO*_2_(*t*), and *cHb*(*t*) = *cHb*(*t*_0_) + Δ*cHb*(*t*). In porcine muscular tissue, *v*_*b*_ = 1.0%, *DPL*_*λ*1_ = 9.65 and *DPL*_*λ*2_ = 9.09 (*59*, *60*). The baseline *StO*_2_ is defined as 70%, and the concentration of total hemoglobin is 130 g/L (2 mM), which corresponds to baseline *cHb*(*t*_0_) = 0.6 mM and *cHbO*_2_ (*t*_0_) = 1.4 mM.

### Large Animal Studies

All large animal studies were performed according to the protocol approved by the Northwestern University Institutional Animal Care and Use Committee (study #00018098). Female Yorkshire Landrace crossbred pigs (27 – 30 kg) were obtained from Oak Hill Genetics (Ewing, IL, USA) and examined by veterinary staff upon arrival. Pigs were housed in groups of two to three animals on a 14:10-hour light:dark (LD) cycle (lights on at 6:00 AM) during an acclimation period of at least 5 days prior to any surgical intervention. Following device implantation surgery, pigs were single-housed and remained on the same 12:12 LD cycle. The temperature in the facility ranged from 21 to 24 °C, while the humidity was maintained between 30 and 70%. Pigs were fed a standard diet of pig feed twice daily throughout experiments, which were conducted during the light cycle. Pigs were fasted for 12 hours preceding any surgery or experimental procedures, water was available ad libitum. Figure S30 shows the general experiment timeline.

#### Device Implantation Procedure

Ethylene oxide gas sterilized the closed-loop devices in a standard 24-hour cycle (Anprolene AN74, Andersen Sterilizers). Utilizing aseptic technique, the devices were filled with drug solution containing a 9:1 volumetric mixture of naloxone (NLX; Naloxone HCl Injection 1 mg/mL, International Medication Systems) and gadolinium (Gd) contrast agent (ProHance, gadoteridol injection 279.3 mg/mL, Bracco Diagnostics, Inc.) immediately prior to implantation. A sterile one-way duckbill valve (11582, QOSINA Corp.) was attached to the device once filled.

Pigs were sedated and prepared for surgery with telazol (6 mg/kg), propofol (0.5 – 1.5 mg/kg), and atropine (0.05 mg/kg) and intubated. Isoflurane anesthetic (1 – 5%) was used to effect, throughout all surgical procedures. Carprofen (4 mg/kg) and bupivacaine (4 – 10 mL) served as pre-operative and local analgesics, respectively. Antibiotic cefazolin (22 mg/kg) and anticoagulant heparin (100 mg/kg) were administered intra-operatively. Figure S20 illustrates the implantation procedure for devices in the jugular vein. Incision sites were shaved and scrubbed three times with alternating betadine and ethanol swabs to disinfect. For devices in the jugular vein, a 3 – 5 cm incision was made in the ventral neck and blunt dissected to expose the external jugular vein. An 8 – 10 cm incision bisecting the brachiocephalic muscle was made along the dorsal aspect of the neck, and blunt dissection produced a subcutaneous pocket for the device. A trocar created a small subcutaneous tunnel between the device pocket and the vascular access incision, and the intravenous catheter (5 – 8 French) was pulled through. The catheter was measured and cut to a length which ensured 7 – 10 cm of catheter in the vessel and minimal slack between the device and vascular access. A blunt needle or similar Luer connector supplied by the manufacturer connected the device to the free end of the catheter which had been locked with a solution of dextrose/heparin (500 IU/mL sodium heparin, 25 wt.% dextrose) to maintain patency (*61*). At this point, the device was inserted into the subcutaneous pocket with the PCB side facing the muscular tissue, and the catheter protruded from the neck incision. The catheter was introduced to the vessel and secured with finger-trap sutures (3-0 silk). A suture (4-0 silk) through the fixation wing on the device secured it to the underlying muscular tissue. The subcutaneous space was closed with absorbable tacking sutures (3-0 PDS, polydioxanone). Finally, the skin was closed in layers with absorbable sutures (3-0 PDS) in the subcutaneous layer and barbed absorbable sutures (2-0 Quill) in the dermis.

A similar method was employed in the implantation of intravenous devices in the mammary vein, with an incision just above the nipple line for venous access and a subcutaneous device pocket along the upper aspect of the iliocostalis. Figures S31 and S32 show examples of healed devices implanted in the jugular and mammary veins, respectively.

Pigs were administered carprofen (2 mg/kg) for two days post-operatively, and antibiotics (16 mg/kg, Clavamox), aspirin (325 mg tablet), and clopidogrel (75 mg tablet, Plavix) for the duration of the study. Pigs recovered for at least 2 days following device implantation before additional surgical procedures, i.e. catheterization. When necessary, devices were recharged transdermally (Fig. S6) with a custom wireless charger consisting of a transmitting coil (diameter 10 – 12 cm, 2 turns) connected to a radio frequency power supply (13.56 MHz, 10 W, Neurolux, Inc.).

#### Catheterization Procedure (Porcine)

Pre-operative medication, sedation, and aseptic procedures were identical to that used in device implantation surgery. A standard central venous catheterization approach was employed, in which the pig is positioned in dorsal recumbency and a midline incision (2 – 4 cm) is made in the ventral neck. Blunt dissection and isolation of the carotid artery and jugular vein followed by insertion of catheters (8 – 9.5 French), secured with finger trap sutures (3-0 silk) established vascular access. Subcutaneous tunneling with a trocar to the dorsal aspect of the neck created exit sites for both catheters. Catheter patency was confirmed, and the indwelling lines were locked with heparinized saline before closure. The midline incision was closed in layers with absorbable sutures (3-0 PDS) in the subcutaneous layer and barbed absorbable sutures (2-0 Quill) in the dermis. Additional finger-trap sutures secured the lines to the skin at their exit site and a veterinary jacket (MPS-Top Shirt Dog, Medical Pet Shirts) worn by the animal minimized the risk of complications from the indwelling lines.

#### Fentanyl Overdose Experiments (Porcine)

In anesthetized studies, pigs were sedated with telazol (6 mg/kg) and atropine (0.05 mg/kg) and intubated. A propofol infusion of 6 – 12 mg/kg/hour maintained light anesthesia. Physiological monitoring devices were placed to monitor heart rate (electrocardiography), SpO_2_ and pulse (PPG), and respiratory vitals (capnography) with a veterinary vitals machine (LifeWindow 6000V, Digicare Biomedical). A handheld blood gas analyzer (i-STAT 1, CG8+ cartridges, Abbott Laboratories) provided on-site measurements of blood gases (ABG) from fresh arterial samples. Samples for small molecule pharmacokinetic (PK) analysis were collected in EDTA plasma tubes (K_2_EDTA Vacuette, Greiner Bio-One), stored on ice, and centrifuged within one hour of collection.

The pig was breathing spontaneously throughout the duration of the study, and the inhalational gas mixture was titrated from 100% oxygen to 100% medical air in the 30 minutes prior to fentanyl administration. Baseline data was recorded with the implanted devices and physiological monitoring devices. A baseline blood sample for ABG and PK was drawn within the 5 minutes preceding fentanyl administration (fentanyl citrate, intravenous, 2.5 – 10 µg/kg) which occurred at time = 0. Blood samples for PK were collected every 1 minute for the first 10 minutes, every 2 – 3 minutes for the following 10 minutes, and every 10 minutes thereafter until time = 60 minutes. Blood samples for ABGs were collected at the same frequency as PK samples until recovery (approximately 20 minutes), limited by the running time of the instrument.

In ambulatory experiments, pigs with implanted devices were contained in a large-animal transport cart equipped with a squeeze panel. A portable clip-on oximetry device (Vetcorder Pro, Sentier Connect) accomplished external monitoring of SpO_2_ and pulse. Baseline and blood draw procedures remained the same as anesthetized experiments. Fentanyl (30 – 100 µg/kg) was administered at time = 0. Food was withheld until after recovery from overdose.

Upon detecting overdose, the devices autonomously triggered NLX release. In the “control” case, devices were configured to record StO_2_ but not armed to trigger the rescue operation. Animals were observed for as long as possible until imminent veterinary intervention was required, at which point rescue dose(s) of intravenous NLX were administered. Each rescue dose was 1 mg (1 mg/mL Naloxone HCl solution). Additional lifesaving measures were supplied as indicated.

#### Longitudinal Pharmacokinetic Studies (Porcine)

Pigs were sedated with telazol (6 mg/kg) and atropine (0.05 mg/kg) and intubated. Inhalational isoflurane (1 – 3%) maintained adequate anesthetic depth. Physiological monitoring devices were placed to monitor heart rate (electrocardiography), SpO_2_ and pulse (PPG), and respiratory vitals (capnography) with a veterinary vitals machine (LifeWindow 6000V, Digicare Biomedical). Samples for small molecule PK analysis were collected in EDTA plasma tubes (K_2_EDTA Vacuette, Greiner Bio-One), stored on ice, and centrifuged (3500 rpm, 8 minutes) within one hour of collection.

A baseline blood sample for PK was drawn within the 5 minutes preceding NLX administration, which occurred at time = 0 via manual pump activation with the mobile application. Blood samples for PK were collected every 1 minute for the first 10 minutes, every 2 – 3 minutes for the following 10 minutes, and every 10 minutes thereafter until time = 60 minutes. In “control” intravenous studies, a continuous infusion pump (BeneFusion SP3 Vet, Mindray Animal Care) supplied the same NLX/Gd mixture used in devices at a flow rate of 30 mL/hr. In “control” injection studies, manual subcutaneous injection of the NLX/Gd mixture occurred at time = 0.

#### Freely Moving Oximetry (Porcine)

A device connected with the mobile application was placed outside of the home cage of a pig with implanted devices. A command from the mobile application started oximetry recording. The implanted device streamed continuously until the recording was stopped or the battery was drained on either the implanted or cellular device. Normal feeding and animal care occurred without disruption during the recording period.

#### Hypoxia Challenge Experiments (Porcine)

The hypoxia procedure was approved by the Institutional Animal Care and Use Committee of Washington University in St. Louis. Female Yorkshire Landrace crossbred pigs (45 – 55 kg) were obtained from Oak Hill Genetics (Ewing, IL, USA). The pigs were sedated, intubated, and brought under general anesthesia with isoflurane (1 – 5%). Carotid arterial and femoral venous lines placed via cutdown allowed for blood sampling. Devices were implanted subcutaneously in the foreleg (deltoid and triceps) and medial rectus abdominis, and StO_2_ recording commenced. Arterial and venous blood gases (ABG and VBG) were measured from time-synchronized blood samples and analyzed with a blood gas analyzer (StatPrime CCS Analyzer, Nova Biomedical). A commercial ear-mounted probe measured SpO_2_ while a skin-mounted commercial probe system (T.Ox, Vioptix) measured cutaneous StO_2_ to compare with device recordings.

Baseline data was collected at normoxic (21% O_2_) conditions, followed by periods of hypoxia induced by modulation of the inspired oxygen fraction (balance: nitrogen). Blood samples for ABG/VBG were drawn many times across the spectrum of inhaled gas mixtures. Oxygen saturation (SO_2_) in capillary blood was defined by a weighted average of venous and arterial SO_2_ (70V:30A) and compared to StO_2_ measured by the implanted device. This experimental approach was partially based on validation studies for an FDA-approved tissue oximetry device (*62*). The hypoxia challenge occurred in 2 animals, with one animal containing 2 devices and the other utilizing 1. Resulting comparisons between Naloximeter StO_2_ and the other oximetry measurements are plotted in Fig. S17 and S18.

#### Cardiac Biomarker Analysis (Porcine)

Samples for cardiac biomarker analysis were collected before and after intravenous Naloximeter activation. Troponin I was quantified with a canine/feline assay (Troponin I, IDEXX Laboratories, Westbrook, Maine USA). Samples were collected in serum clot activator tubes (CAT Serum Clot Activator Vacuette Tubes, Greiner Bio-One). Pre-activation samples were collected at the time of baseline PK blood collection and post-activation samples were collected between 60 – 65 minutes after pump activation. Blood samples were incubated at room temperature for 30 minutes before centrifugation (3000 rpm, 10 minutes), then stored at -4°C and shipped same-day to the analytical lab. The results of this analysis are in Fig. S37.

### Small Animal Studies

All procedures were approved by the Institutional Animal Care and Use Committee of Washington University in St. Louis. Male and female Sprague Dawley rats (250 – 350g) were used for all rodent experiments. Rats were initially group-housed with two to three animals per cage on a 12:12-hour LD cycle (lights on at 7:00 AM) and acclimated to the animal facility holding rooms for at least 7 days. The temperature in the holding rooms ranged from 21 to 24°C, while the humidity was maintained between 30% and 70%. Rats received food and water ad libitum throughout experiments, which were conducted during the light cycle.

#### Surgical Procedures

Animals were deeply anesthetized using isoflurane (5% induction, 2% maintenance) and administered carprofen (5 mg/kg, SC), enrofloxacin (8 mg/kg, SC, Baytril), and bupivacaine (5 mg/kg, intravenous catheterization) or lidocaine (8 mg/kg, device implantation) for pre-and intra-operative analgesia. Clippers shaved the surgical site and 3 applications of betadine/ethanol swab sterilized the area. A recirculating heated water blanket warmed animals during surgery.

#### Intravenous Catheterization (Rodent)

A small incision on the ventral surface of the neck provided access to the jugular vein, wherein an indwelling catheter was inserted and sutured in place with non-absorbable (6-0 silk) suture. The catheter was tunneled subcutaneously to the dorsal neck and exited via another small incision. The ventral incision in the muscle was closed with absorbable (4-0 vicryl) suture and the skin incision was closed using non-absorbable (6-0 silk) suture. The exposed catheter was connected to a backpack device (Vascular Access Harness, Instech) containing a port for drug administration. Rats received carprofen tablets (Rimadyl 2 mg, MD150-2, Bio-Serv) for two days after surgery to assist in wound healing and analgesia. A sterile gentamicin/saline solution (0.3 mL at 1.33 mg/mL gentamicin) administered daily maintained catheter patency. Rats were single-housed following surgery and allowed to recover for one week prior to device implantation surgery.

#### Device Implantation (Rodent)

Ethylene oxide exposure (24-hour cycle) served as the method for device sterilization. A five-centimeter incision was made on the back between the shoulder blade and spine to expose the latissimus dorsi and external oblique muscles. Blunt dissection created a subcutaneous pocket sized appropriately to the device. Aseptic technique was used to fill the devices with a prepared NLX/saline solution (6.67 mg/mL naloxone hydrochloride dihydrate; pharmaceutical grade, Millipore Sigma) immediately before implantation, and Kwik-Sil sealed the filling port. The device was inserted into the pocket and a running stitch of absorbable suture (5-0 PDS) was used to make tacking sutures to close the excess subcutaneous space. The skin incision was closed using absorbable (5-0 PDS) subcuticular sutures and topical tissue adhesive (GLUture, World Precision Instruments) was applied to the incision. Betadine and topical lidocaine ointment were applied over the closed incision for their antiseptic and analgesic properties, respectively. Post-operatively, rats were treated with anti-inflammatory carprofen (5 mg/kg) and antibiotic enrofloxacin (8 mg/kg, Baytril) once daily for up to seven days.

#### Hypoxia Challenge Experiments (Rodent)

Hypoxia-induced changes in pulse oxygenation (SpO_2_) and tissue oxygenation (StO_2_) were examined in catheterized rats with implanted devices. A gas blender (Hypoxydial; Starr Life Sciences) regulated the mixture of inhaled gases (O_2_ and N_2_) for precise oxygen concentration. Rats were lightly anesthetized with isoflurane (3% induction, 2% maintenance) and fitted with a pulse oximeter collar (MouseOx v2.0, Starr Life Sciences) to measure SpO_2_ while the implanted device measured StO_2_. Baseline data was collected at normoxic (21% O_2_) conditions, followed by stepwise adjustment in 5-minute episodes down to a minimum of 8% O_2_. Additional datasets used in the correlation of SpO_2_ and StO_2_ (Fig. 2H) are included in Fig. S16.

#### Fentanyl Overdose Experiments (Rodent)

Rats with implanted devices and catheters were lightly anesthetized with isoflurane (5% induction, 1 – 1.5% maintenance) and fitted with a pulse oximeter collar (MouseOx v2.0, Starr Life Sciences). Cardiorespiratory parameters (oxygen saturation, heart rate and respiratory rate) were collected throughout the experiment using the collar oximeter and the implanted device. All animals received an intravenous administration of fentanyl citrate (20 µg/kg) at time = 0. After 60 seconds, animals in the manual injection treatment group received a subcutaneous injection of NLX (Naloxone HCl, 1 mg/kg at 6.67 mg/mL concentration; Millipore Sigma). Animals in the closed-loop treatment group had implanted devices which automatically triggered NLX release according to the overdose detection mechanism (see “Automated Rescue Implementation”). Animals in the self-recovery group did not receive any intervention.

### Naloxone (NLX) and Fentanyl Pharmacokinetic Quantification

Plasma specimens were analyzed using a modification of a validated liquid chromatography tandem mass spectrometry (LC-MS/MS) assay (*63*). Briefly, 100 µL of plasma was transferred to a conical tube with lid (1.5 mL tube, Eppendorf) and 100 μL of methanol containing the internal standards (10 ng/mL) was added. Samples were vortexed for 3 min and centrifuged for 10 min (26,000 g, 4 °C). Addition of 170 µL of sample to an HPLC vial containing 800 µL of water formed the analyzed solution. An LC-MS/MS tandem mass spectrometer (ABSciex 5500, SCIEX) with a turbo V ion source operated in positive electrospray ionization (ESI) mode quantified fentanyl and NLX in the plasma. A liquid chromatography system (1200 Series, Agilent Technologies) equipped with a quaternary pump, a temperature-controlled column compartment, and an autosampler (HTC PAL autosampler, Leap Technologies) performed the chromatography. The column (2.6 μm, 3.0 x 100 mm Kinetex F5 core-shell column, Phenomenex) separated the analytes using an HPLC flow rate of 1 mL/min with two mobile phases, A: 0.2% aqueous formic acid and B: acetonitrile (LC-MS grade). The column temperature was 50 °C. Limits of quantifications were 0.5 ng/mL for fentanyl and NLX. Upper limit of quantification was 1000 ng/mL for fentanyl and 250 ng/mL for NLX.

### Gadolinium (Gd) Pharmacokinetic Quantification

PK data collected with the Gd tracer element were analyzed with inductively coupled plasma mass spectrometry (ICP-MS). Pre-weighed metal-free tubes (MetalFree™ 15 mL, Labcon) collected 200 – 500 μL fresh blood per sample and were kept frozen at -20 °C until post-weighed and workup began. Blood samples were treated with 0.5 mL trace grade nitric acid (> 69%, Thermo Fisher Scientific) and 0.5 mL trace grade hydrogen peroxide (> 30%, GFS Chemicals) and placed at 65 °C for at least 3 hours to allow for complete sample digestion. Ultrapure water (18.2 MΩ·cm) was then added to produce a final solution of 5.0% nitric acid (v/v) in a total volume of 10 mL.

Dilution of a quantitative standard (1,000 μg/mL Gd elemental standard, Inorganic Ventures) to 500 ng/g element concentration in 5.0% nitric acid (v/v) formed the quantitative standard. A subsequent 100x dilution created a second quantitative standard of 5 ng/g Gd in 5.0% nitric acid (v/v). A solution of 5.0% nitric acid (v/v) was used as the calibration blank.

ICP-MS was performed on a computer-controlled (QTEGRA software) instrument (iCapQ, Thermo Fisher Scientific) operating in STD mode and equipped with an autosampler (ESI SC-2DX PrepFAST, Elemental Scientific, Inc.). Internal standard was added inline using the prepFAST system and consisted of 1 ng/mL of a mixed element solution containing Li, Sc, Y, In, Tb, Bi (IV-ICPMS-71D, Inorganic Ventures). The prepFAST system also carried out inline dilutions to generate a calibration curve consisting of 500, 100, 50, 25, 10, 5, 1, 0.5, 0.25, 0.1, and 0.05 ppb Gd. Each sample was acquired using 1 survey run (10 sweeps) and 3 main (peak jumping) runs (40 sweeps). The isotopes selected for analysis were ^156,157^Gd, and ^115^In, ^159^Tb (chosen as internal standards for data interpolation and machine stability). Instrument performance is optimized daily through autotuning followed by verification via a performance report (passing manufacturer specifications).

### Statistical Analysis

Animals were randomly assigned to treatment groups. Statistical analyses were performed using Prism (GraphPad Software). Sample sizes are reported as number of animals (*N*) or trials/devices (*n*) in the figure captions. All data are expressed as mean ± SD or as individual plots. One-way analysis of variance (ANOVA) was used for multiple group comparisons, and *P* values less than 0.05 were considered significant. The before-after comparison in Fig. S37 is a paired, two-sided (conservative) t-test. Raw data (*λ*_1_and *λ*_2_) is plotted without smoothing or adulteration. FFT spectra from the raw optical signals were computed with MATLAB (Mathworks, Inc.).

## Acknowledgments

We thank the members of the Rogers laboratory for useful discussions of the project, particularly S. Madhvapathy for advice on several aspects of device survivability. We also thank E. Dempsey, I. Stepien, N. Haack, and C. Haney for advice related to small animal experiments; H. Saleh, M. McIntyre, K. Dunlap, and C. Jones for assistance with large animal studies; and M. Park for preliminary efforts in device development. This research was supported by the National Institutes of Health through the NIH HEAL Initiative (https://heal.nih.gov/) under award numbers UG3DA050303 and UH3DA050303. J.L.C. graciously acknowledges support from the National Science Foundation Graduate Research Fellowship under Grant No. DGE-2234667. Swine studies were assisted by the Large Animal Surgical team in the Northwestern University (NU) Center for Comparative Medicine. This work made use of the NUFAB facility at the NUANCE Center, which has received support from the ShyNE Resource (NSF ECCS-2025633), the IIN, and Northwestern’s MRSEC program (NSF DMR-2308691). Metal analysis for gadolinium PK studies was performed at the Quantitative Bio-element Imaging Center (QBIC, NU) generously supported by the NIH under grant S10OD020118. This work employed the facilities of the High Throughput Analysis Laboratory (HTAL, NU) and the Center for Clinical Pharmacology at the University of Health Science and Pharmacy. Figures 1A, 4A, 4D, 4G, and 4J were created with illustrations from Biorender.com.

## Author contributions

Conceptualization: JLC, AVG, JAR, RWG, JAM

Methodology: JLC, AVG, MRT, RAO, AJM, JAR, AVG

Software: AVG, JT

Formal analysis: JLC, AVG, VEB

Investigation: JLC, AVG, VEB, JP, BR, MRT, EAC, ARB, RAS, PG, JK, RA, MS, MAK, BVH, MCM, NM

Resources: MP, YH, JAM, JAR, RWG

Writing - Original Draft: JLC, AVG

Writing - Review & Editing: JLC, AVG, JP, PG, VEB, BR, RWG, JAR

Visualization: JLC, AVG, JP

Supervision and Funding Acquisition: JAR, RWG

## Data and materials availability

All data are available in the main text or the supplementary materials.

## Supplementary Text

### Wireless BLE Integration with Implanted Device

The implanted device established BLE wireless communication with a cellular device that runs a custom mobile application shown in Fig. S3. The user can send device configuration parameters (optical sensor: sampling rate and optical intensity; pump: pulse width modulation parameters for actuation) or action commands (start/stop oximetry recording, reset device, and manual pump activation with direction control for the needle injector) to the device using the mobile application graphical user interface. The implanted device transmitted two packets of data every one second. The first packet contained one-second worth of raw photocurrent values and the second contained device status indicators (temperature, pump activation status, battery voltage and wireless charging status). Current consumption during various modes of operation was characterized using a power profiler (Power Profiler Kit II, Nordic Semiconductor) and is reported in Fig. S4 and Table S1.

The mobile application sent a timestamp to the implanted device every 300 seconds, synchronized with respect to the coordinated universal time (UTC), to update the internal real time clock of the MCU. Thus, the mobile application received UTC timestamped data packets and uploaded them to a remote server via internet connection. This data management strategy allowed large-scale data collection from multiple devices and subjects simultaneously.

### Automated Rescue Implementation

Prior to administration of fentanyl, the implanted devices received the “Start Oximetry” command and began data transmission to the cellular device. Upon detecting a drop in oxygenation due to opioid-induced respiratory depression, the mobile application prompted a WARNING, then triggered the RESCUE operation. First, the mobile application sent the device a command to activate drug delivery via BLE. Then, it relayed an emergency call using the cellular network. In the case of animal experiments, the cellular device dialed an emergency contact set up in the mobile application rather than 9-1-1 due to Federal Communications Commission restrictions on non-emergency 9-1-1 calls. In the case of rodent devices, a MATLAB (Mathworks, Inc.) script replaced the mobile application for device control and data collection.

### Theoretical Modeling of Drug Delivery with Electrolytic Pump

We employed an analytical model for electrolytic drug delivery microsystems, derived from singular perturbation methods in our previous work (*64*), to predict the delivery time over a range of geometrical and elasticity parameters in the flexible membrane. The model used a combination of normalized non-dimensional device parameters, including the initial environmental pressure in the body *P*_0_^∗^, initial volume of gas in the electrolyte chamber *V*_0_^∗^, and microfluidic resistance based on the microfluidic layouts *M*^∗^ to characterize the drug delivery process and predict the drug delivery time. For a negligible *V*_0_^∗^, small *M*^∗^, and known initial environmental pressure *P*_0_^∗^, the relationship between the time *t*^∗^and drug volume *V*^∗^ is given by the following non-dimensional parametric expression:

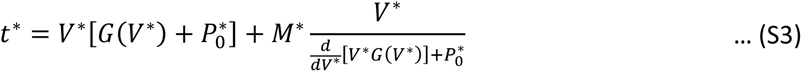

where *G*(*V*^∗^) is the normalized pressure-volume function describing the deformation of the flexible membrane. Equation S1 can be re-written dimensionally as

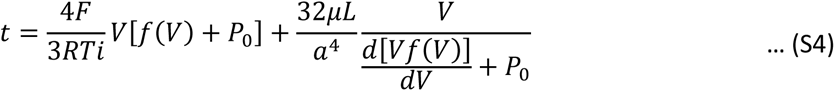

where *F* is Faraday’s constant, *R* is the ideal gas constant, *T* is temperature, *i* is the electrical current applied to the electrodes, *f*(*V*) is the pressure-volume function of the flexible membrane, *μ* is the drug viscosity, *L* is the length of the microfluidic channels, and *a* is the side length of the channel.

The first term in Eq. S2 quantifies the time required for the membrane to overcome the external pressure and deform into a spherical cap and the second term quantifies the time required for the drug to travel through the microchannels. In this work, microchannels are exchanged for an intravenous catheter, which has characteristic length an order of magnitude larger. In this regime, the microfluidic resistance becomes negligible, and instead the “dead volume” (*V*_*DV*_) inside of the catheter tubing becomes significant. Thus, we rewrite Eq. S2 to account for the intravenous device configuration as:

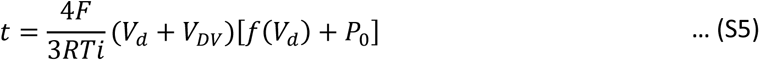

Where the volume delivered to the bloodstream 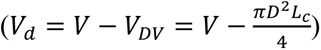 is the difference between the volume of gas generated by electrolysis (*V*) and *V*_*DV*_ for a given the catheter internal diameter (*D*) and length (*L*_*c*_).

The deformation of the membrane from flat to spherical cap (for finite-deformation) is approximated for a linear-elastic material model as

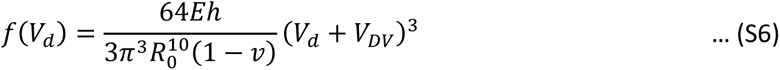

where *E* is the Young’s Modulus, *h* is the thickness, *v* is Poisson’s ratio, and *R*_0_ is the radius of the membrane. A parametric study, shown in Fig. S8, was performed to model the effect of varying the Young’s Modulus (*E* = 1 – 3.5 MPa), thickness (*h* = 40 –150 µm), and radius (*R*_0_ = 10 − 15 mm) of the flexible membrane in the time required to deliver 1000 µL of drug. This was substantiated by experimental evidence tracking the membrane deformation versus time, provided in Fig. S9.

### Preliminary Studies with Cutaneous Naloximeter Sensors for Humans

Cutaneous Naloximeter sensors (oximeters) shared the electronics of the large animal devices but omitted the drug delivery pumps. Two-sided f-PCBs, containing a planar coil for wireless recharging, supported electronic components on one side and a battery on the other. The same firmware, logic, and mobile application was used in both cases. The key components (MCU, LiPo battery, dual-wavelength optical sensor) and fabrication processes were identical with the electronics module of the large animal Naloximeter. Encapsulation of the devices followed a similar PDMS molding process as described above. For measurement sessions, oximeters were secured to the skin of the volar forearm with a skin-safe adhesive (KM 40A, Katecho) and optionally covered with a layer of film dressing (Tegaderm, 3M, Inc.).

Human studies were performed according to the protocol approved by the Northwestern University Institutional Review Board (IRB #: STU00220375). Informed consent was obtained prior to participation. Collection of marked data from an active movement sequence began with baseline data recording for several minutes. The active sequence consisted of a series of movements: finger pressing on the device, running, walking, biking, aerobic exercise (e.g. jumping jacks), and resting (seated and supinated). Start and stop times for each exercise were noted, as well as any observations of physiological condition.

### Overdose Detection Algorithm (ODA)

The overdose detection algorithm (ODA) is founded on the temporal dynamics of physiological signals characteristic to changes in systemic oxygenation. Measurements of the optical absorption of biological tissues allow for quantification of the oxygenated and deoxygenated hemoglobin species, and thus estimation of oxygenation percentage (*65*). Wearable and implantable devices with strategically engineered sensors have demonstrated this optical technique for interrogating biological metrics (*66–69*). While the optical approach is very sensitive to changes in hemoglobin oxygenation *in vivo*, the underlying mathematical model is blind to confounding events produced by motion artifacts or postural changes that disrupt blood flow in tissue surrounding the sensor. These events can lead to calculation of false fluctuations of oxygenation. Furthermore, obstructive sleep apnea represents a confounding affliction that produces real desaturation events of -20% SpO_2_/min lasting ∼25 – 30 s, occurring in a characteristic oscillatory pattern (*70–73*). In a study using cerebral oximetry to determine StO_2_ during apnea-hypopnea events, the StO_2_ desaturation was determined to be -4.02 ± 0.06% in events 12 – 16 s in duration (*73*). Our ODA was designed to discriminate against confounding artifacts that might lead to false overdose detections, such as sleep apnea or motion. The decomposition of the temporal data collected from the dual-wavelength optical sensor permits the extraction of oxygenation level, its temporal changes and motion indices to produce a series of four logical metrics (M1 – M4). The collective multivariable analysis yields a robust set of data analytics that identifies the physiological signatures of an overdose in real-time.

The ODA received packets of data at 1 Hz containing the two raw optical signals, red (*λ*_1_) and infrared (*λ*_2_). The ODA operates at this frequency to perform calculations on the current and preceding nine packets of data. A digital low-pass Butterworth filter (0.1 Hz, 4th order) processed the incoming raw data to calculate the *StO*_2_using Eq. 1 and Eq. 2 and the differential optical signal (Δ*λ* = *λ*_1_ − *λ*_2_). In parallel, a fast Fourier transform (FFT) calculated the power density spectrum on both red and infrared signals filtered using a band-pass Butterworth filter (passband 1-5 Hz, 4th order). The average of the integrated power density spectra provided the power density spectrum index (*PDSi*). Summation of the *PDSi* over the previous 1 and 10 minutes calculated the short-term and long-term motion indices, *STMi* and *LTMi*, respectively. Linear interpolation of *λ*_1_, *StO*_2_, and Δ*λ* followed by differentiation provided the rates of change 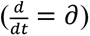 of the red signal (*δλ*_1_), optical difference (*∂*Δ*λ*) and tissue oxygenation (*∂StO*_2_), calculated every 10 seconds over a 60-second window. A comparison of these rates, the *StO*_2_and the *STMi*, against experimentally derived threshold values produced four logical metrics:

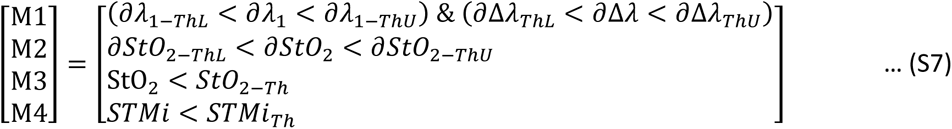

where the subindices in M1 and M2, *ThL* and *ThU*, correspond to the lower and upper thresholds, respectively, and *StO*_2−*Th*_ and *STMi*_*Th*_ to the tissue oxygenation and short-term motion index thresholds, respectively.

An OD-relevant event (EVENT) existed when the mutually inclusive condition of the four metrics in Equation S5 in the ODA was prompted, which evolved into a WARNING after *N* consecutive EVENTs: *N* = 3 if *LTMi* > *LTMi*_*Th*_, corresponding to active/awake state; or *N* = 6 if *LTMi* < *LTMi*_*Th*_, corresponding to rest/sleep state. Upon triggering a WARNING, the ODA prompted the user with a push notification to override the warning, if acknowledged. If one additional EVENT accumulated in the 10 seconds following WARNING without user feedback, the ODA triggered the RESCUE response: activating drug delivery and relaying an emergency call. Figure 3H shows the graphical illustration of the workflow associated with these data analytics.

The following thresholds were used for the metrics:

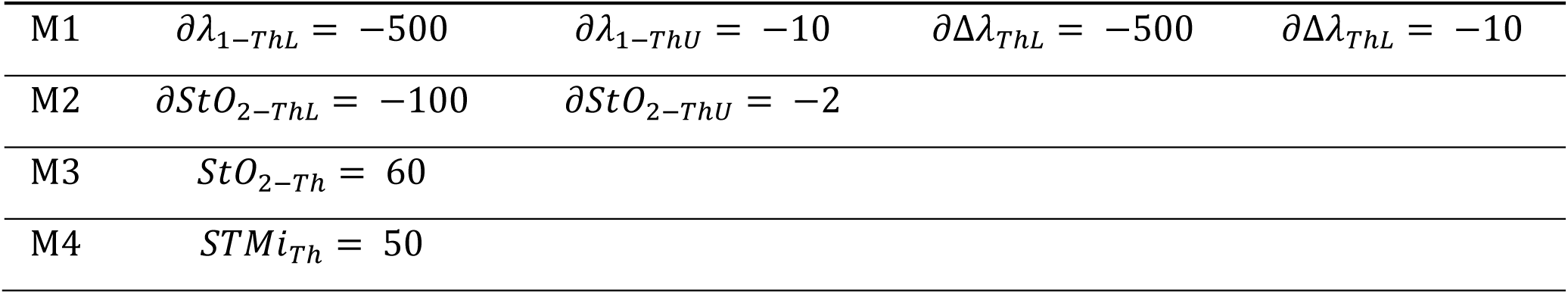

These values were determined as follows. The thresholds for *δλ*_1_, *∂*Δ*λ*, *∂StO*_2_were extracted from data collected during the fentanyl dosage desaturation experiments (e.g. Fig S21 and S25). The 60% threshold for *StO*_2_ corresponds to a physiological level that indicates hypoxia and is equivalent to ∼ 80% SpO_2_ (Fig. 2H) or ∼ 60% SO_2_ (Fig. 2I). Finally, the threshold for the *STMi* is estimated from marked data in human during rest (Fig. S23) and in pigs during rest/sleep (Fig. S24 and S26).

---

**Fig. S1.**
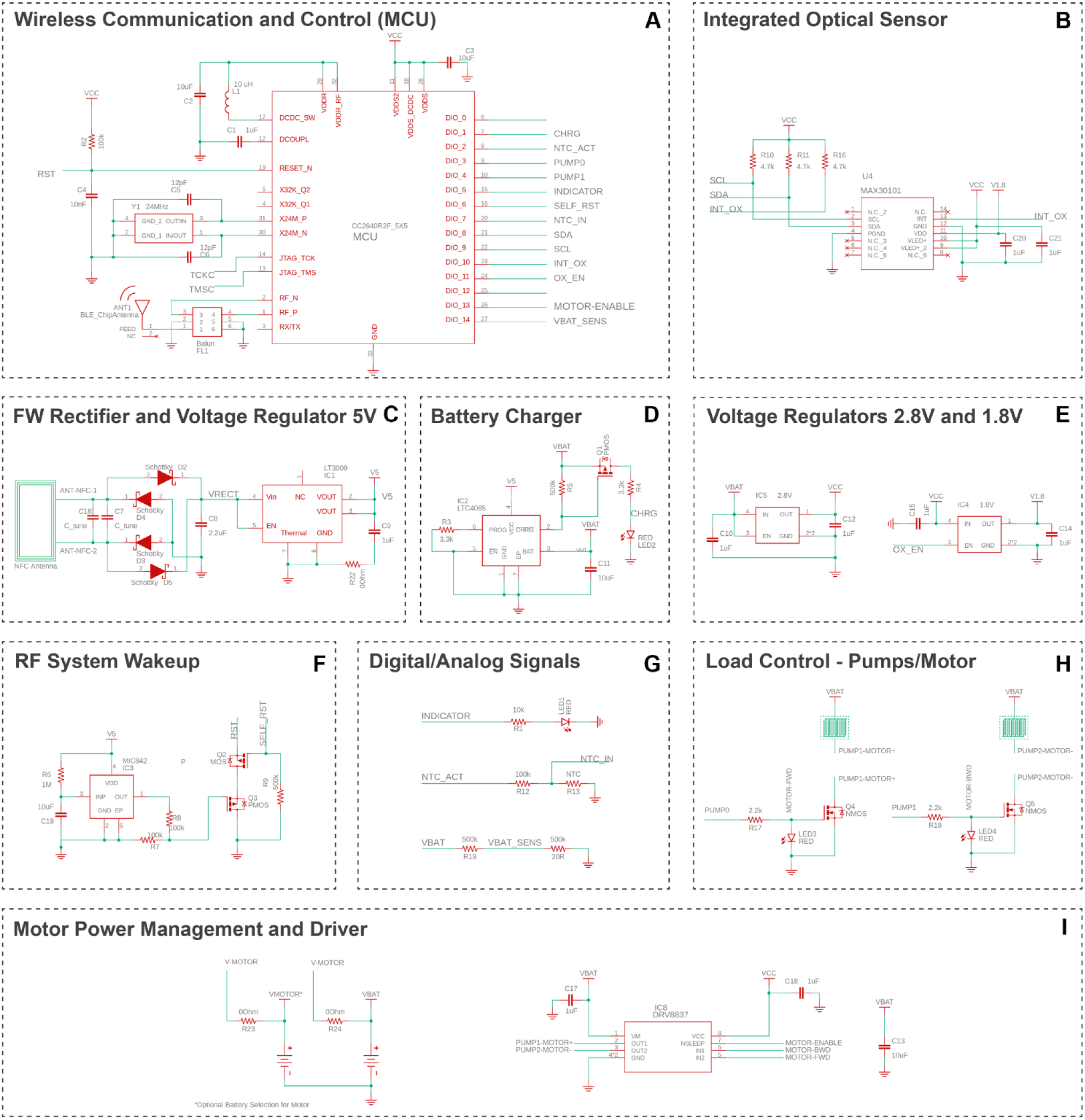
Circuit diagrams for subsystems of the Naloximeter platform for large animals. **(A)** The CC2640R2F microcontroller (MCU) uses BLE wireless communication and controls the operation of the device. **(B)** The integrated optical sensor MAX30101 connects with the MCU using I^2^C serial communication. **(C)** A full-wave (FW) voltage rectifier and regulator converts the harvested input AC to 5 V DC. **(D)** The battery charger is powered by wireless power transfer. **(E)** The step-down voltage regulator supplies 2.8 V to the electronics (left). A 1.8 V regulator provides voltage to the optical sensor only (right). **(F)** An RF actuated wake up system provides a remote reset to the microcontroller in the event of a failed cold start. **(G)** General inputs/outputs control function in the device such as analog battery voltage monitoring, indicator operation, input NTC sensor. **(H)** The power module controls the electrolysis-driven pumps, or the logic signals to operate the H-bridge chip that drives the DC motor. **(I)** The power and control module operates to the DC motor.

**Fig. S2.**
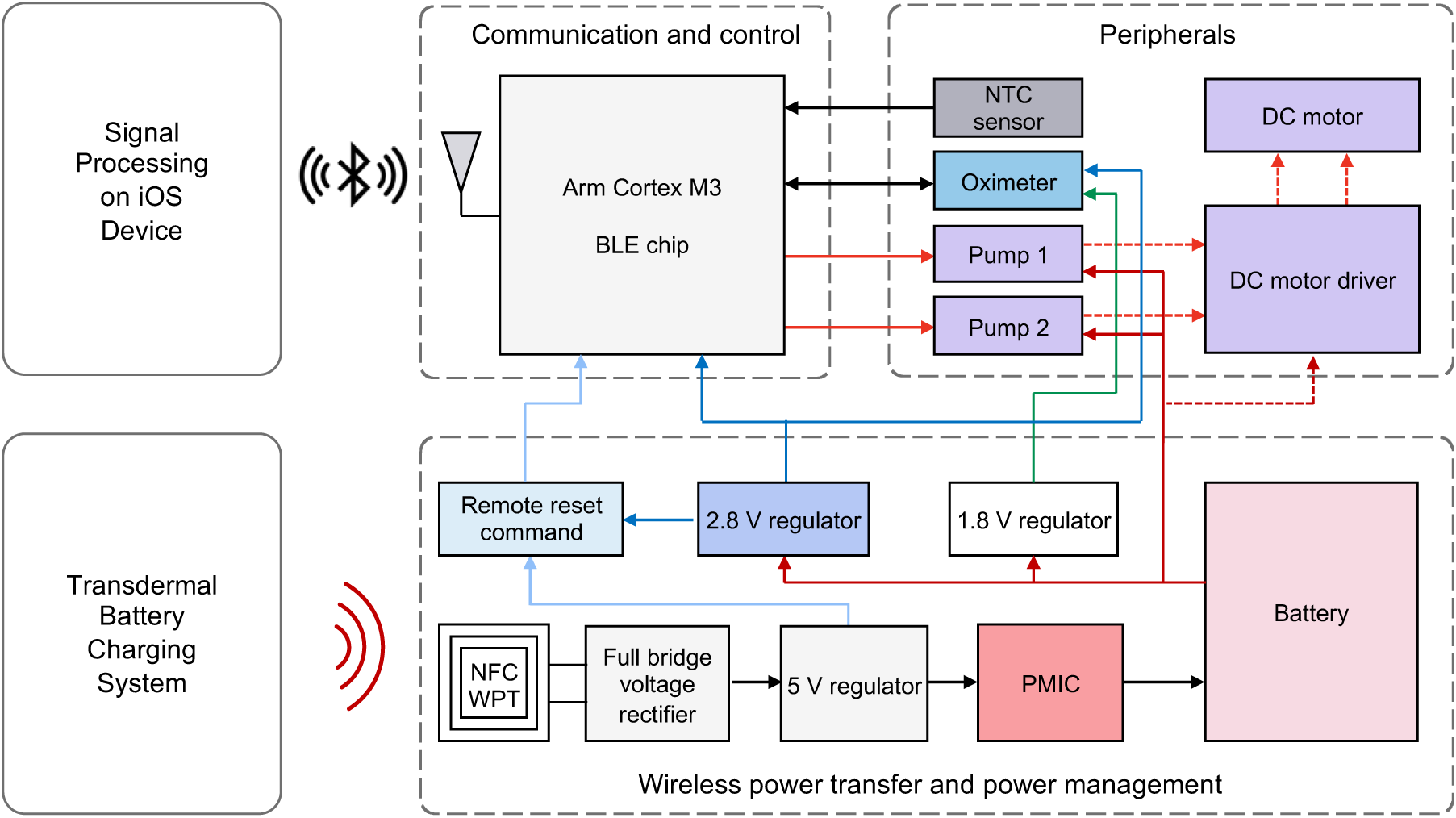
Functional block diagram of the Naloximeter platform for large animals. Modules that construct the electronic system and its interaction with the external iOS device and transdermal battery charging system. The same system, with minor modifications, supports two options for drug delivery, electrolytic pump or DC motor operation.

**Fig. S3.**
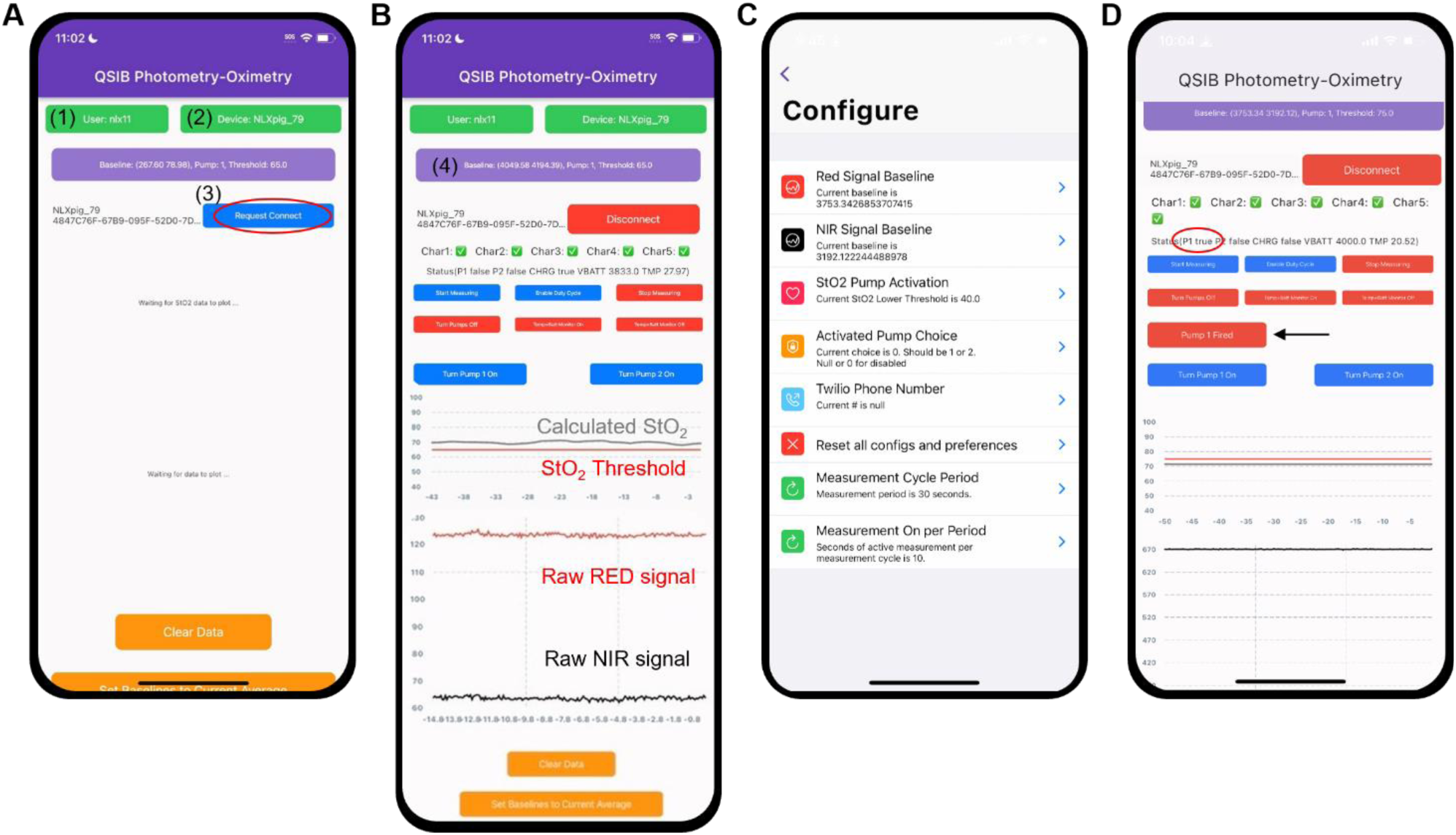
Screenshots of the mobile application on a phone. **(A)** The main app screen indicates (1) cloud account for data management, (2) device selection, and (3) button for establishing a BLE connection to the device. **(B)** View of the main app screen after establishing device connection, with buttons for controlling device operation, including collection of optical sensor data and calculation of StO_2_ in real-time. **(C)** Device configuration window, accessed by clicking on (4). **(D)** Main app screen provides two indicators of NLX deployment: pump (P1) status (red circle) and “Pump 1 Fired” display (black arrow).

**Fig. S4.**
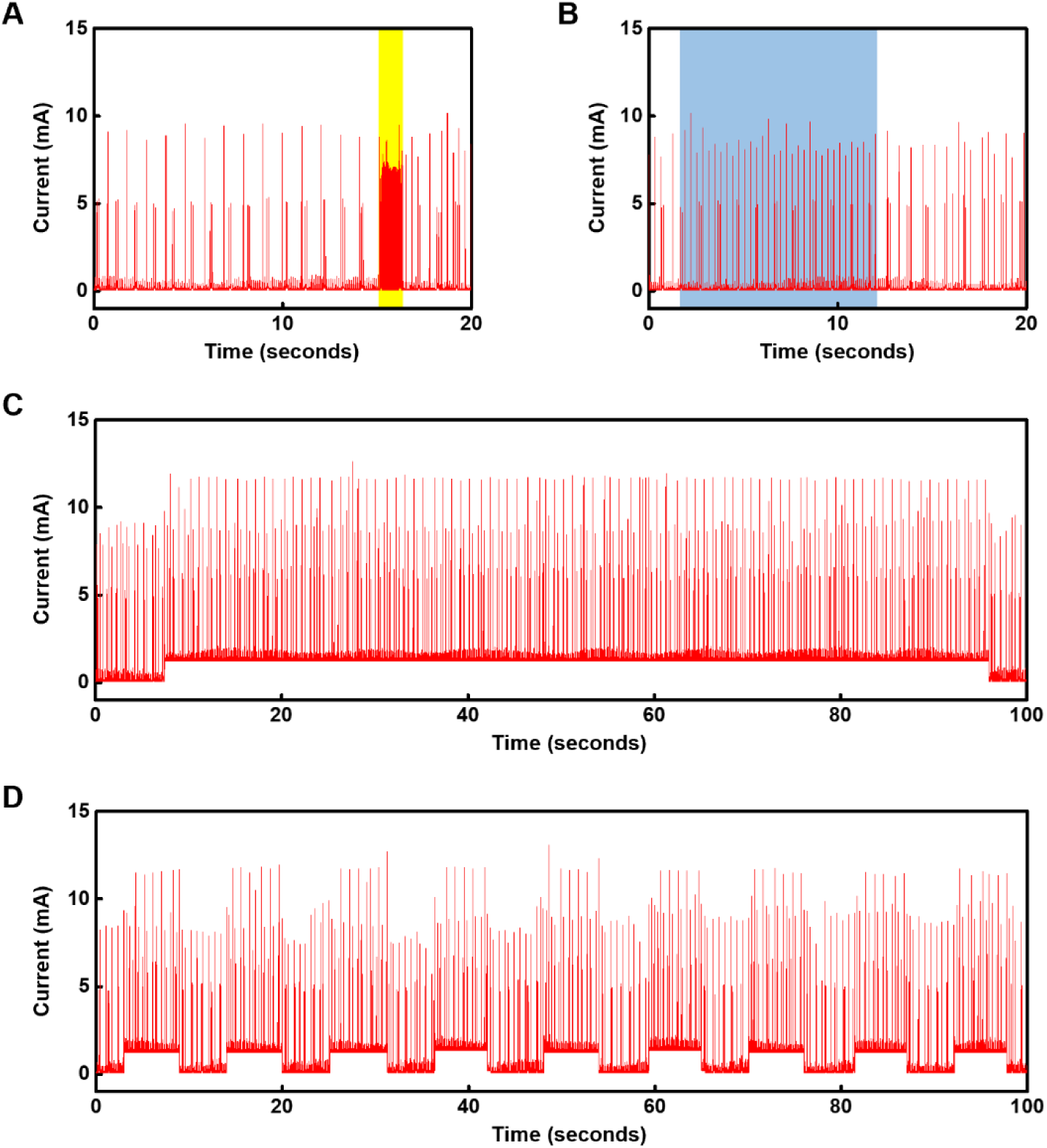
Current consumption during operation of a Naloximeter for large animals. **(A)** Current consumption of the device before, during (yellow area), and after establishing BLE connection to the peripheral device. **(B)** Current consumption of the device while sending status updates (temperature, battery voltage, pump status, and charging status) at intervals of 1 s (blue area) and while idle. **(C)** Current consumption of the device while operating the dual-color optical sensor in normal mode. **(D)** Current consumption of the device while operating the dual-color optical sensor in intermittent mode (50% duty cycle, 10 s period: 5 s ON, 5 s OFF). Supply voltage is 3.8 V in all cases.

**Fig. S5.**
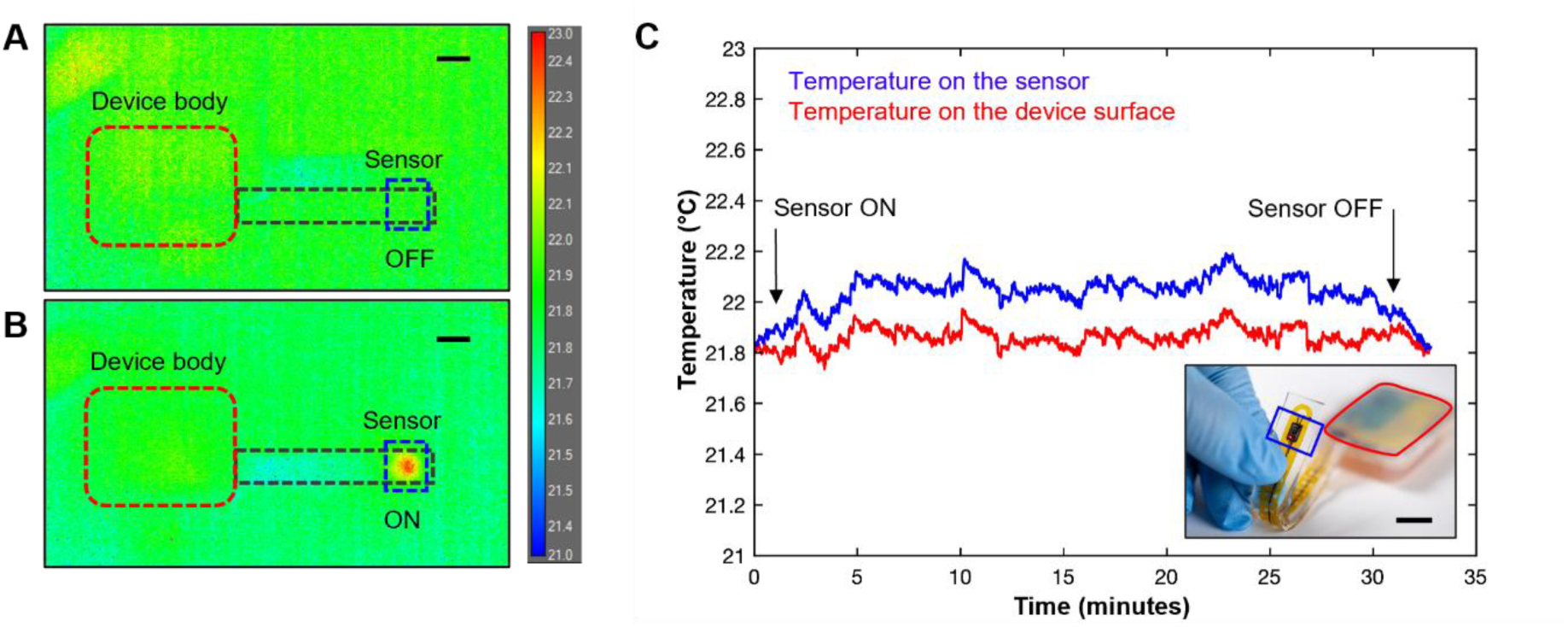
Benchtop characterization of thermal load produced by the dual-wavelength optical sensor during operation. Thermographic images of an encapsulated test of a device with **(A)** the sensor OFF and **(B**) sensor ON at room temperature. Scale bars, 1 cm. **(C)** Surface average temperature recorded in the area above the electronic board and on the sensor. Inset: optical image of the device used in this characterization study, with sensor separated from the body of the device. Scale bar, 1 cm.

**Fig. S6.**
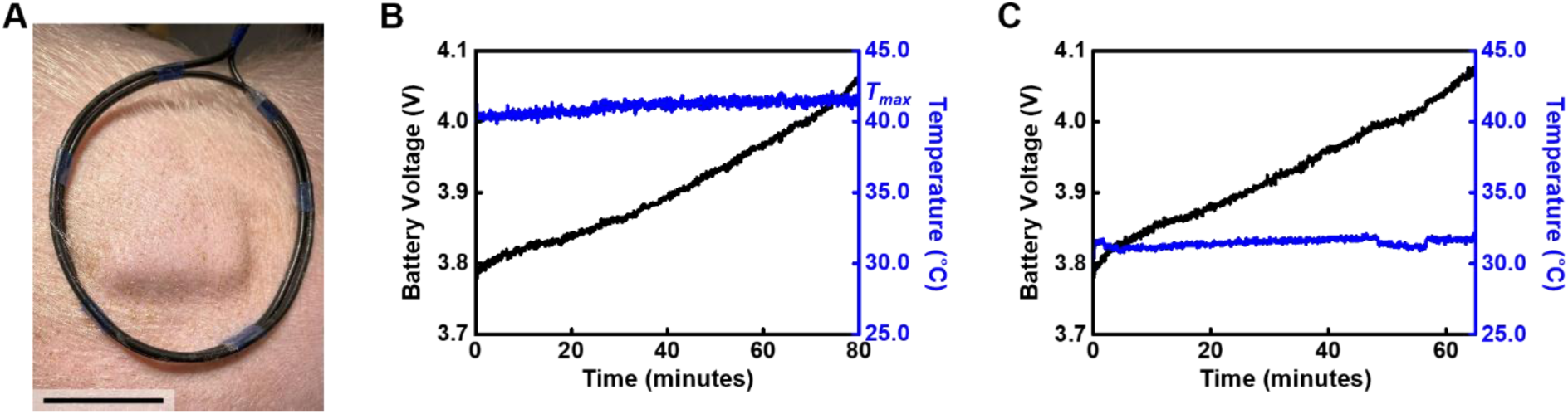
Transdermal charging of a Naloximeter device for large animals. **(A)** Photograph showing wireless transdermal charging of a device implanted in a pig model after healing. Scale bar, 5 cm. **(B)** Battery voltage and temperature measured by the NTC in the device during transdermal charging, maximum temperature labeled as *T_max_*. **(C)** Battery voltage and temperature measured by the NTC in the device during charging on the benchtop.

**Fig. S7.**
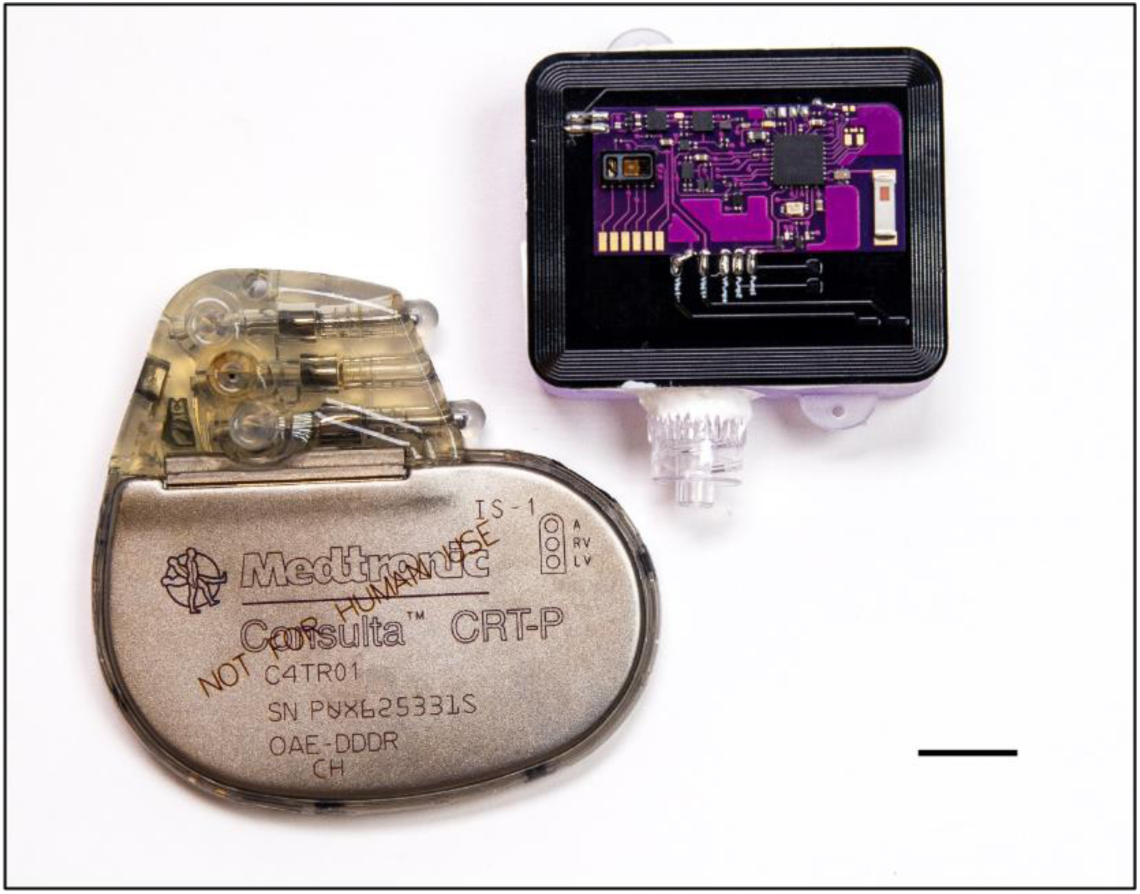
Optical image of a Naloximeter and a commercial pacemaker (Consulta, Medtronic). Scale bar is 1 cm.

**Fig. S8.**
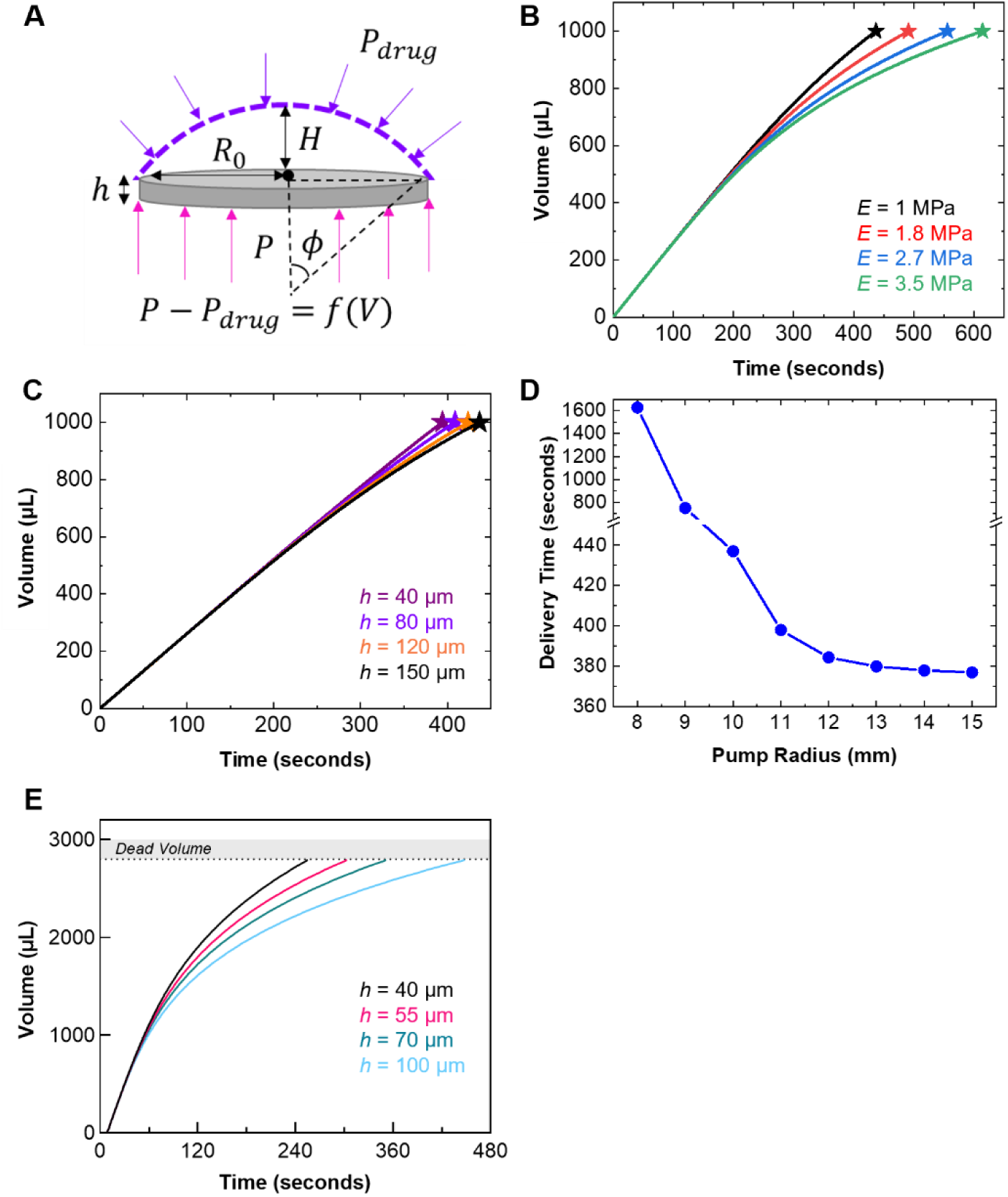
Parametric FEA modeling of the operation of electrolytic pumps. **(A)** Schematic diagram of forces on the membrane relevant to these simulations. **(B)** Volume vs. time curves for 1 mL pumps varying the modulus of elastic membrane (*h* = 1 μm). **(C)** Volume vs. time curves for 1 mL pumps varying the membrane thickness (*E* = 1 MPa). **(D)** Effect of pump radius on delivery time for 1 mL drug volume, (*E* = 4.3 MPa and *h* = 70 μm). Calculations in (A – D) run with constants: *i =* 15 mA, T = 310 K. **(E)** Evaluation of the effects of membrane thickness for 3 mL pumps using parameters measured in actual devices, *i =* 120 mA and *E* = 4.3 MPa.

**Fig. S9.**
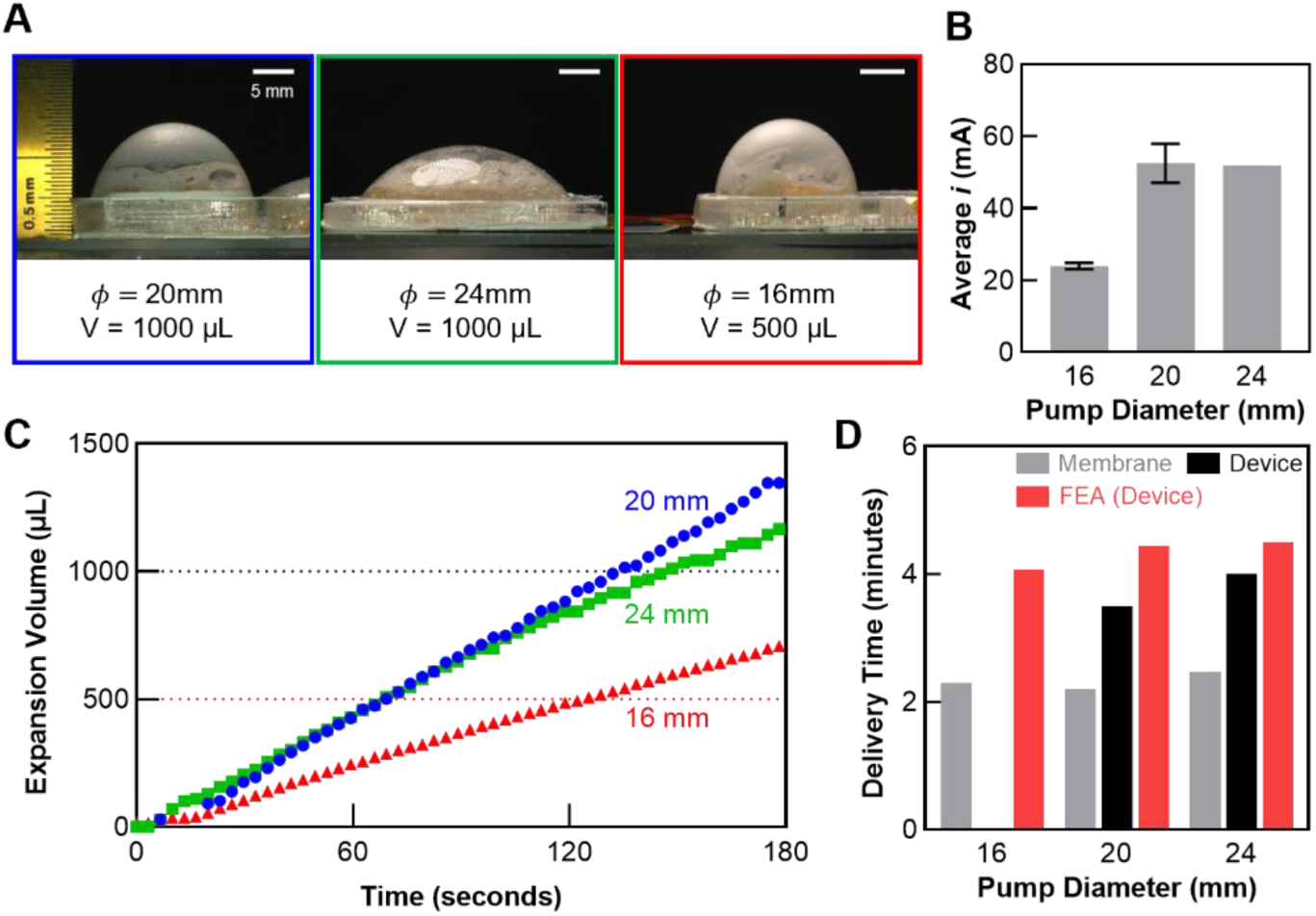
Benchtop studies of the effects of pump geometry on the profile of membrane deformation. **(A)** Photograph of three electrolytic pumps with different geometries at inflated states. Scale bars, 5 mm. **(B)** Average current consumption for pumps with different diameters, recorded during electrolysis with the device PCB and battery. **(C)** Expansion volume calculated by image analysis as the membrane deforms during electrolysis. **(D)** Estimated time to deliver the total drug reservoir content according to benchtop studies of the membrane deformation, full device testing, and FEA modeling *(E* = 4.3 MPa, *h* = 70 μm, *T* = 310 K, and *i* from experiment).

**Fig. S10.**
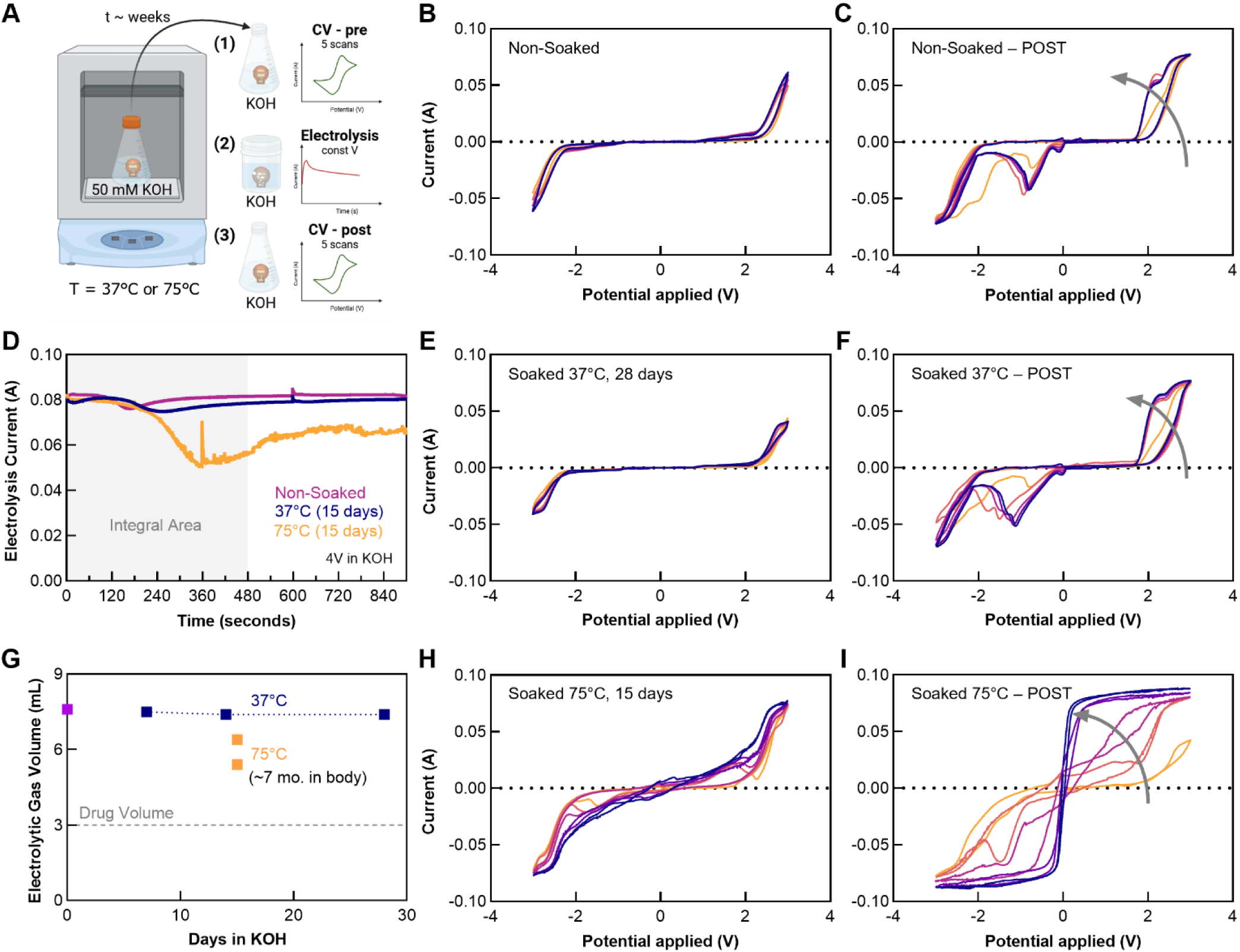
Benchtop studies of electrode degradation. **(A)** Schematic illustration of the experimental setup and study design. The experiments consisted of soaking in potassium hydroxide electrolyte (50 mM KOH) at elevated temperature (37°C or 75°C) for several weeks followed by (*1*) cyclic voltammetry (CV), (*2*) constant voltage electrolysis, and (*3*) post-electrolysis CV. All CV tests involved scans of voltage between +/- 3 V at a rate of 50 mV/sec for 5 cycles. Constant volage electrolysis testing at an applied potential of 4 V. All electrochemical characterization was carried out in the electrolyte solution, 50 mM KOH. Cyclic voltammetry plots for a fresh non-soaked control electrode **(B)** before and **(C)** after electrolysis. (D) Electrolysis current at constant voltage for control and aged electrodes. Cyclic voltammetry plots for an electrode aged at 37°C for 28 days (E) before and **(F)** after electrolysis. **(G)** Volume of gas generated by electrolysis, as calculated with Faraday’s Law and measured current (panel D) plotted according to aging time and temperature. Cyclic voltammetry plots for an electrode aged at 75°C for 15 days – equivalent to 7 months at body temperature, **(H)** before and **(I)** after electrolysis.

**Fig. S11.**
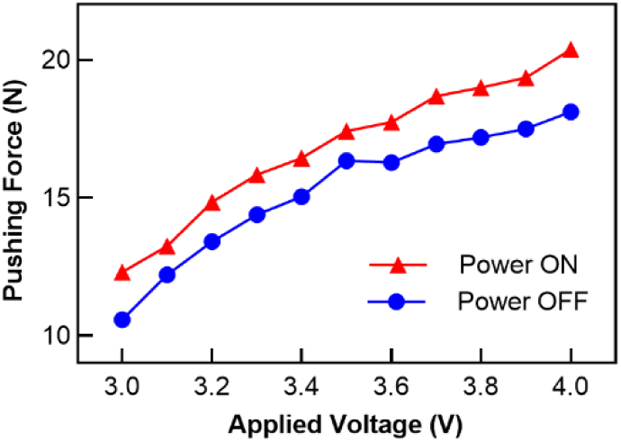
Force supplied by the plunger and DC motor as a function of applied voltage. Each data point corresponds to the saturated force with the maximum load on the motor.

**Fig. S12.**
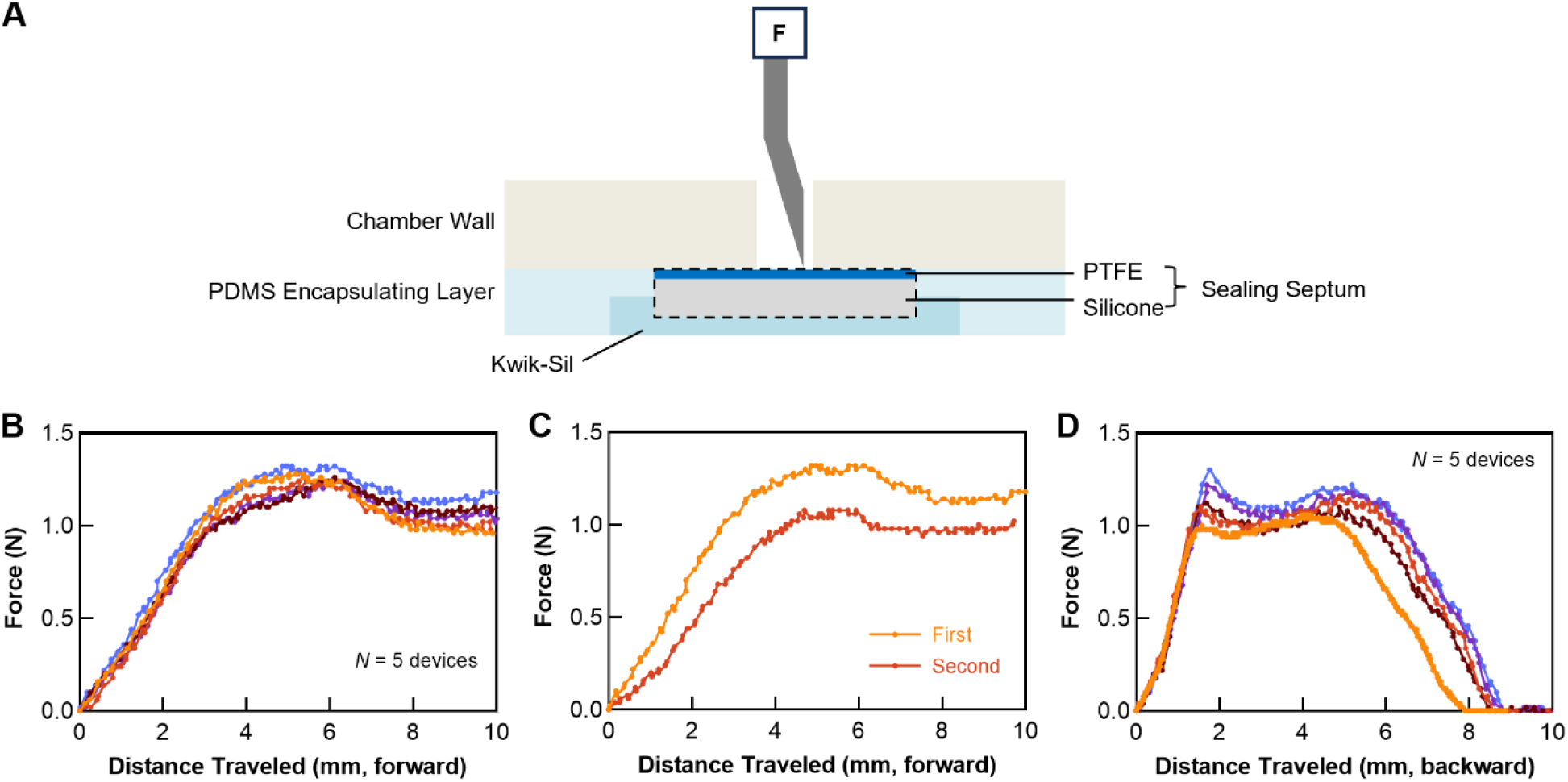
Force to deploy and retract the Huber needle through the sealing septum. **(A)** Schematic illustration of the setup to measure the force to deploy and retract the needle through the sealing septum. **(B)** Required force for the needle to pierce the front sealing septum. **(C)** Required force for the needle to pierce the front sealing septum for the first and second time. **(D)** Required force to retract the deployed needle through the sealing septum.

**Fig. S13.**
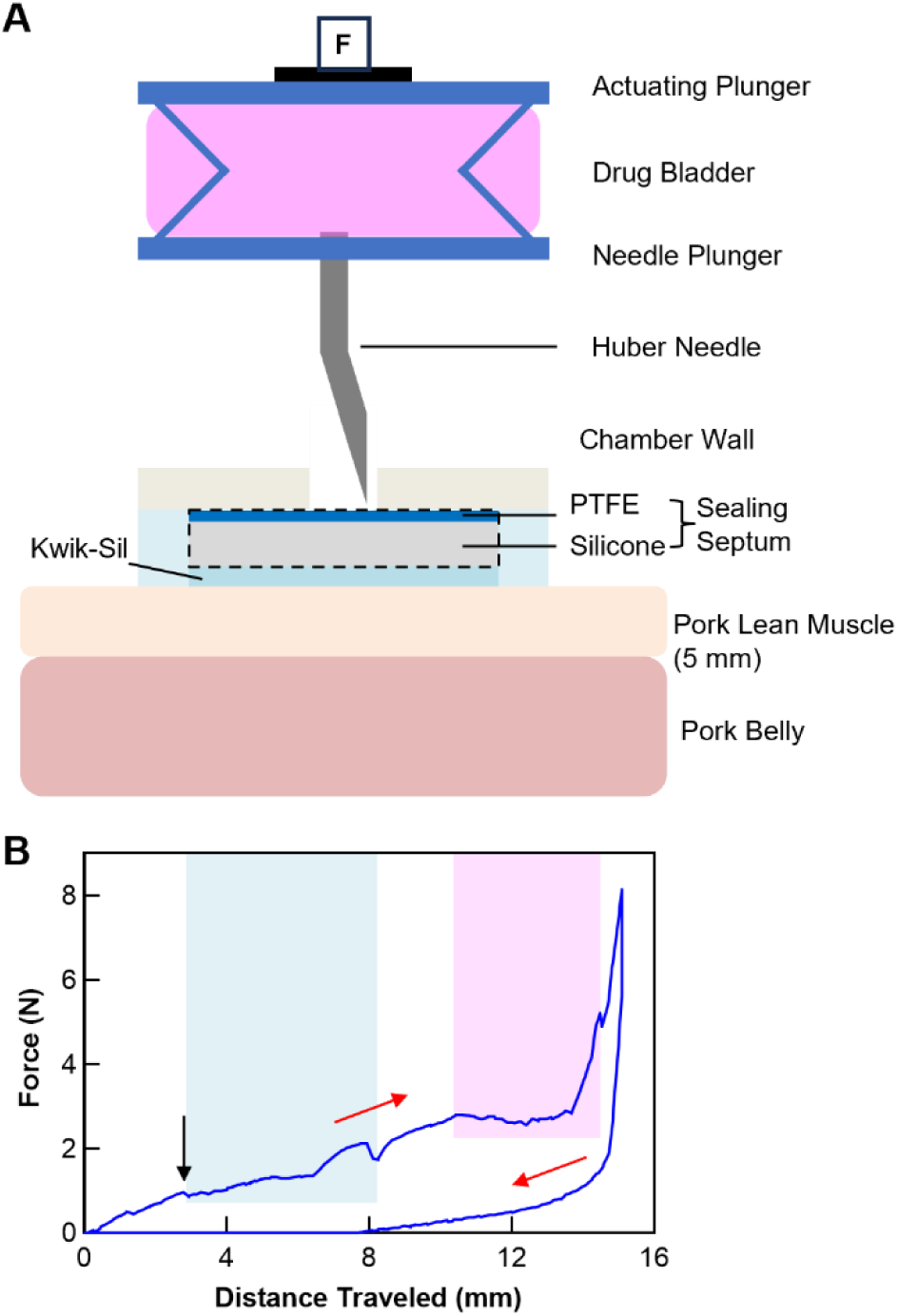
Force during a successive sequence of deploying the needle, injecting the solution, and retracting the needle. **(A)** Schematic image of the experimental setup. The bladder is filled with 1.5 mL of solution for this characterization. **(B)** Plot of force during a successive sequence of deploying the needle into a two-layer pork tissue model, injecting the solution, and retracting the needle. Red arrows indicate the progression of time. The black arrow indicates completion of the process of piercing the sealing septum. The blue area corresponds to the regime for piercing two layers of tissue. The pink area corresponds to the regime for injecting the solution from the bladder into the pork belly.

**Fig. S14.**
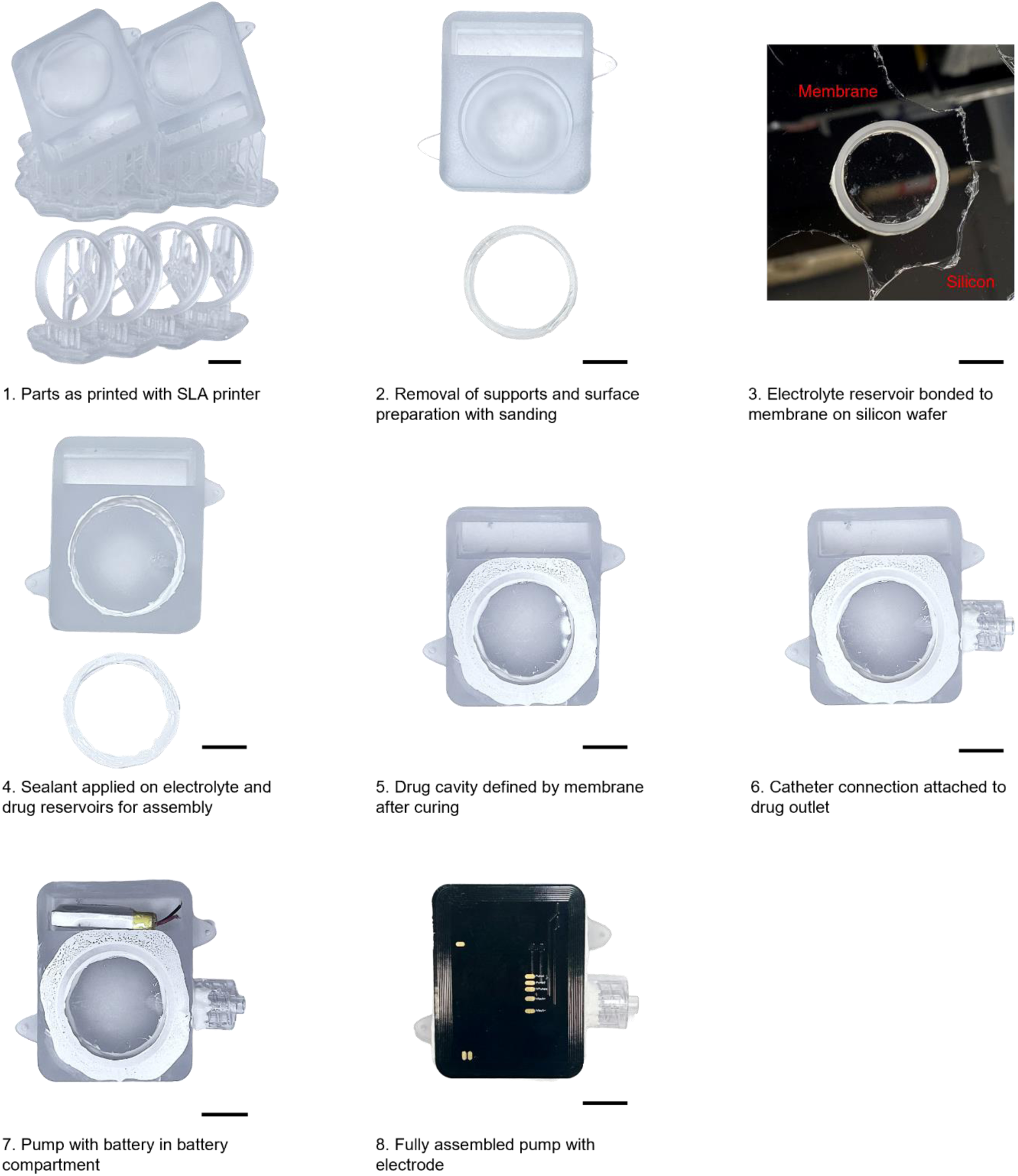
Steps for fabricating intravenous Naloximeter devices. Scale bars, 1 cm.

**Fig. S15.**
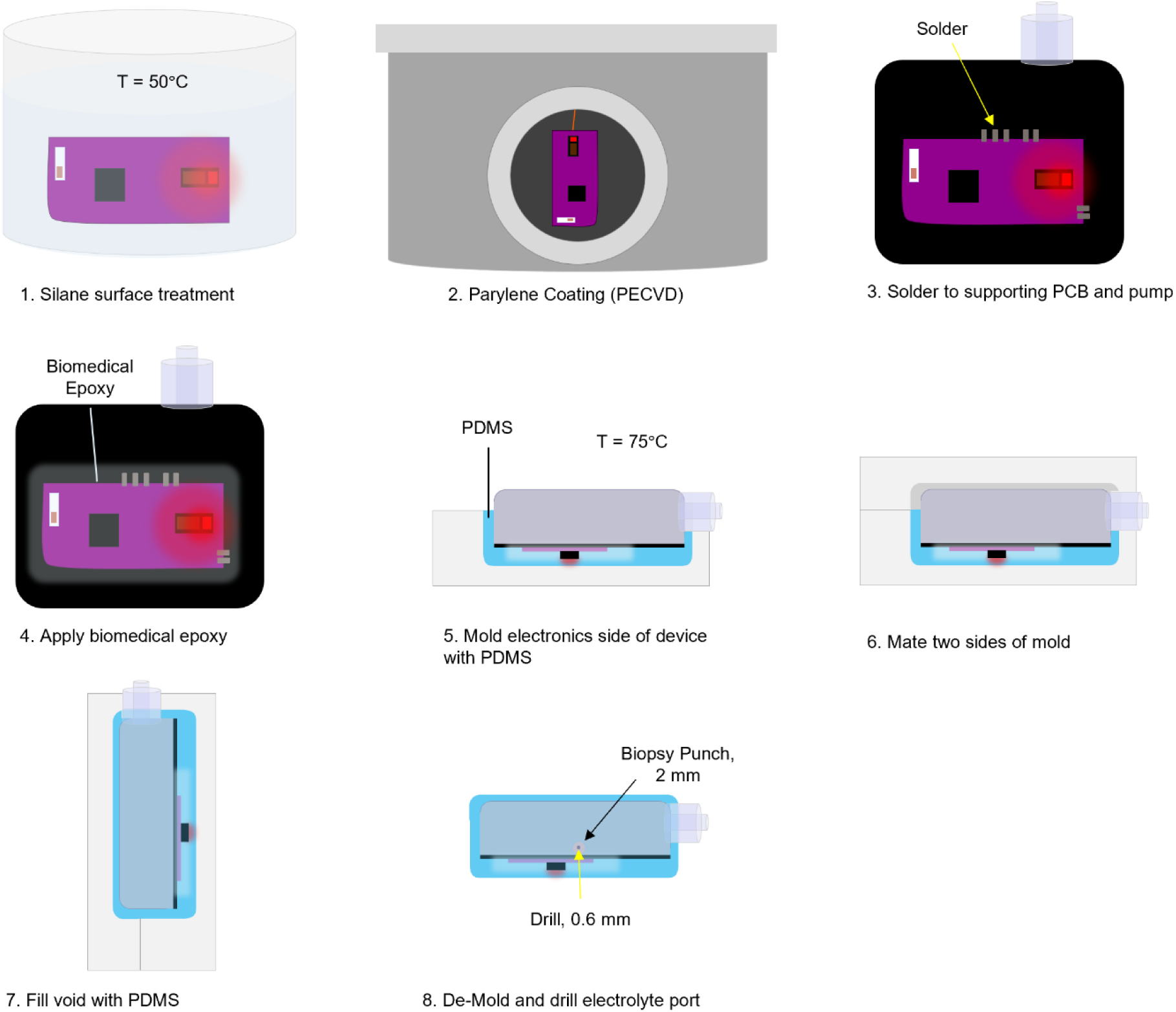
Steps for encapsulating Naloximeter devices, illustrated with the intravenous device.

**Fig. S16.**
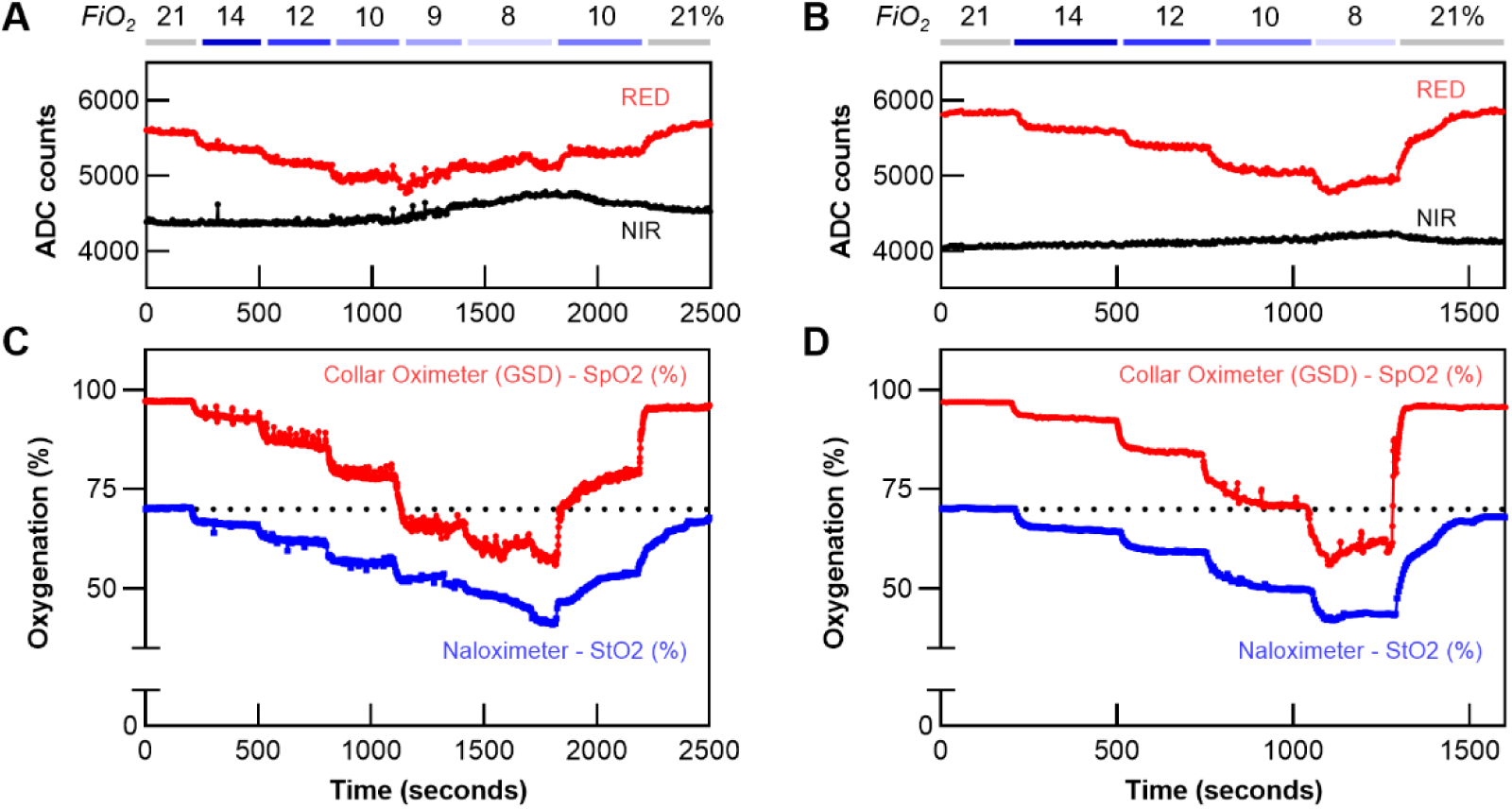
Hypoxia studies in rodent models. **(A, B)** Raw red and near-infrared (NIR) optical signals from an NFC Naloximeter during hypoxia induced by modulation of the fraction of inspired oxygen (FiO_2_). **(C, D)** Corresponding calculated StO_2_ from the Naloximeter and SpO_2_ measured by a gold standard device (GSD, collar oximeter). Each column represents one animal.

**Fig. S17.**
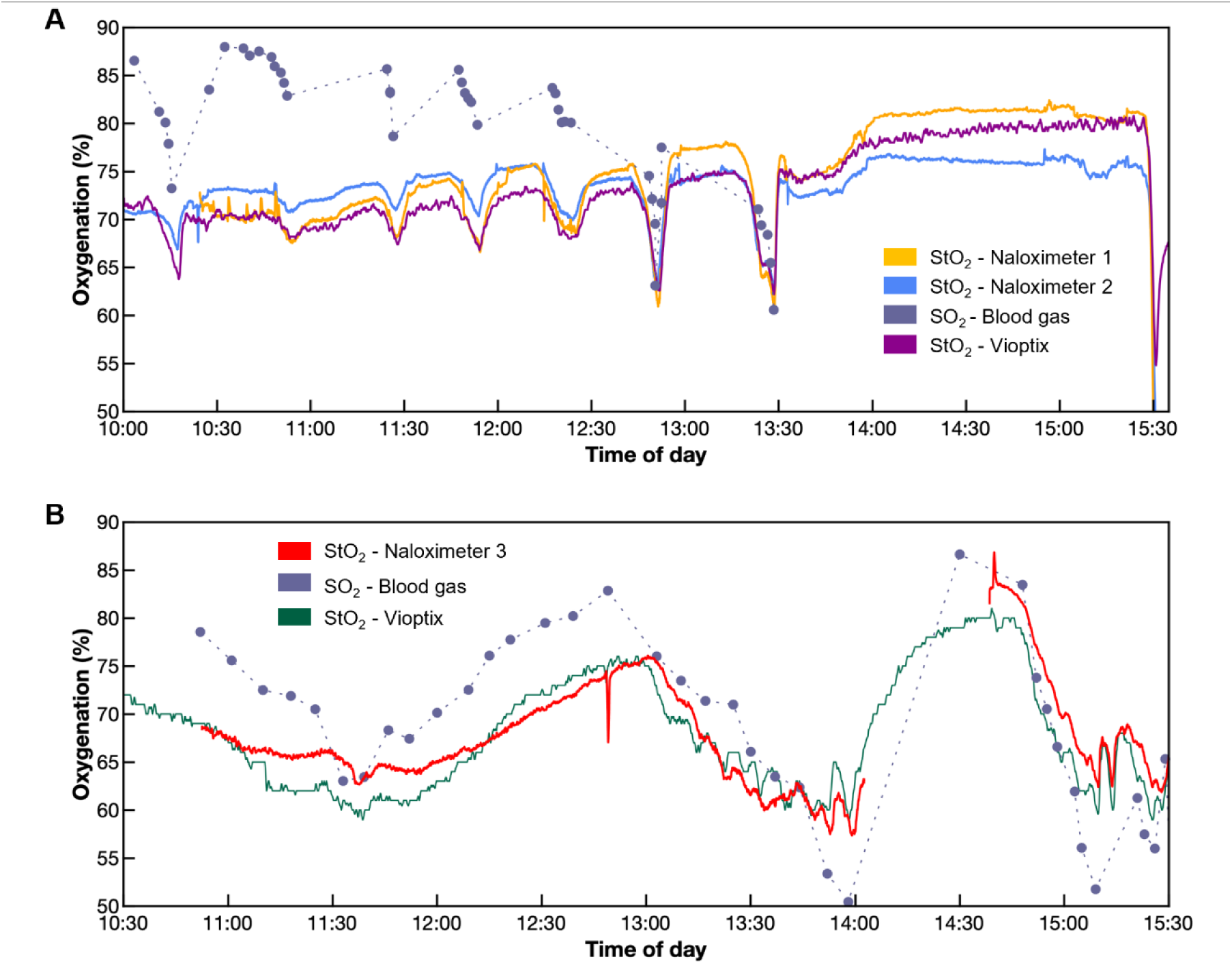
Hypoxia studies in a porcine model with comparison to standard clinical devices and measurements. **(A, B)** Blood gas saturation is calculated as a mixture of 70% venous and 30% arterial blood.

**Fig. S18.**
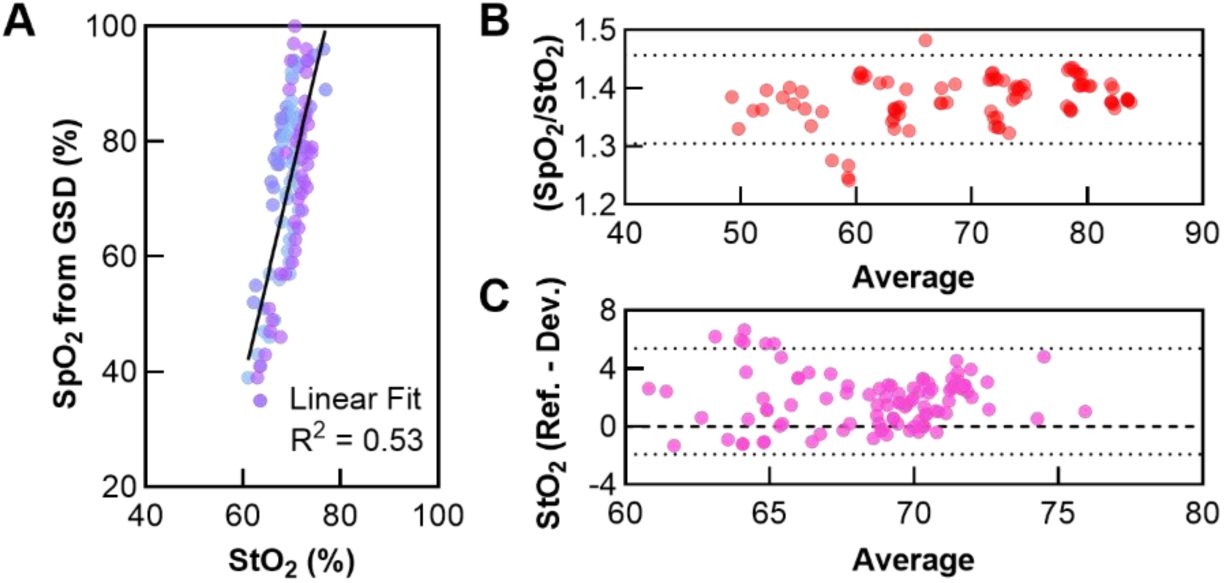
Extended analyses of hypoxia studies in swine and rodent models. **(A)** Comparison of SpO_2_ measured with gold standard device (GSD, clip-on pulse oximeter) and StO_2_ measured with a Naloximeter during hypoxia in a porcine model. **(B)** Bland-Altman plot to compare (ratio versus average) SpO_2_ measured with gold standard device and StO_2_ measured with the Naloximeter. Dashed lines depict the 95% confidence interval. **(C)** Bland-Altman plot to compare (difference versus average) StO_2_ measured with a clinical device (Vioptix) and StO_2_ measured with the Naloximeter in porcine model. Dashed lines depict the 95% confidence interval.

**Fig. S19.**
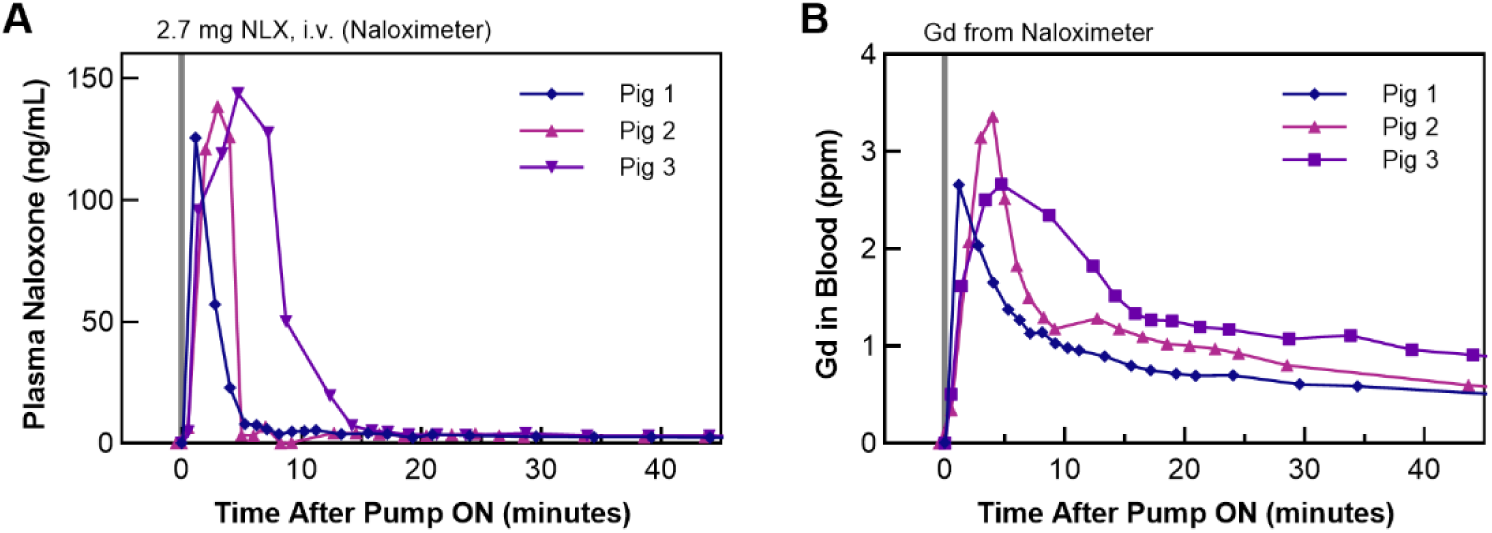
Gadolinium (Gd) and Naloxone pharmacokinetics in porcine models with intravenous Naloximeter devices. **(A)** Naloxone and **(B)** Gd measured in plasma and blood, respectively, after delivery of 3 mL of drug solution from Naloximeter device. *N* = 3 animals, the same blood draw was split into samples for NLX and Gd quantification to facilitate direct comparison between the methods.

**Fig. S20.**
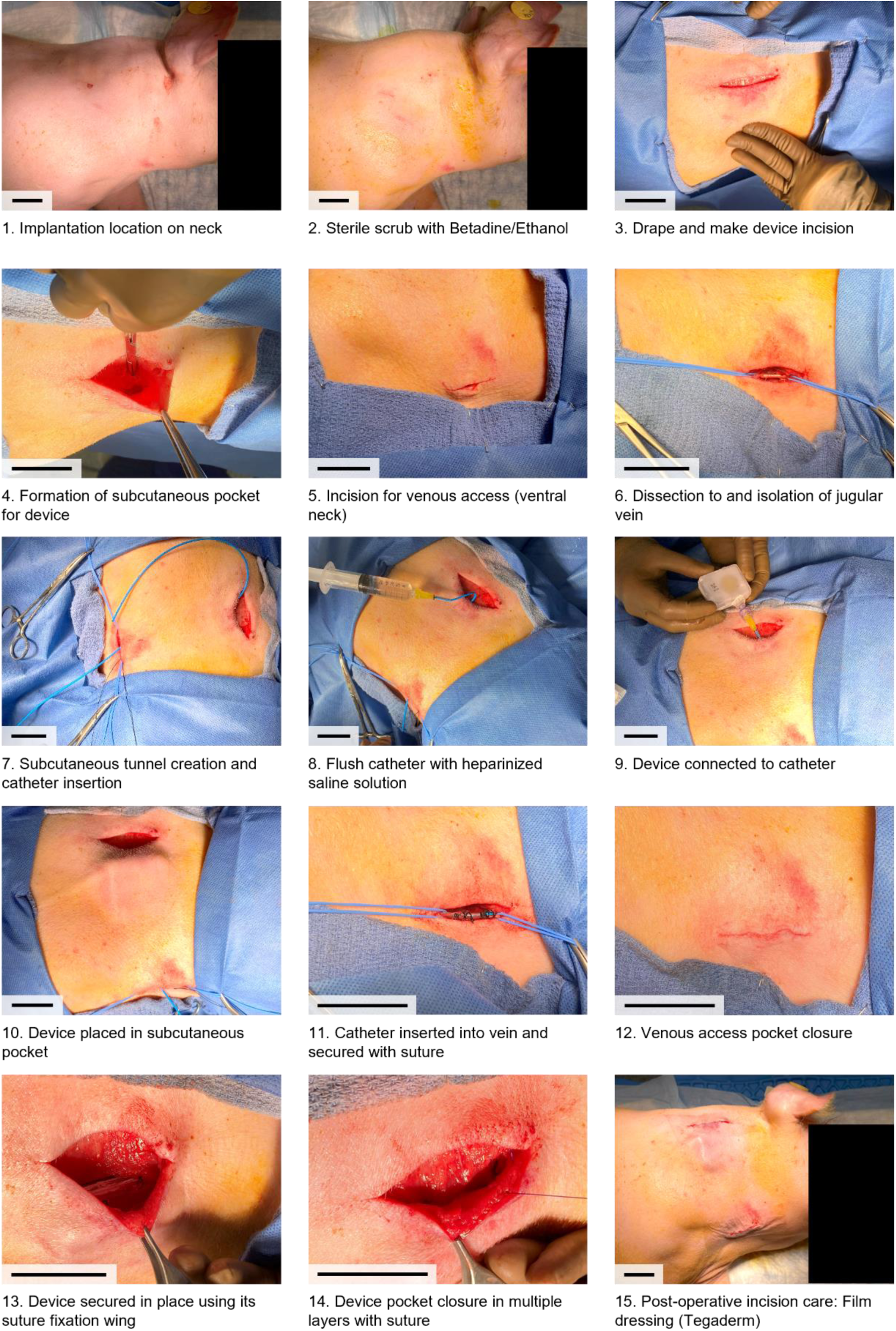
Surgical approach for implanting an intravenous Naloximeter device in the external jugular vein of a pig. Scale bars, 5 cm.

**Fig. S21.**
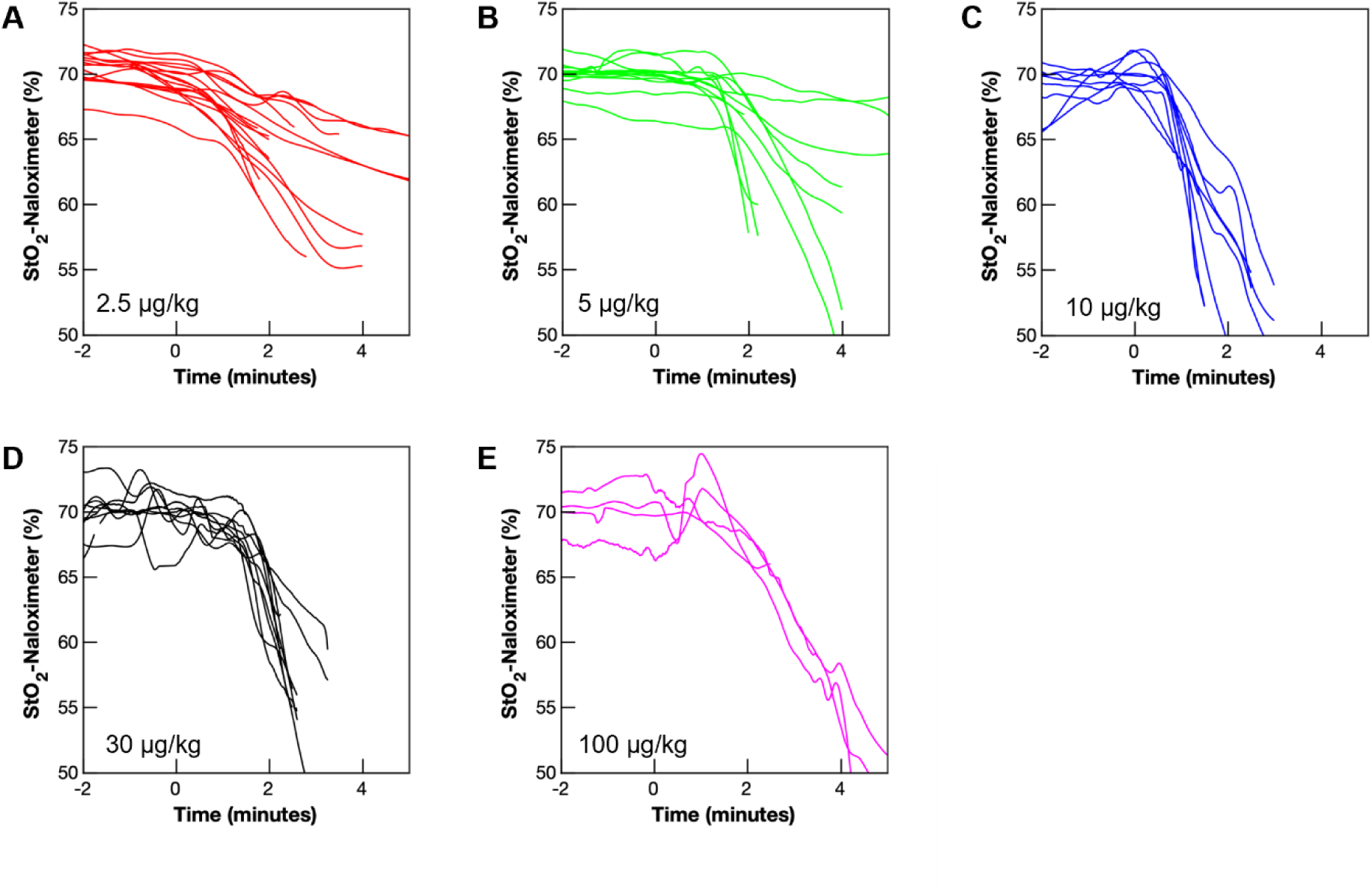
Tissue oxygenation (StO_2_) recorded with Naloximeters in porcine models at different fentanyl doses. **(A)** Fentanyl dose of 2.5 µg/kg, *N* = 6 subjects and *n* = 17 devices. **(B)** Fentanyl dose of 5 µg/kg, *N* = 4 subjects and *n* = 12 devices. **(C)** Fentanyl dose of 10 µg/kg, *N* = 9 subjects and *n* = 9 devices. **(D)** Fentanyl dose of 30 µg/kg, *N* = 6 subjects and *n* = 10 devices. **(E)** Fentanyl dose of 100 µg/kg, *N* = 4 subjects and *n* = 4 devices. Fentanyl was administered at time = 0. Overall *N* = 29 subjects, *n* = 52 devices.

**Fig. S22.**
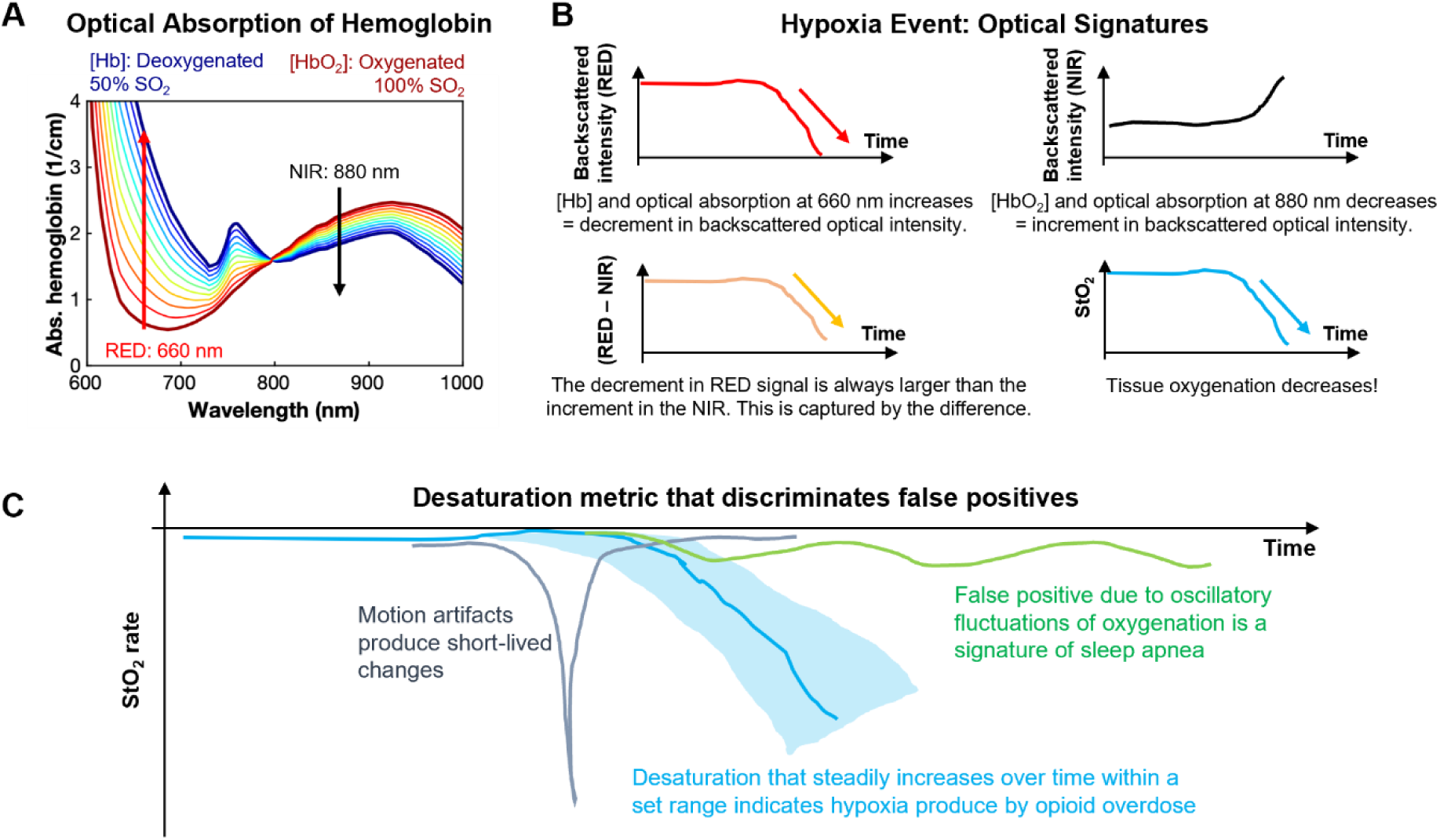
Physiological dynamics during hypoxia produce measurable signatures that contrast with false positives produced by motion artifacts and sleep apneas. **(A)** Optical absorption of hemoglobin depends on the concentration of its oxygenated (HbO_2_) and deoxygenated (Hb) states. Based on data from ref (*50*). Graph shows the change in the optical absorption as hemoglobin oxygenation changes from 100% SO_2_ (dark red) to 50% SO*_2_* (dark blue). Differential spectroscopy, at the red = 660 nm (***λ***_***1***_) and near-infrared (NIR) = 880 nm (***λ***_***2***_) wavelengths, is used to calculate the tissue oxygenation (***StO***_***2***_). These signals capture the physiological dynamics. **(B)** Hypoxia is accompanied by clear changes in the optical signal and the calculated StO_2_. These data produce a set of variables to detect desaturation. **(C)** Typical confounding events follow from motion artifacts and behaviors associated with sleep apnea. Motion artifacts can lead to false positives that are typically eliminated based on criteria associated with the rate of change of the optical signal. Episodes of desaturation due to sleep apnea can be eliminated by their characteristic oscillatory patterns and desaturation rate.

**Fig. S23.**
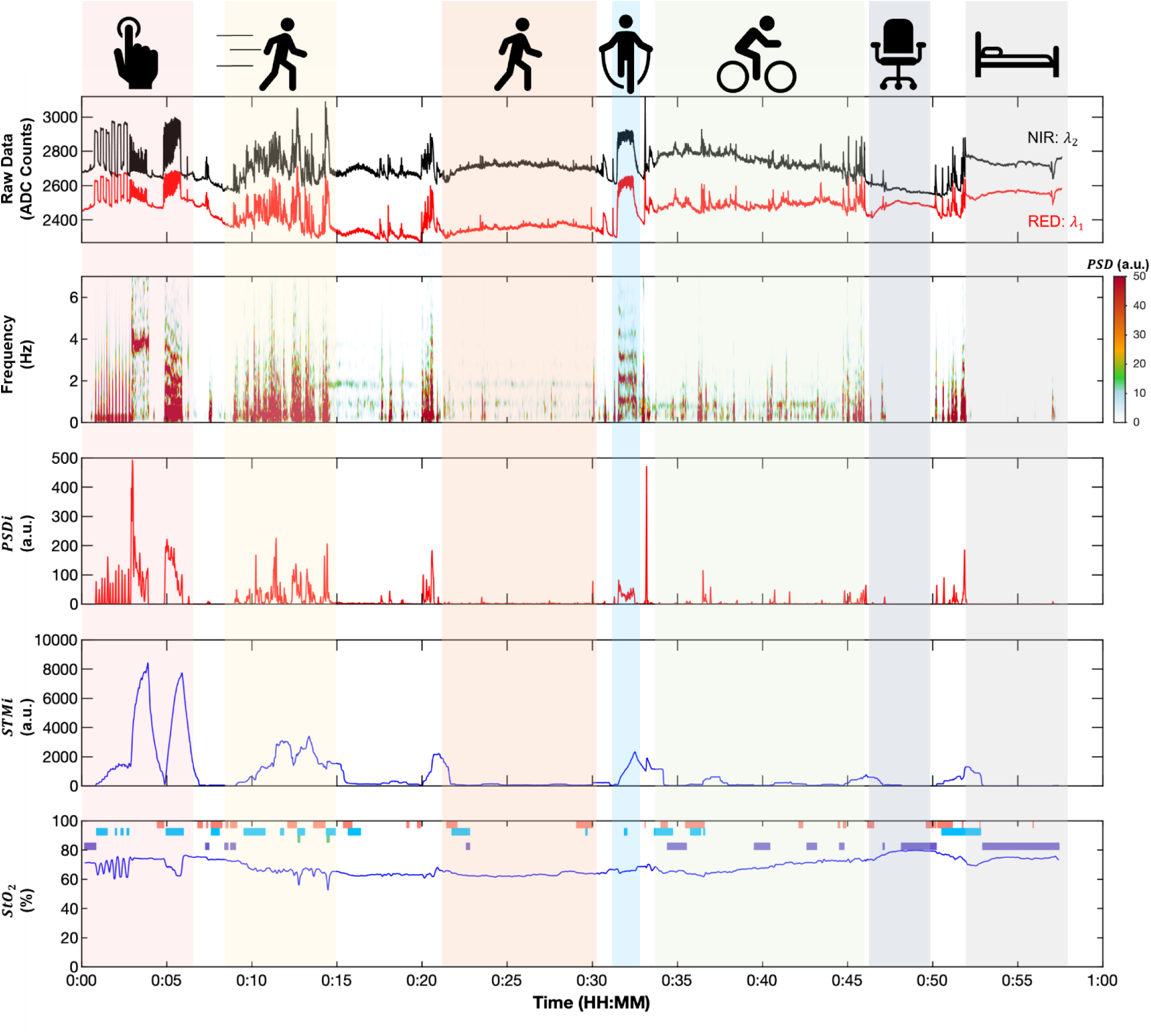
Data from cutaneous optical sensors in humans associated with different types of physical activity. Data recorded with dual-wavelength optical sensor mounted on the skin in the arm of a human subject volunteer performing a series of movements. Movement sequence: Applying pressure to the skin (red), running (yellow), walking (orange), aerobic jumping (blue), cycling (green), resting while seated (blue-gray), resting while supinated (gray). Plots from top-down: red and near-infrared (NIR) optical signals; spectrogram; power spectral density index (***PSDi***) in the 1-5 Hz frequency band; short-term motion index (***STMi***, 60 s) and calculated tissue oxygenation (***StO***_***2***_).

**Fig. S24.**
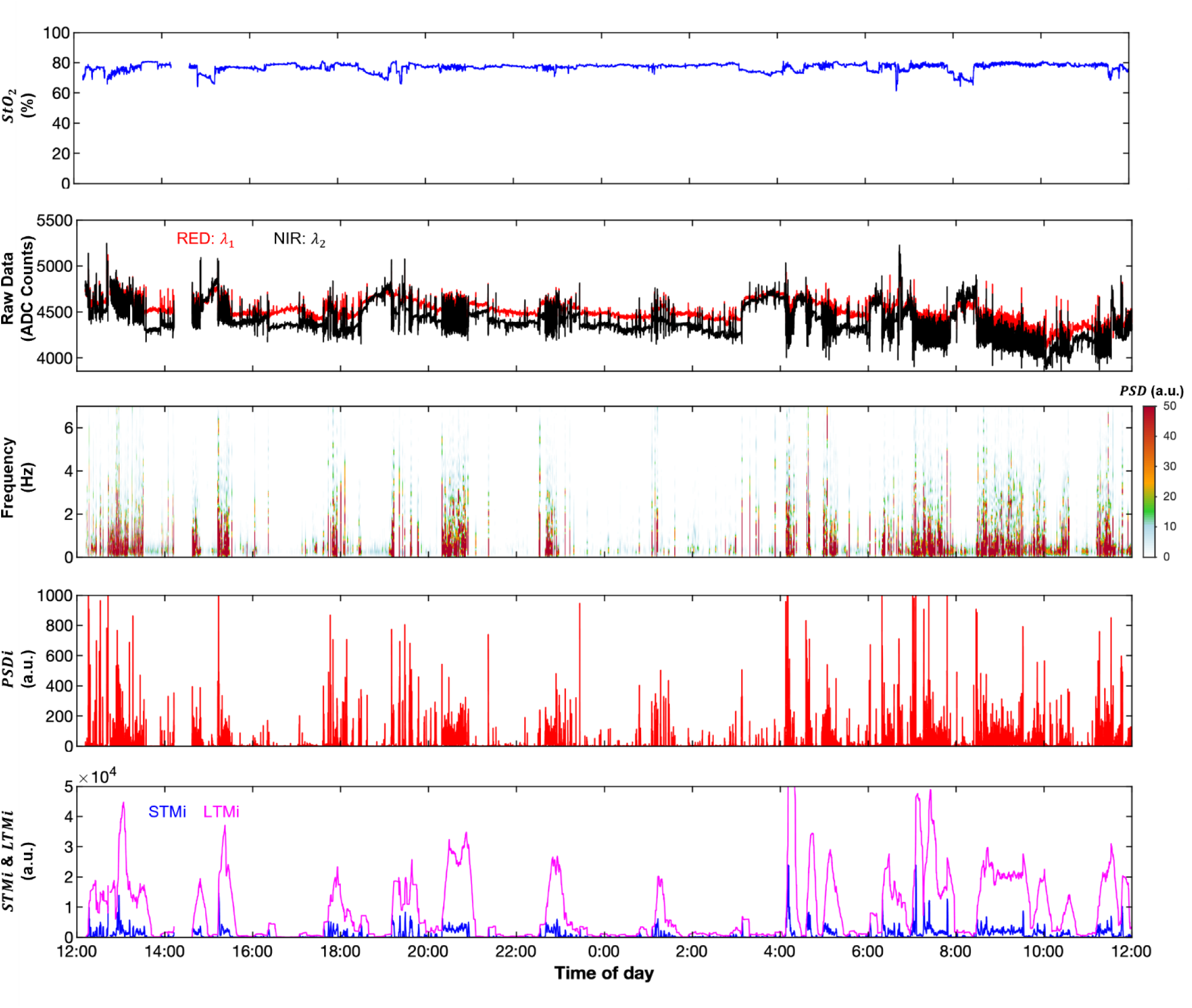
Cumulative power spectral density index for defining episodes of physical activity. Data recorded with dual-wavelength optical sensor in a freely moving porcine model over 24 hours. From top-down: calculated tissue oxygenation (***StO***_***2***_); red and near-infrared (NIR) raw optical signals; spectrogram; power spectral density index (***PSDi***) in the 1-5 Hz frequency band; short-term motion index (***STMi***, 60 s) and long-term motion index (***LTMi***, 10 min). Episodes of high motion index, apparent in the raw data, corresponds with the episode of increased ***LTMi***, power spectral density and ***PSDi***.

**Fig. S25.**
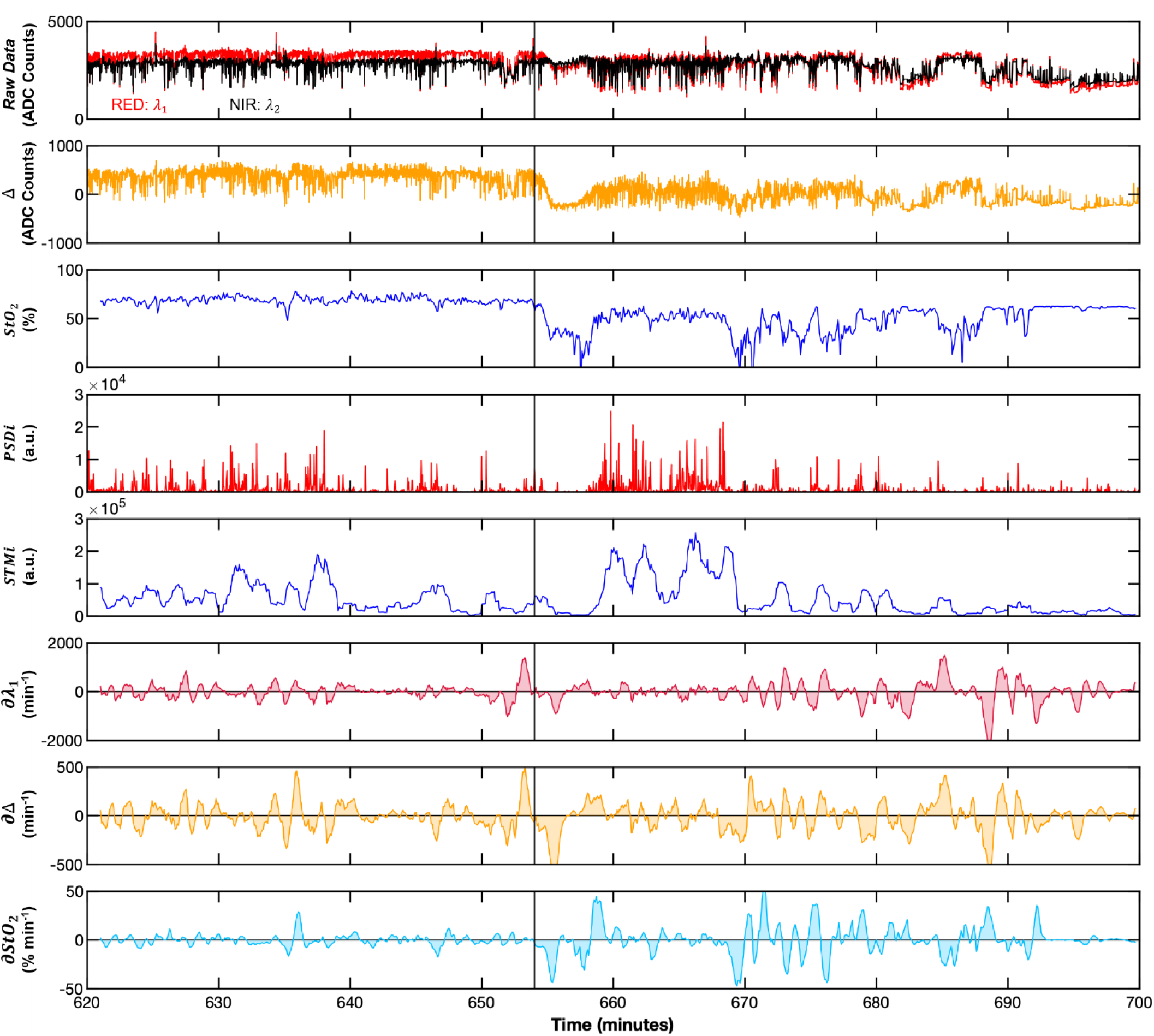
Optical sensor data, and its multivariable derivatives, during fentanyl overdose in a freely moving pig. From top-down: raw data of the red (RED, ***λ***_***1***_) and near-infrared (NIR, ***λ***_***2***_) optical signals; difference between these signals (Δ = ***λ***_***1***_ − ***λ***_***2***_); calculated tissue oxygenation (***StO***_***2***_); power spectral density index (***PSDi***) in the 1–5 Hz frequency band; short-term motion index (***STMi***); rate of change (***∂***) of red (***∂λ***_***1***_); difference (***∂***Δ); and tissue oxygenation (***∂StO***_***2***_). The time of fentanyl administration is indicated with a vertical line.

**Fig. S26.**
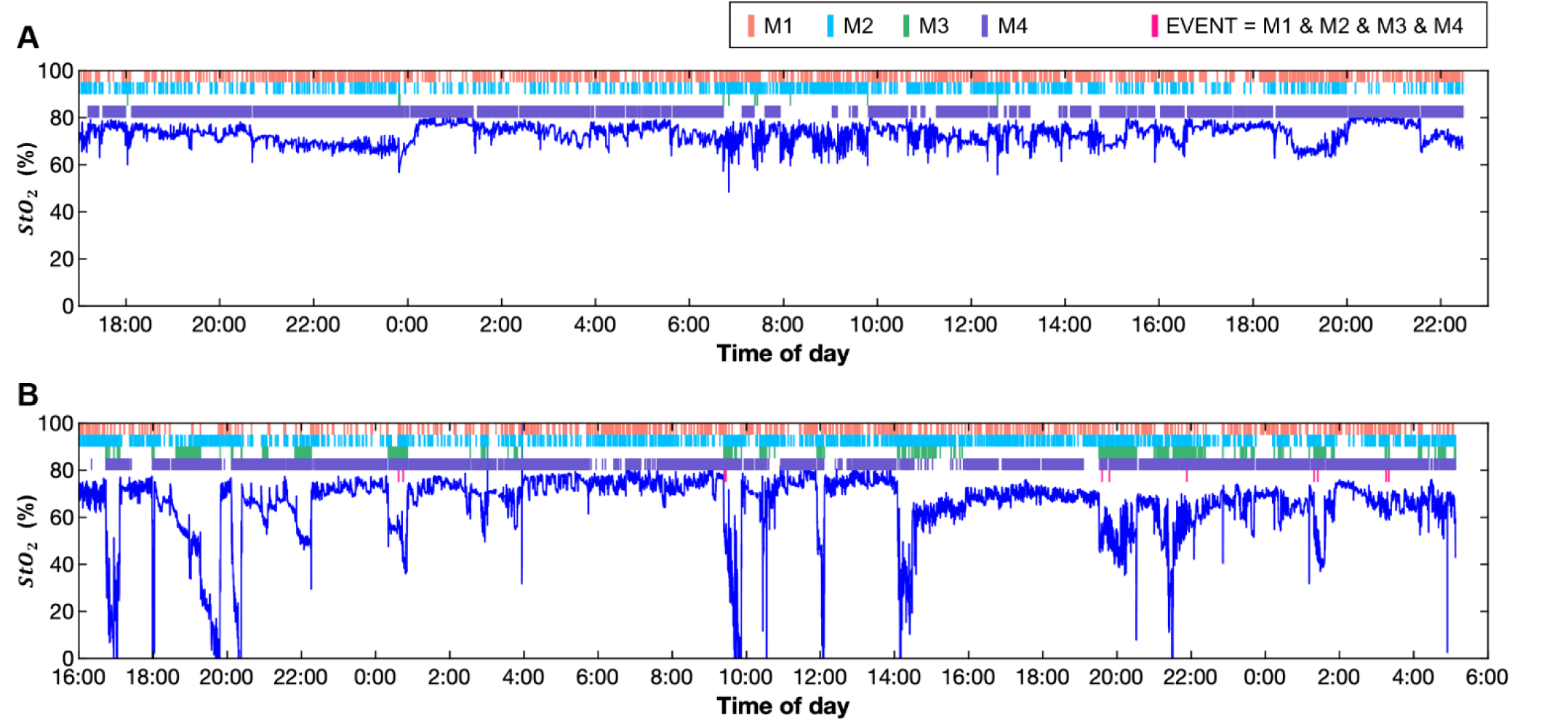
Extended recordings of oxygenation (StO_2_) with Naloximeters in freely moving pigs. **(A, B)** Colored bars represent instances where the logical metrics (M1 – M4) are true. EVENTS (M1 & M2 & M3 & M4) are registered in (B) but without satisfying the sequential condition to quality for an overdose. *N* = 2 animals with Naloximeters implanted in **(A)** the neck (jugular approach) for 15 days and **(B)** the flank (mammary approach) for 32 days.

**Fig. S27.**
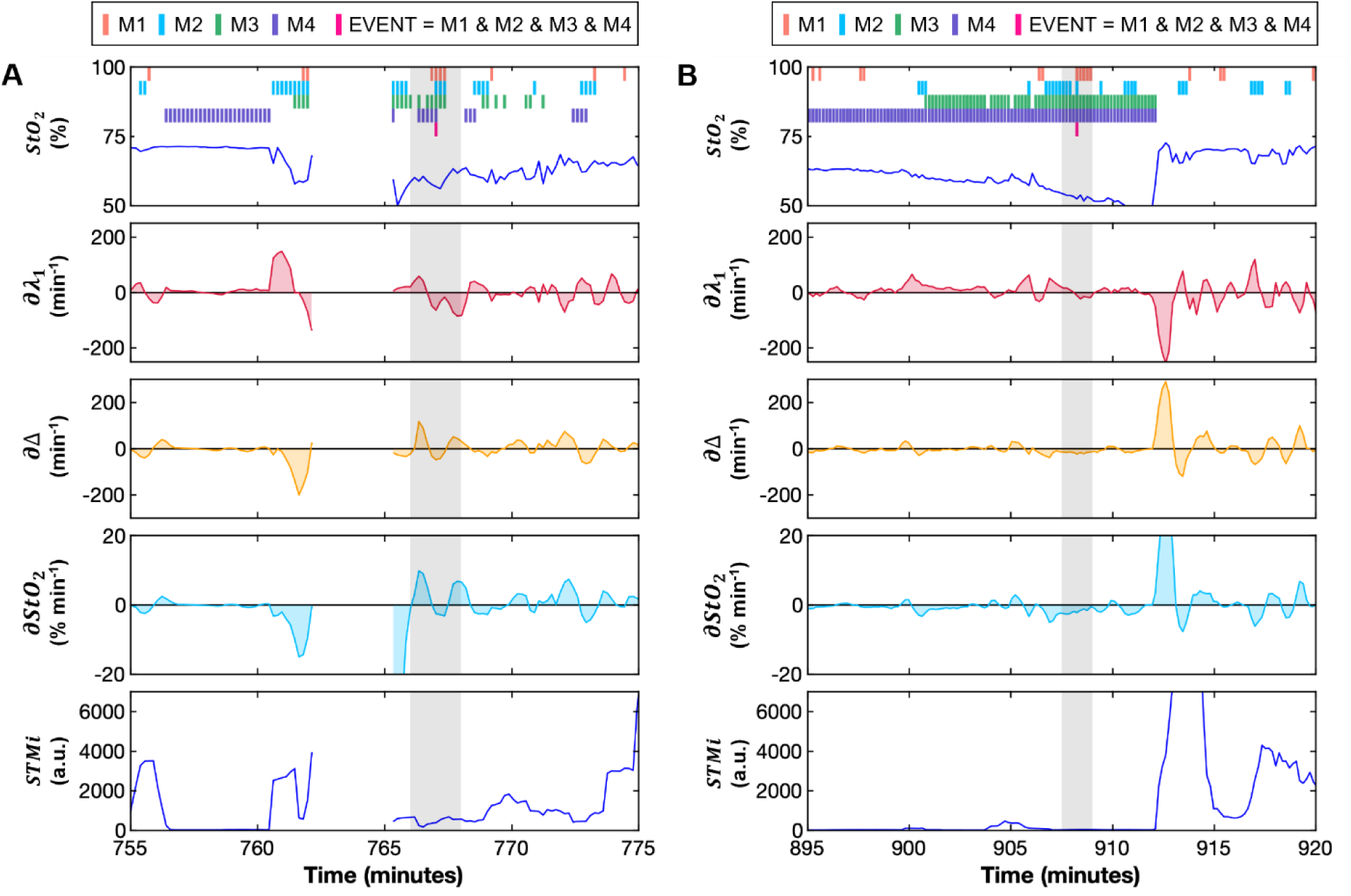
Metrics associated with algorithm validation over 24 hours at selected times where an event was detected. **(A, B)** From top-down: calculated tissue oxygenation (***StO***_***2***_) showing when the logical metric conditions (M1, M2, M3, and M4) are true; rate of change (***∂***) of red (***∂λ***_***1***_); difference (***∂***Δ); and tissue oxygenation (***∂StO***_***2***_); short-term motion index (***STMi***). The instances of EVENTs (M1 & M2 & M3 & M4) are isolated and did not produce a WARNING (three consecutive occurrences). Each column represents a distinct episode.

**Fig. S28.**
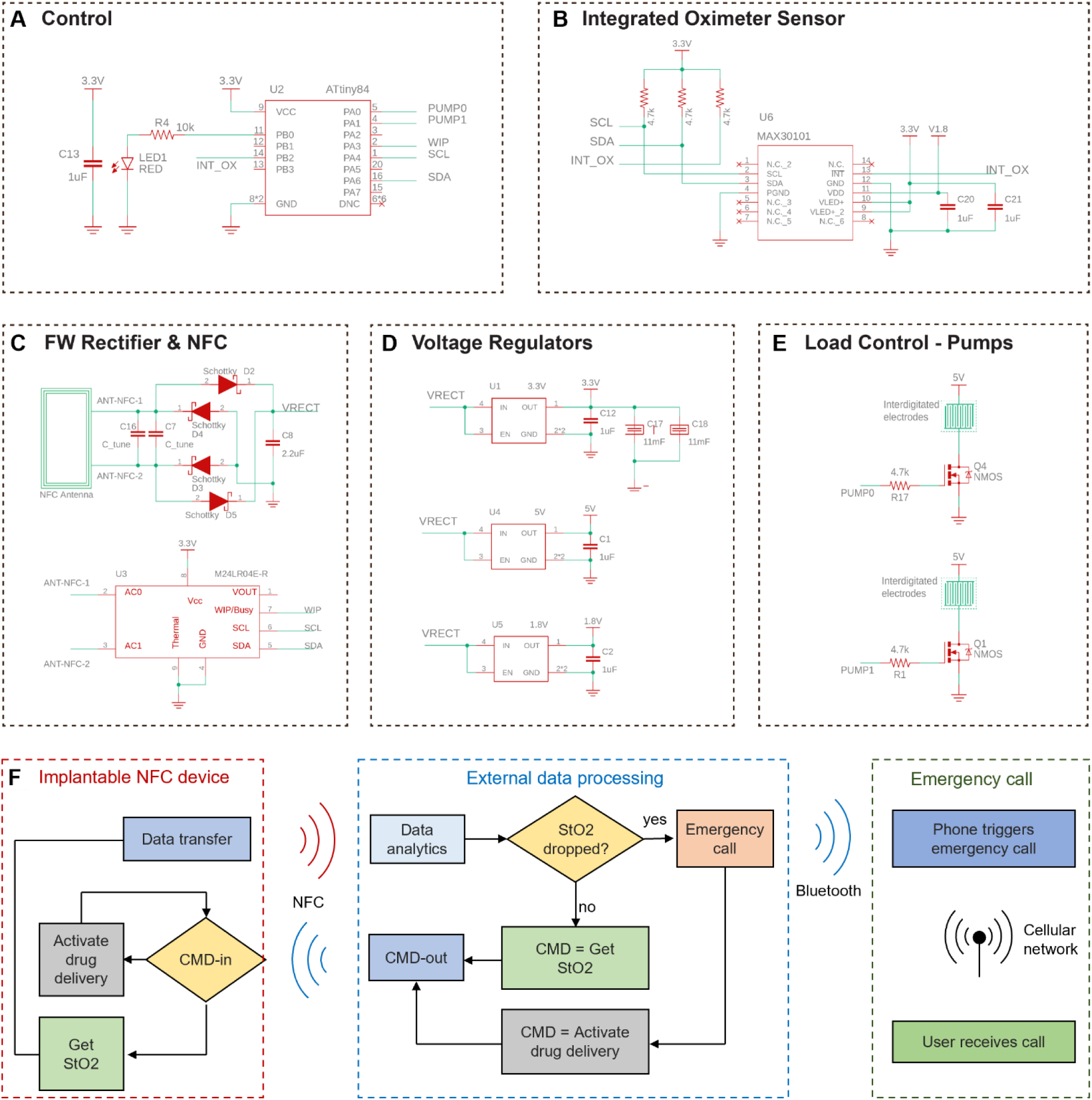
Electronic circuit and functional block diagrams for the battery-free, NFC Naloximeter for rodents. **(A)** The ATTiny84 microcontroller controls the operation of the device and the manages the wireless communication. **(B)** The integrated optical sensor MAX30101 connects with the MCU using I^2^C communication. **(C)** RF front end containing a full-wave rectifier and an RF random access memory used for bidirectional NFC communication. **(D)** A voltage regulator supplies 3.3 V to supercapacitor bank (top). A 5 V regulator provides voltage to the electrolytic pumps (center) and a 1.8 V regulator supplies voltage to the dual-wavelength optical sensor (bottom). **(E)** Pump control system with MOSFETs and interdigitated gold electroplated electrodes. Each pump is controlled independently. **(F)** Functional block diagram of the NFC Naloximeter that demonstrates the close-loop implementation in rodents. From left to right: Implanted NFC device; external data processing unit implemented in MATLAB; emergency call management system.

**Fig. S29.**
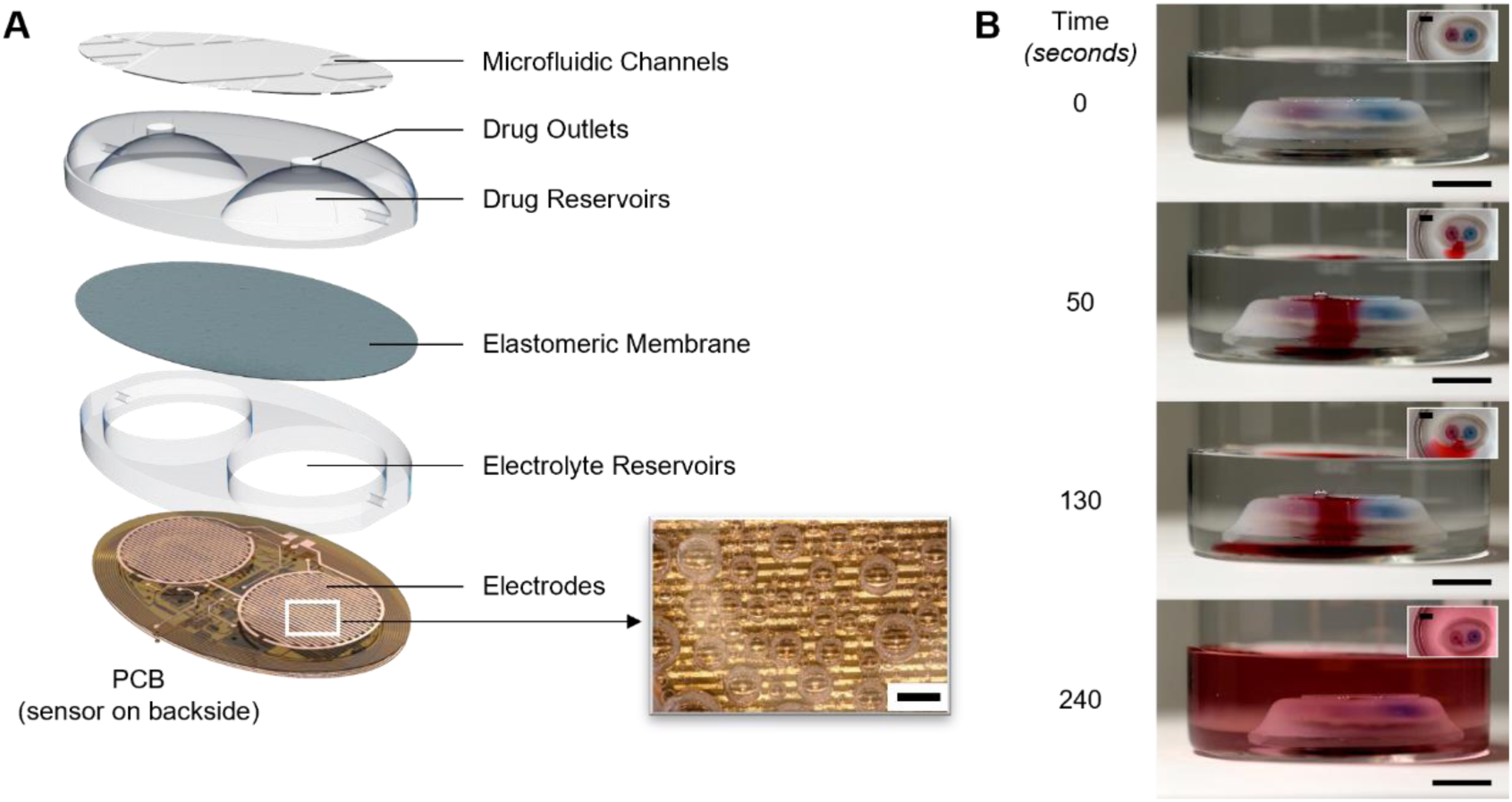
Battery-free, NFC Naloximeter for rodents. **(A)** Exploded view schematic illustration of the device with inset image of electrolytic gas generation by interdigitated electrodes. Scale bar, 1 mm. **(B)** Photographs that show drug delivery, using food dye as drug proxy for purposes of visualization. Scale bars, 1 cm.

**Fig. S30.**
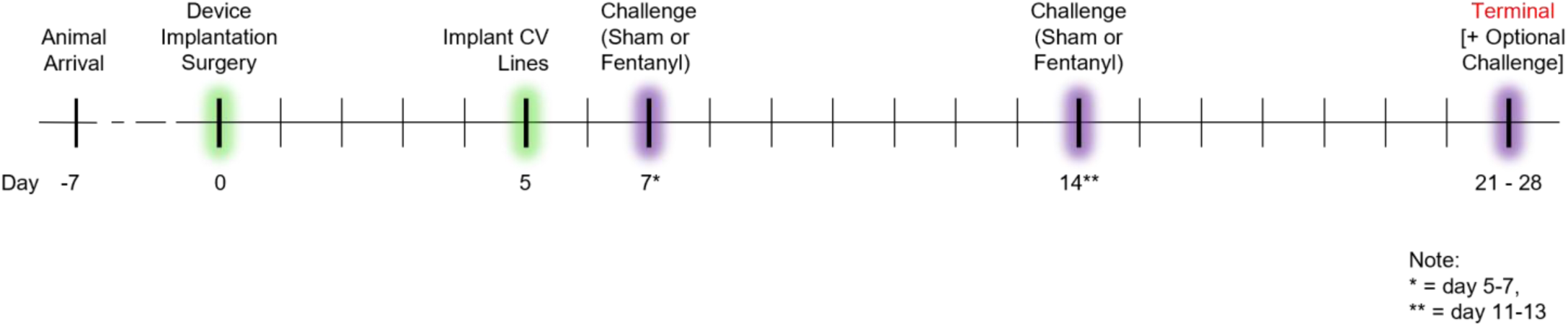
General timeline for large animal survival studies.

**Fig. S31.**
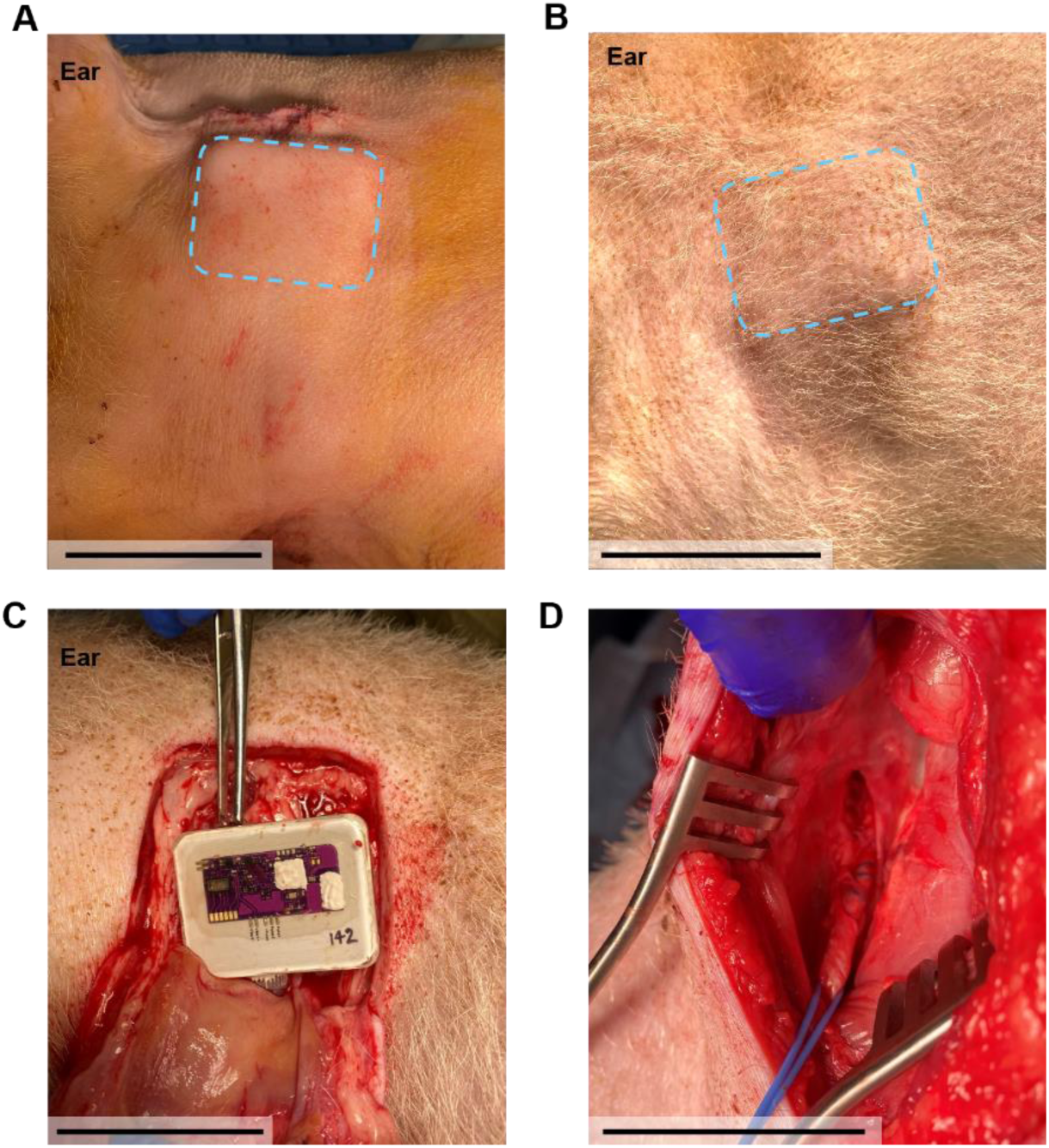
Healing progression for an intravenous Naloximeter implanted in the jugular vein of a pig model. Photographs near the area **(A)** immediately after implantation and **(B)** on day 44 post-implantation. Dashed boxes denote the device outline. Photographs of the **(C)** implanted Naloximeter and **(D)** catheter intact in the jugular vein at day 44 post-implantation. Scale bars, 5 cm.

**Fig. S32.**
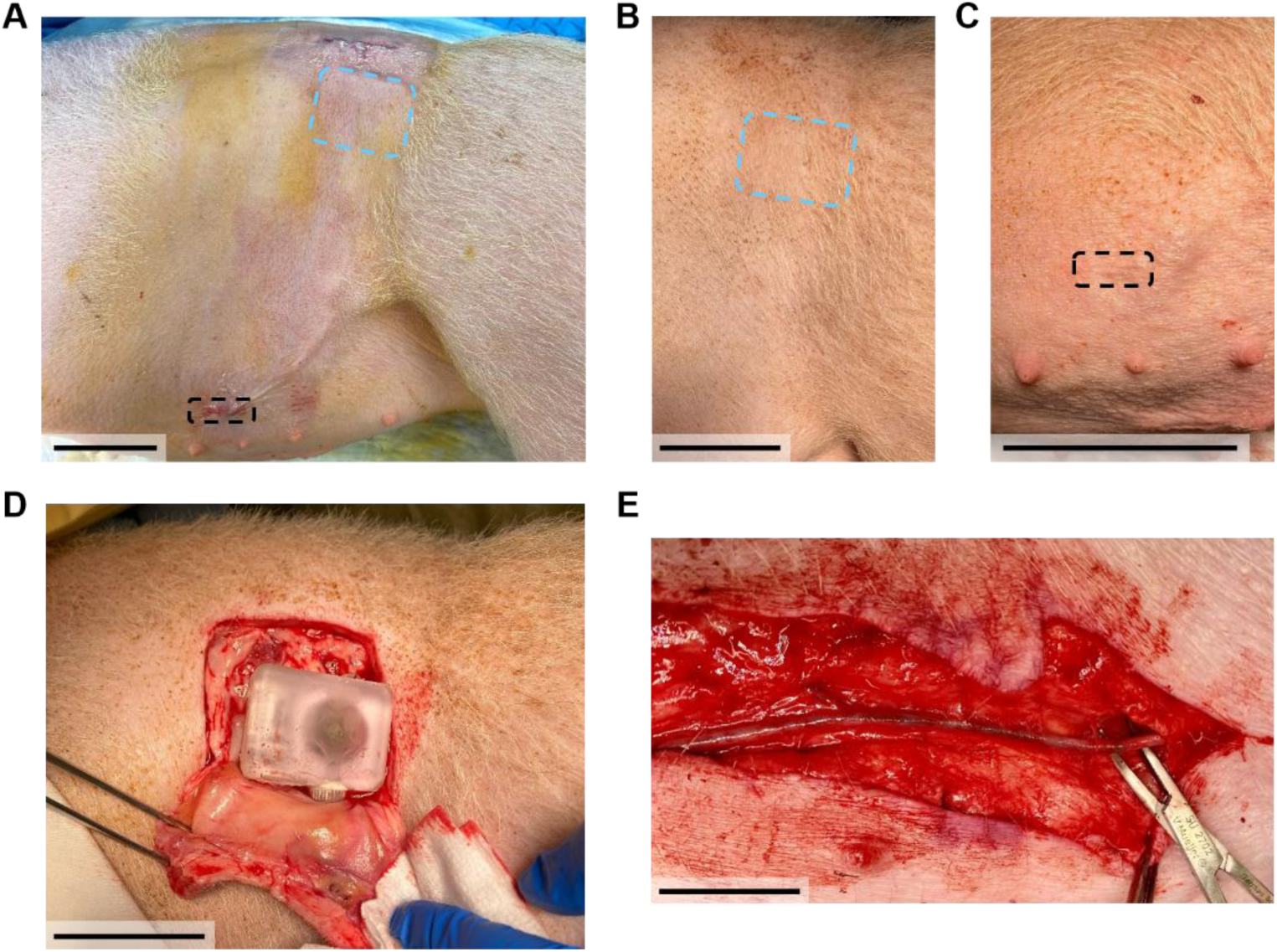
Healing progression for an intravenous Naloximeter implanted in the mammary vein. Photographs near the area **(A)** immediately after implantation and **(B, C)** on day 44 post-implantation at the device and catheter locations, respectively. Dashed blue boxes denote the device outline. The dashed black boxes denote the vascular access incision. Photographs of the **(D)** implanted device and **(E)** catheter intact in the mammary vein at day 44 post-implantation. Scale bars, 5 cm.

**Fig. S33.**
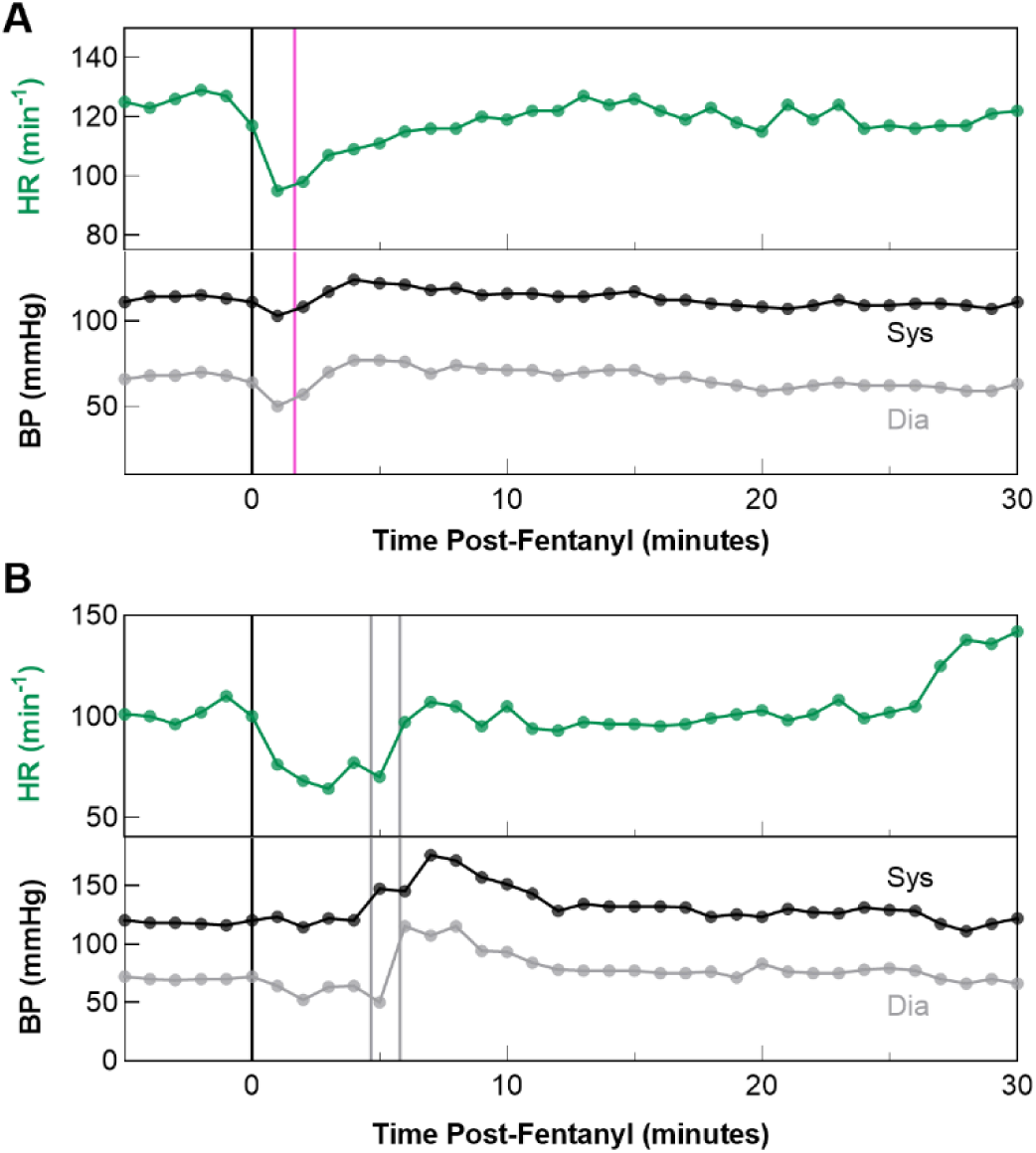
Additional vital signs from (A) closed-loop rescue with Naloximeter and (B) the control case without Naloximeter in anesthetized pig model at 2.5 μg/kg fentanyl dose. Top: Heart rate from electrocardiogram (ECG), bottom: invasive blood pressure. Black line denotes time of fentanyl (i.v.) dose, pink line denotes time of closed-loop NLX (i.v.), gray line denotes time of manual rescue NLX (i.v.). Data in A, B correspond to overdoses in Fig. 4E, 4F, respectively.

**Fig. S34.**
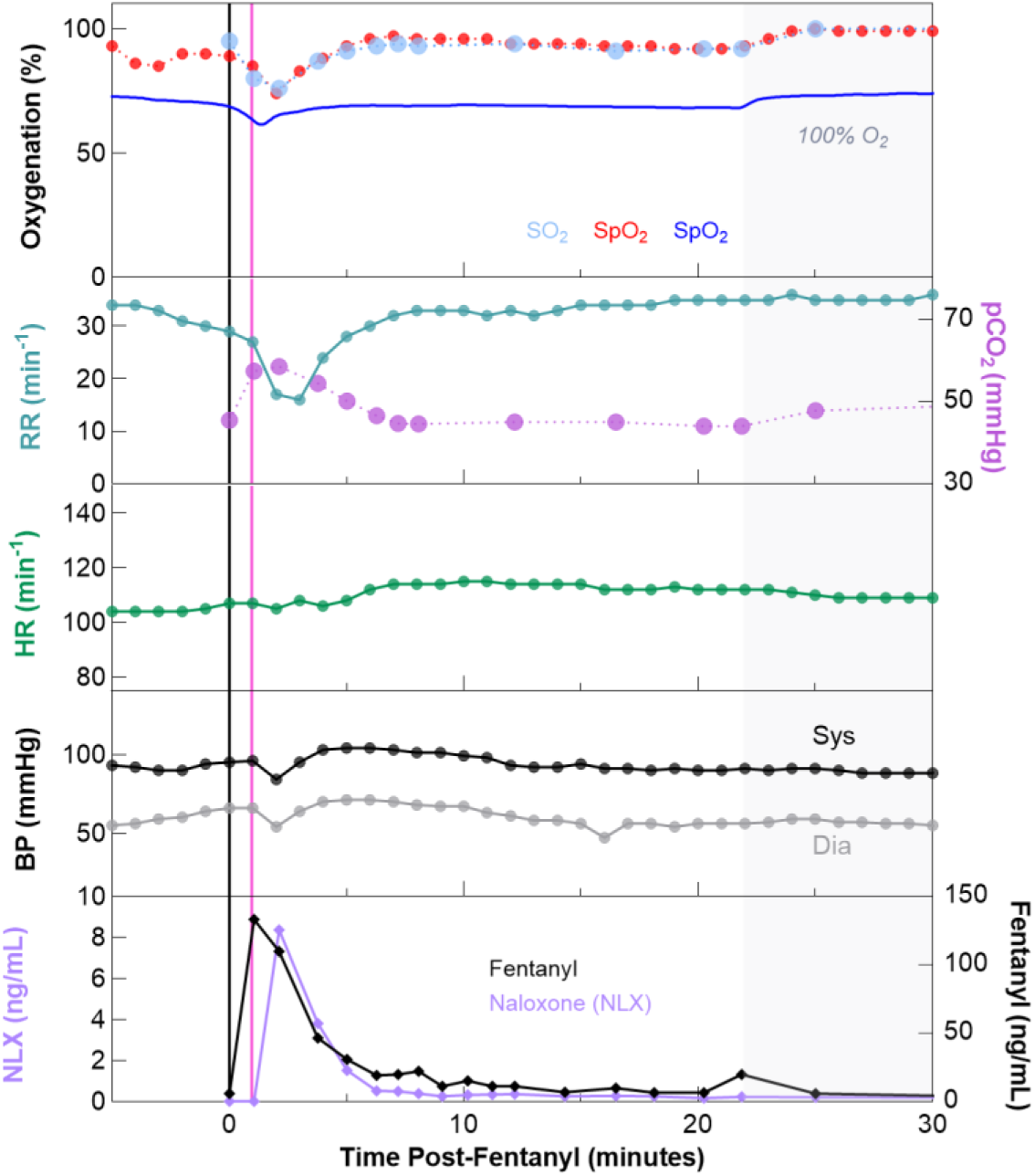
Closed-loop rescue demonstration with intravenous Naloximeter in anesthetized pig model at 2.5 μg/kg fentanyl dose. From top-down: Oxygenation (SO_2_: Arterial Blood Gas, SpO_2_: commercial pulse oximeter, StO_2_: Naloximeter), respiratory vitals, heart rate from electrocardiogram (ECG), invasive blood pressure, and pharmacokinetics of naloxone (NLX) and fentanyl. Black line denotes time of fentanyl (i.v.) dose, pink line denotes time of closed-loop NLX.

**Fig. S35.**
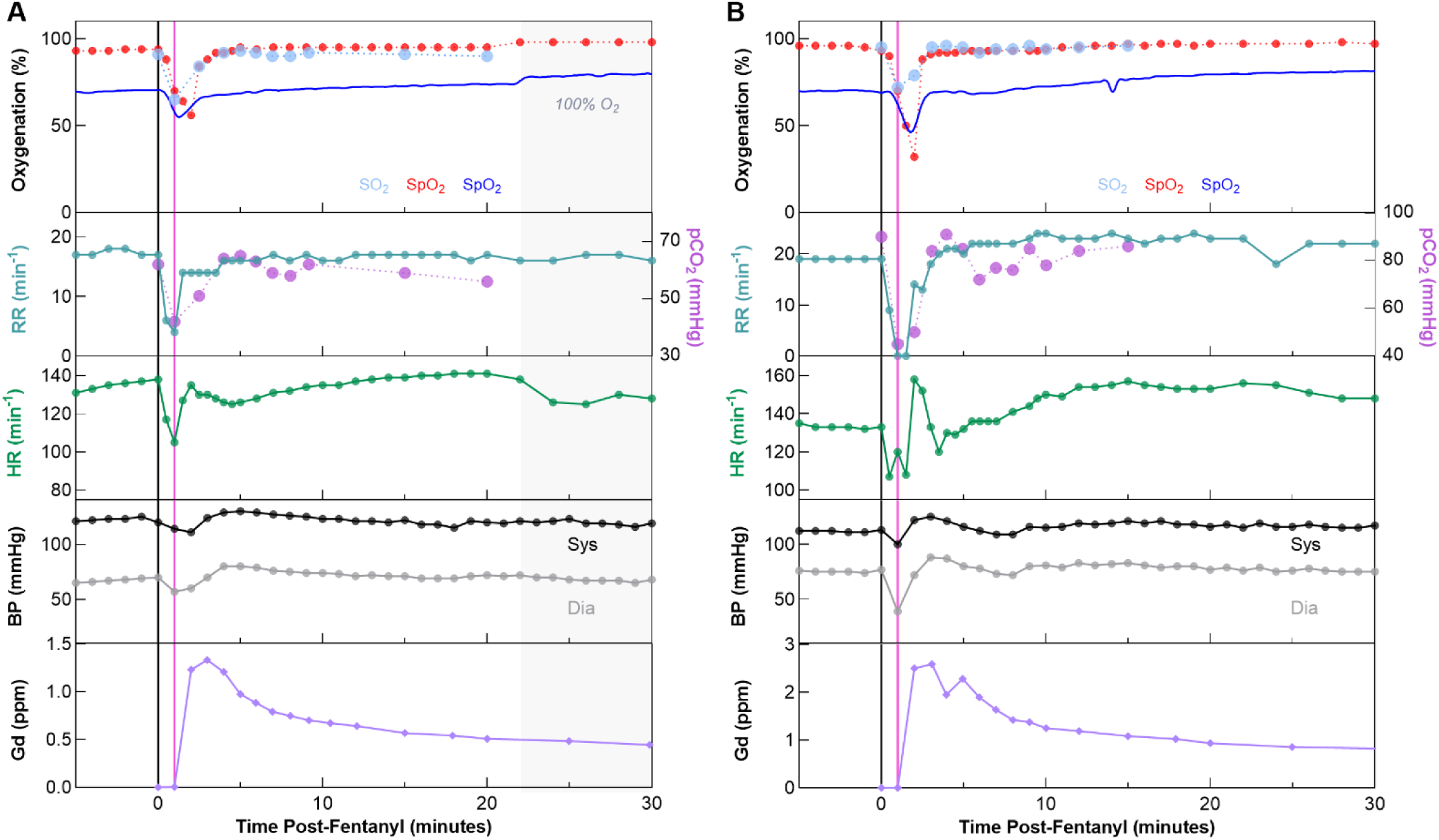
Closed-loop rescue demonstrations with intravenous Naloximeter in anesthetized pig model at 5 μg/kg fentanyl dose. **(A, B)** From top-down: Oxygenation (SO_2_: Arterial Blood Gas, SpO_2_: commercial pulse oximeter, StO_2_: Naloximeter), respiratory vitals, heart rate from electrocardiogram (ECG), invasive blood pressure, and gadolinium pharmacokinetics. Black line denotes time of fentanyl (i.v.) dose, pink line denotes time of closed-loop NLX (i.v.).

**Fig. S36.**
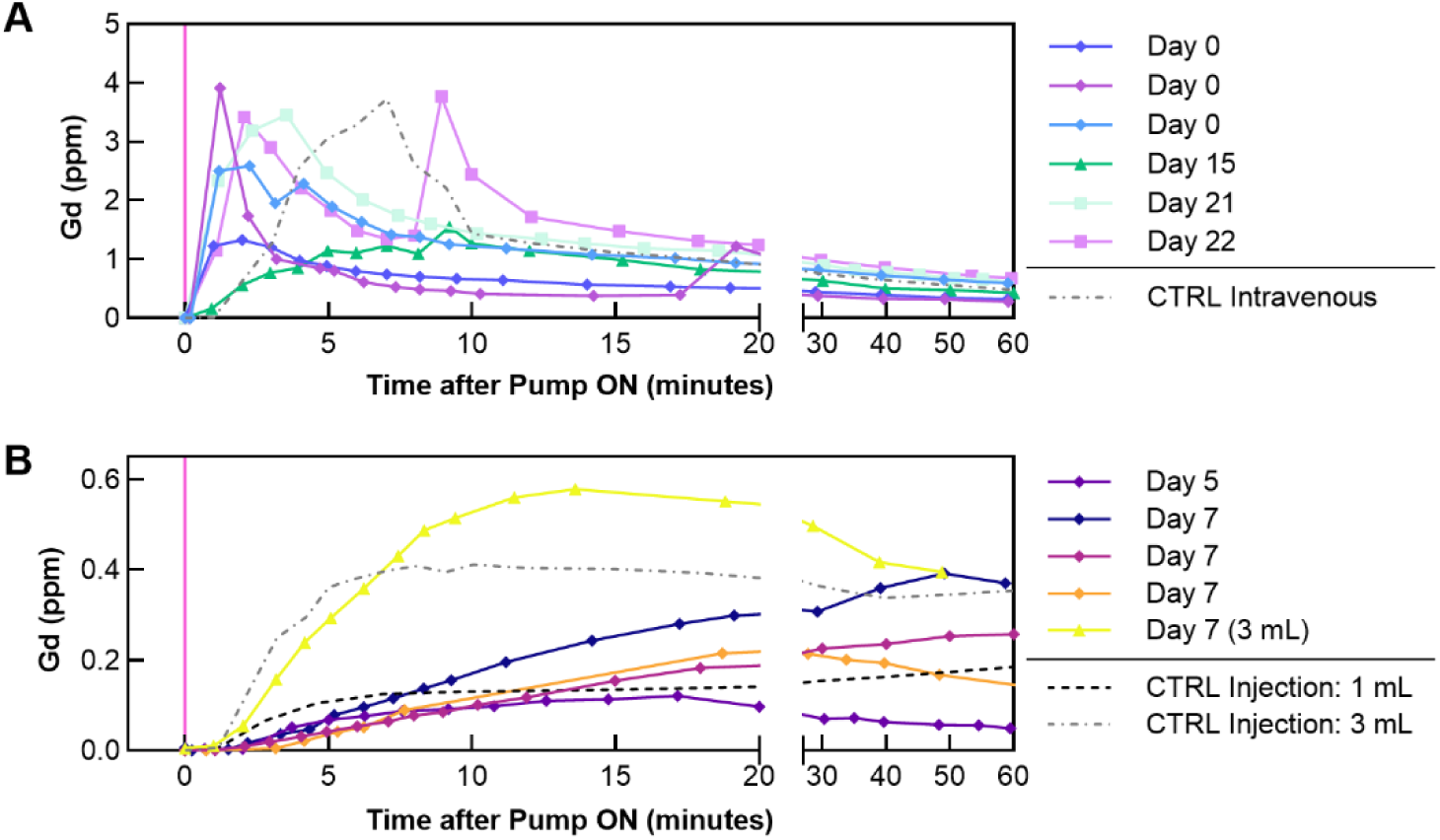
Pharmacokinetics of NLX proxy, Gadolinium (Gd), when delivered by the intravenous Naloximeter after various implantation durations. **(A)** Gadolinium pharmacokinetics when delivered intravenously by the Naloximeter (mammary and jugular vein) between 0 and 22 days after implantation, or a medical infusion pump in the auricular vein (CTRL Intravenous). Dose is 3 mL. **(B)** Gadolinium pharmacokinetics when delivered subcutaneously by the Naloximeter (without a catheter) between 5 and 7 days after implantation, or by manual subcutaneous injection in tissue surrounding healed devices at 21 and 71 days after implantation (CTRL Injection). The subcutaneous dose is 1.5 mL unless otherwise specified.

**Fig. S37.**
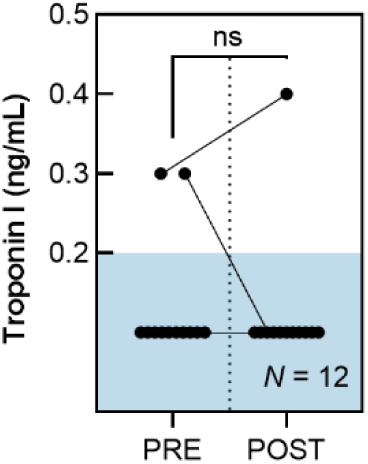
Blood biomarkers for cardiac function before (PRE) and after (POST) intravenous Naloximeter drug delivery. Troponin I quantified from blood serum with a veterinary Troponin I test (IDEXX Laboratories). Values above 0.2 ng/mL are considered elevated, values less than this are plotted within threshold box (lower reporting limit 0.2 ng/mL). Difference is not significant between groups (paired t-test, P = 0.674), though one case showed elevated Troponin levels in both PRE and POST samples.

**Table S1.**
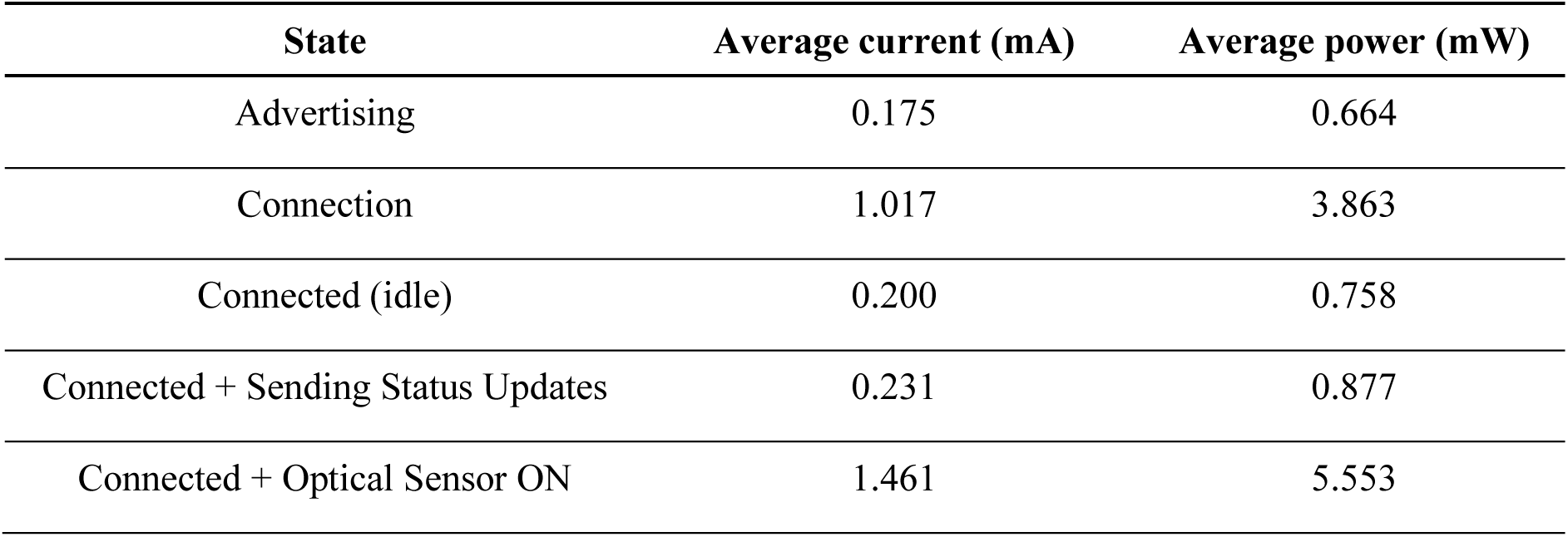
Summary of power consumption of the Naloximeter for large animals during different operational states. Average current and power consumption calculated over a 5 s window.

## Movie S1

**Demonstration of drug delivery from intravenous Naloximeter device.** Pink dye (Rhodamine B) represents NLX solution, clear container of water represents vasculature. Video speed: 1× and 8× (noted in video).

## Movie S2

**Demonstration of drug delivery with injector Naloximeter device, showing dual-injection.** Blue solution indicates NLX solution and yellow solution represents biofluid in the body. Speed: 1×. Scale bar, 1 cm.

