## Supplementary Materials for "An Autonomous Implantable Device for the Prevention of Death from Opioid Overdose"

$$t^* = V^*[G(V^*) + P_0^*] + M^* \frac{V^*}{\frac{d}{dV^*}[V^*G(V^*)] + P_0^*} \quad \ldots \text{(S1)}$$

where $G(V^*)$ is the normalized pressure-volume function describing the deformation of the flexible membrane. Equation S1 can be re-written dimensionally as

$$t = \frac{4F}{3RTi} V[f(V) + P_0] + \frac{32\mu L}{a^4} \frac{V}{\frac{d[Vf(V)]}{dV} + P_0} \quad \ldots \text{(S2)}$$

where $F$ is Faraday's constant, $R$ is the ideal gas constant, $T$ is temperature, $i$ is the electrical current applied to the electrodes, $f(V)$ is the pressure-volume function of the flexible membrane, $\mu$ is the drug viscosity, $L$ is the length of the microfluidic channels, and $a$ is the side length of the channel.

The first term in Eq. S2 quantifies the time required for the membrane to overcome the external pressure and deform into a spherical cap and the second term quantifies the time required for the drug to travel through the microchannels. In this work, microchannels are exchanged for an intravenous catheter, which has characteristic length an order of magnitude larger. In this regime, the microfluidic resistance becomes negligible, and instead the "dead volume" ($V_{DV}$) inside of the catheter tubing becomes significant. Thus, we rewrite Eq. S2 to account for the intravenous device configuration as:

$$t = \frac{4F}{3RTi}(V_d + V_{DV})[f(V_d) + P_0] \quad \ldots \text{(S3)}$$

Where the volume delivered to the bloodstream ($V_d = V - V_{DV} = V - \frac{\pi D^2 L_c}{4}$) is the difference between the volume of gas generated by electrolysis ($V$) and $V_{DV}$ for a given the catheter internal diameter ($D$) and length ($L_c$).

The deformation of the membrane from flat to spherical cap (for finite-deformation) is approximated for a linear-elastic material model as

$$f(V_d) = \frac{64Eh}{3\pi^3 R_0^{10}(1-v)}(V_d + V_{DV})^3 \quad \ldots \text{(S4)}$$

where $E$ is the Young's Modulus, $h$ is the thickness, $v$ is Poisson's ratio, and $R_0$ is the radius of the membrane. A parametric study, shown in Fig. S8, was performed to model the effect of varying the Young's Modulus ($E$ = 1 – 3.5 MPa), thickness ($h$ = 40 –150 μm), and radius ($R_0 = 10 -$

The ODA received packets of data at 1 Hz containing the two raw optical signals, red ($\lambda_1$) and infrared ($\lambda_2$). The ODA operates at this frequency to perform calculations on the current and preceding nine packets of data. A digital low-pass Butterworth filter (0.1 Hz, 4th order) processed the incoming raw data to calculate the $StO_2$ using Eq. 1 and Eq. 2 and the differential optical signal ($\Delta\lambda = \lambda_1 - \lambda_2$). In parallel, a fast Fourier transform (FFT) calculated the power density spectrum on both red and infrared signals filtered using a band-pass Butterworth filter (passband 1-5 Hz, 4th order). The average of the integrated power density spectra provided the power density spectrum index ($PDSi$). Summation of the $PDSi$ over the previous 1 and 10 minutes calculated the short-term and long-term motion indices, $STMi$ and $LTMi$, respectively. Linear interpolation of $\lambda_1$, $StO_2$, and $\Delta\lambda$ followed by differentiation provided the rates of change ($\frac{d}{dt} = \partial$) of the red signal ($\partial\lambda_1$), optical difference ($\partial\Delta\lambda$) and tissue oxygenation ($\partial StO_2$), calculated every 10 seconds over a 60-second window. A comparison of these rates, the $StO_2$ and the $STMi$, against experimentally derived threshold values produced four logical metrics:

$$\begin{bmatrix} \text{M1} \\ \text{M2} \\ \text{M3} \\ \text{M4} \end{bmatrix} = \begin{bmatrix} (\partial\lambda_{1-ThL} < \partial\lambda_1 < \partial\lambda_{1-ThU})\ \&\ (\partial\Delta\lambda_{ThL} < \partial\Delta\lambda < \partial\Delta\lambda_{ThU}) \\ \partial StO_{2-ThL} < \partial StO_2 < \partial StO_{2-ThU} \\ \text{StO}_2 < StO_{2-Th} \\ STMi < STMi_{Th} \end{bmatrix} \quad \dots \text{(S5)}$$

where the subindices in M1 and M2, $ThL$ and $ThU$, correspond to the lower and upper thresholds, respectively, and $StO_{2-Th}$ and $STMi_{Th}$ to the tissue oxygenation and short-term motion index thresholds, respectively.

An OD-relevant event (EVENT) existed when the mutually inclusive condition of the four metrics in Equation S5 in the ODA was prompted, which evolved into a WARNING after $N$ consecutive EVENTs: $N = 3$ if $LTMi > LTMi_{Th}$, corresponding to active/awake state; or $N = 6$ if $LTMi < LTMi_{Th}$, corresponding to rest/sleep state. Upon triggering a WARNING, the ODA prompted the user with a push notification to override the warning, if acknowledged. If one additional EVENT accumulated in the 10 seconds following WARNING without user feedback, the ODA triggered the RESCUE response: activating drug delivery and relaying an emergency call. Figure 3H shows the graphical illustration of the workflow associated with these data analytics.

The following thresholds were used for the metrics:

| | | | | |
|---|---|---|---|---|
| M1 | $\partial\lambda_{1-ThL} = -500$ | $\partial\lambda_{1-ThU} = -10$ | $\partial\Delta\lambda_{ThL} = -500$ | $\partial\Delta\lambda_{ThL} = -10$ |
| M2 | $\partial StO_{2-ThL} = -100$ | $\partial StO_{2-ThU} = -2$ | | |
| M3 | $StO_{2-Th} = 60$ | | | |

| | |
|---|---|
| M4 | $STMi_{Th} = 50$ |

These values were determined as follows. The thresholds for $\partial\lambda_1$, $\partial\Delta\lambda$, $\partial StO_2$ were extracted from data collected during the fentanyl dosage desaturation experiments (e.g. Fig S21 and S25). The 60% threshold for $StO_2$ corresponds to a physiological level that indicates hypoxia and is equivalent to ~ 80% $SpO_2$ (Fig. 2H) or ~ 60% $SO_2$ (Fig. 2I). Finally, the threshold for the $STMi$ is estimated from marked data in human during rest (Fig. S23) and in pigs during rest/sleep (Fig. S24 and S26).

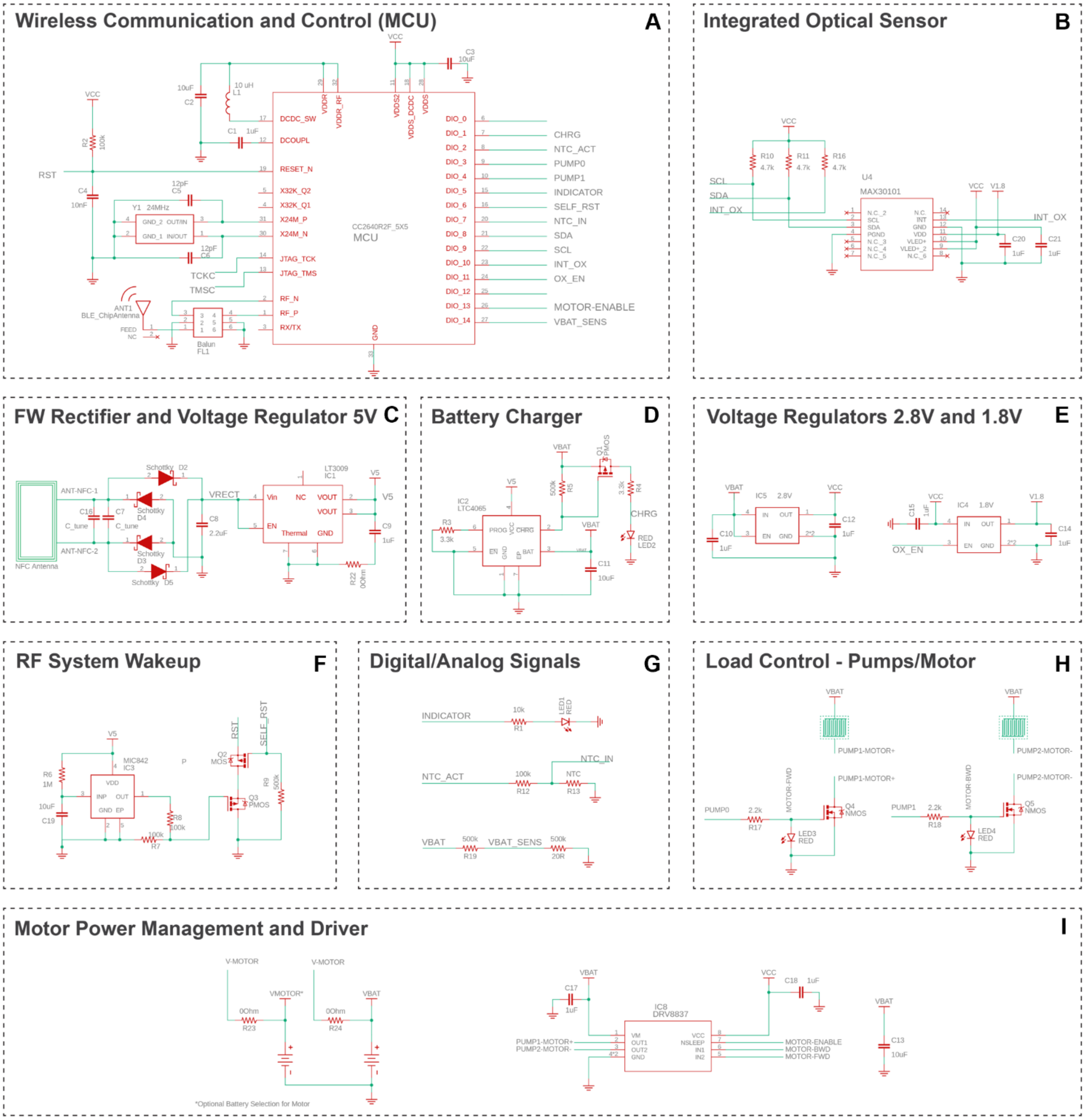


**Fig. S1. Circuit diagrams for subsystems of the Naloximeter platform for large animals. (A)** The CC2640R2F microcontroller (MCU) uses BLE wireless communication and controls the operation of the device. **(B)** The integrated optical sensor MAX30101 connects with the MCU using $I^2C$ serial communication. **(C)** A full-wave (FW) voltage rectifier and regulator converts the harvested input AC to 5 V DC. **(D)** The battery charger is powered by wireless power transfer. **(E)** The step-down voltage regulator supplies 2.8 V to the electronics (left). A 1.8 V regulator provides voltage to the optical sensor only (right). **(F)** An RF actuated wake up system provides a remote reset to the microcontroller in the event of a failed cold start. **(G)** General inputs/outputs control function in the device such as analog battery voltage monitoring, indicator operation, input NTC sensor. **(H)** The power module controls the electrolysis-driven pumps, or the logic signals to operate the H-bridge chip that drives the DC motor. **(I)** The power and control module operates to the DC motor.

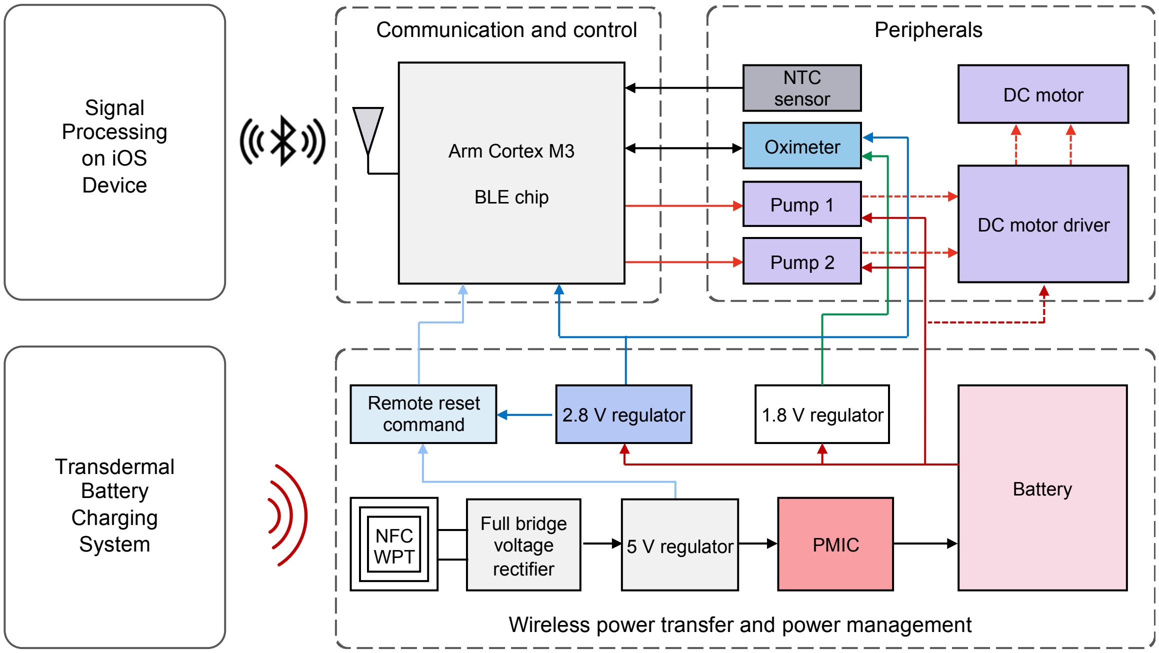


**Fig. S2. Functional block diagram of the Naloximeter platform for large animals.** Modules that construct the electronic system and its interaction with the external iOS device and transdermal battery charging system. The same system, with minor modifications, supports two options for drug delivery, electrolytic pump or DC motor operation.

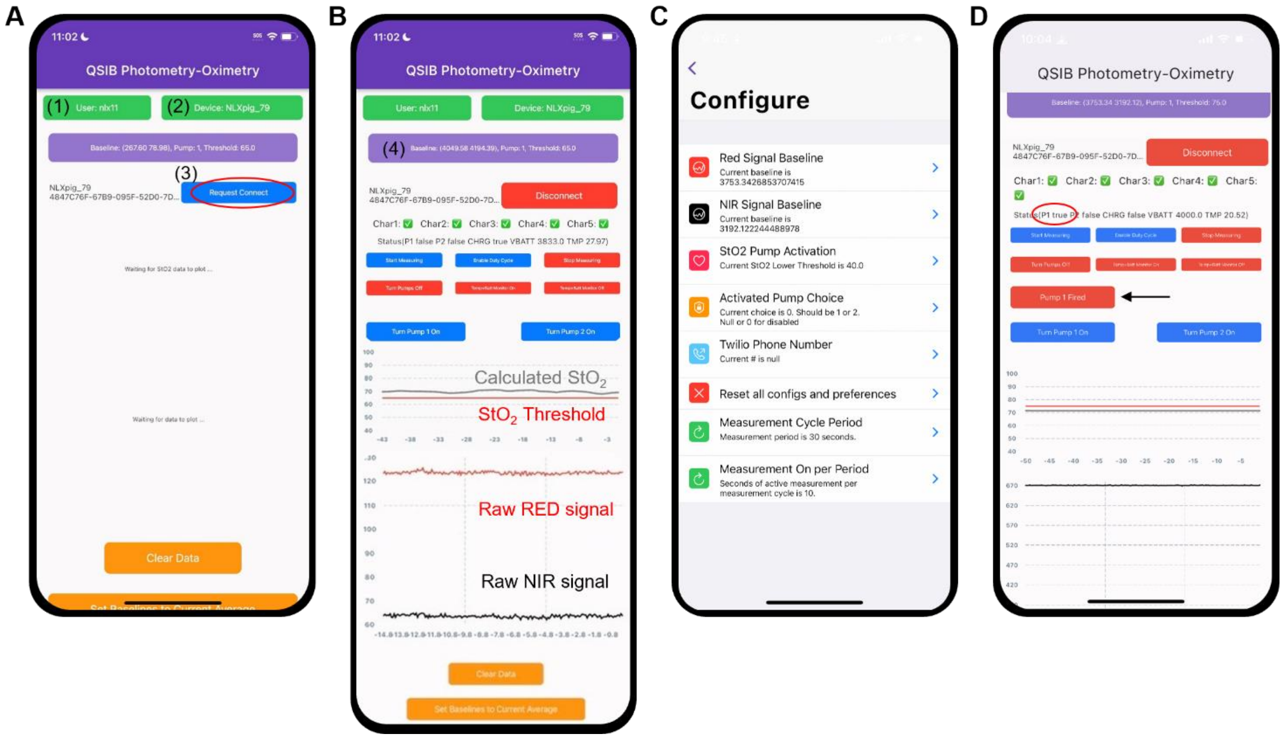


**Fig. S3. Screenshots of the mobile application on a phone. (A)** The main app screen indicates (1) cloud account for data management, (2) device selection, and (3) button for establishing a BLE connection to the device. **(B)** View of the main app screen after establishing device connection, with buttons for controlling device operation, including collection of optical sensor data and calculation of $StO_2$ in real-time. **(C)** Device configuration window, accessed by clicking on (4). **(D)** Main app screen provides two indicators of NLX deployment: pump (P1) status (red circle) and “Pump 1 Fired” display (black arrow).

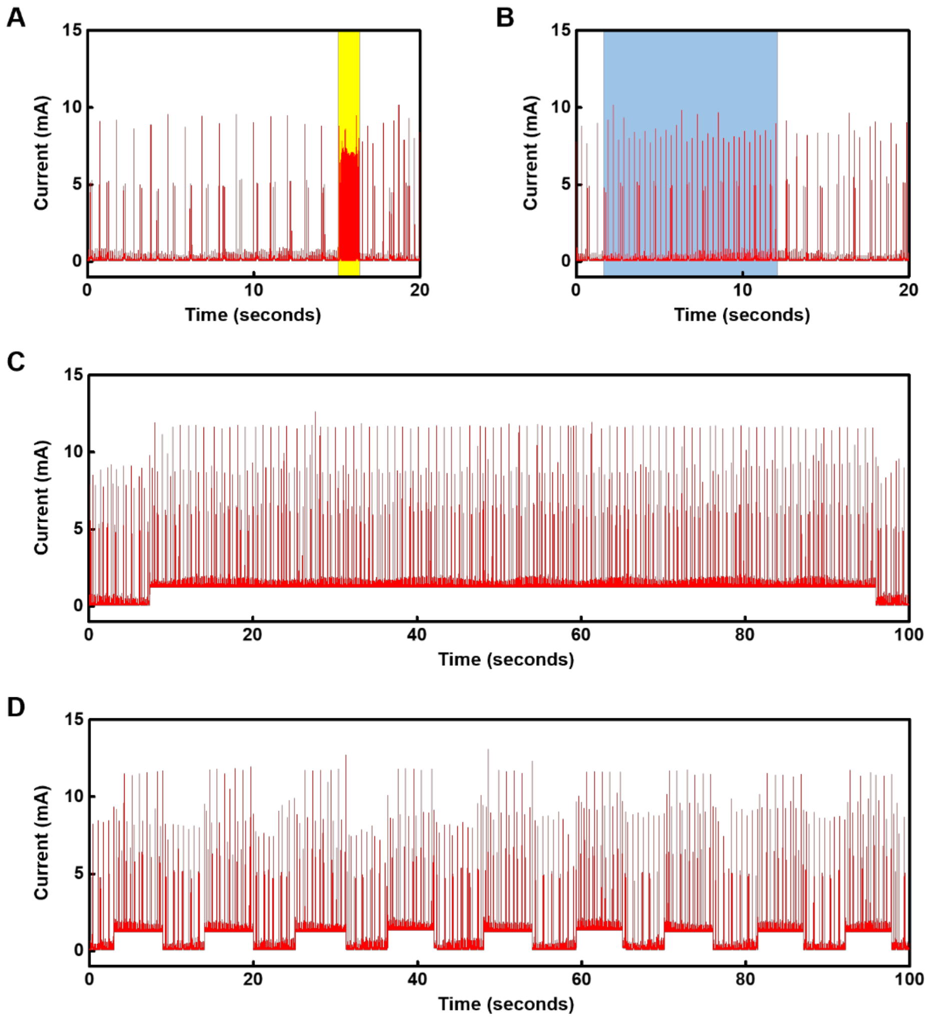


**Fig. S4. Current consumption during operation of a Naloximeter for large animals. (A)** Current consumption of the device before, during (yellow area), and after establishing BLE connection to the peripheral device. **(B)** Current consumption of the device while sending status updates (temperature, battery voltage, pump status, and charging status) at intervals of 1 s (blue area) and while idle. **(C)** Current consumption of the device while operating the dual-color optical sensor in normal mode. **(D)** Current consumption of the device while operating the dual-color optical sensor in intermittent mode (50% duty cycle, 10 s period: 5 s ON, 5 s OFF). Supply voltage is 3.8 V in all cases.

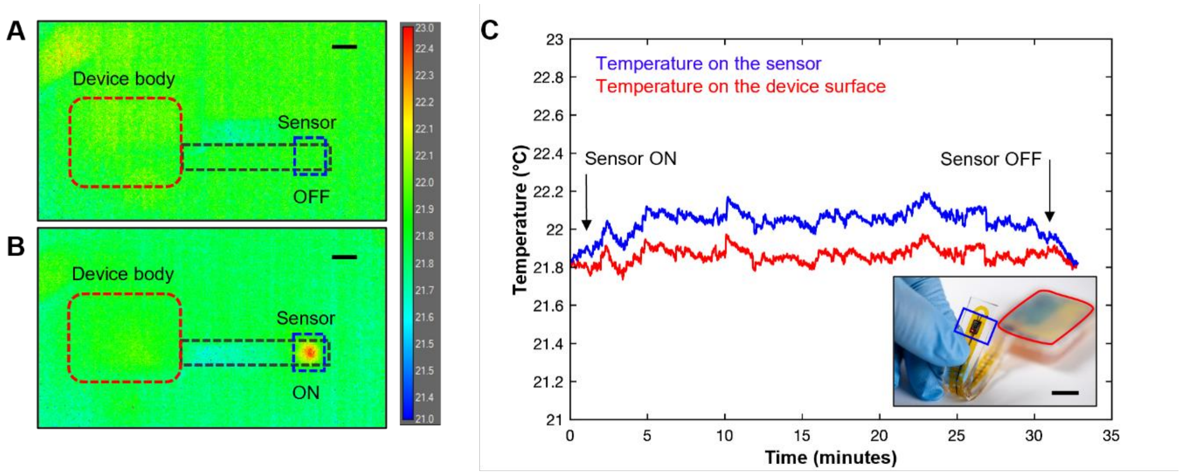


**Fig. S5. Benchtop characterization of thermal load produced by the dual-wavelength optical sensor during operation.** Thermographic images of an encapsulated test of a device with **(A)** the sensor OFF and **(B)** sensor ON at room temperature. Scale bars, 1 cm. **(C)** Surface average temperature recorded in the area above the electronic board and on the sensor. Inset: optical image of the device used in this characterization study, with sensor separated from the body of the device. Scale bar, 1 cm.

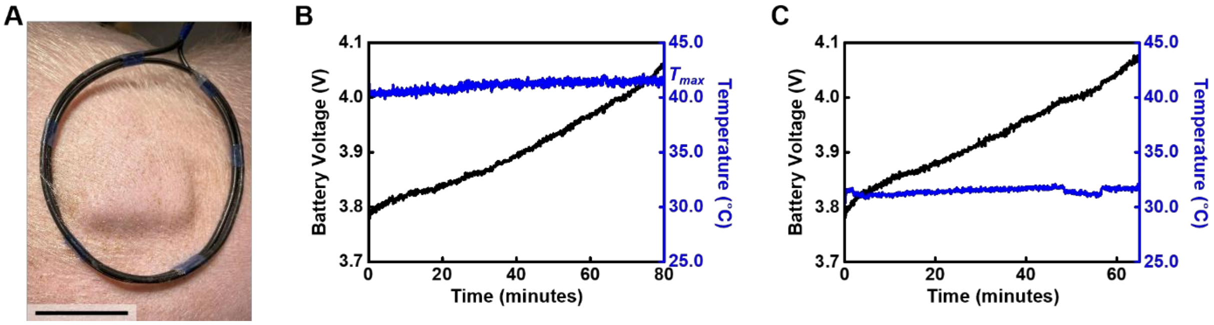


**Fig. S6. Transdermal charging of a Naloximeter device for large animals. (A)** Photograph showing wireless transdermal charging of a device implanted in a pig model after healing. Scale bar, 5 cm. **(B)** Battery voltage and temperature measured by the NTC in the device during transdermal charging, maximum temperature labeled as $T_{max}$. **(C)** Battery voltage and temperature measured by the NTC in the device during charging on the benchtop.

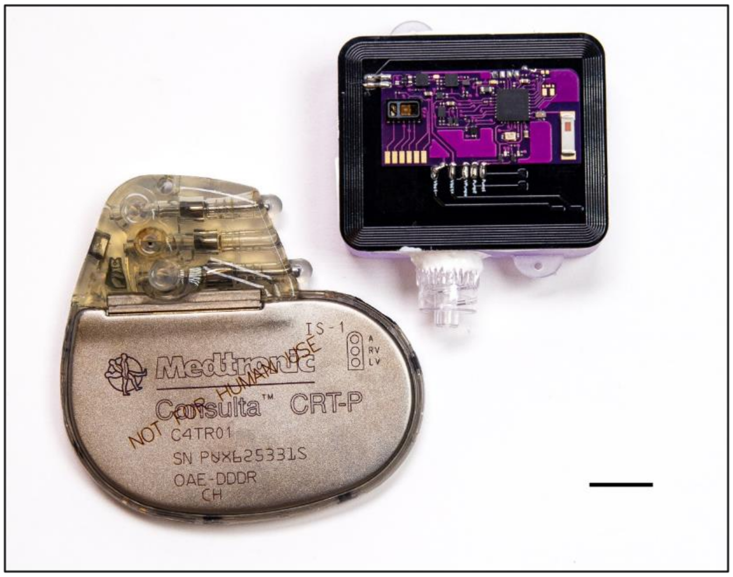


**Fig. S7. Optical image of a Naloximeter and a commercial pacemaker (Consulta, Medtronic).** Scale bar is 1 cm.

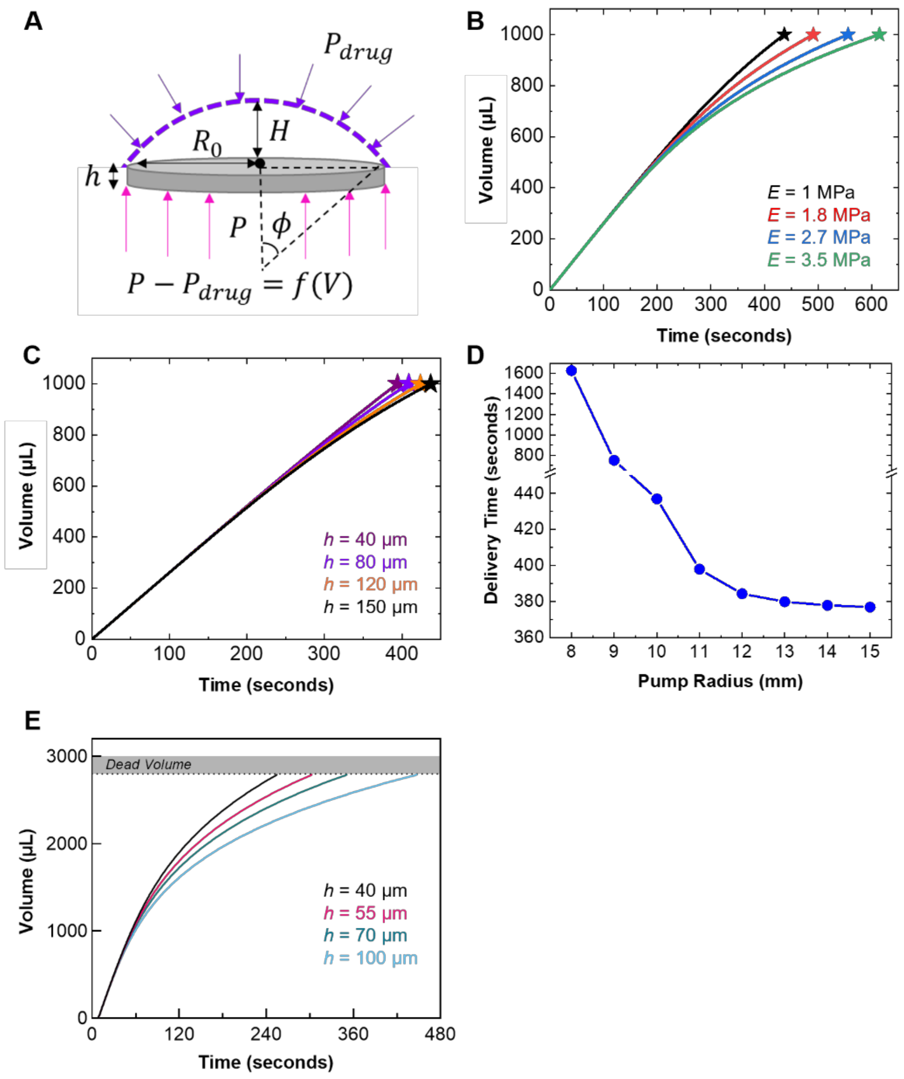


**Fig. S8. Parametric FEA modeling of the operation of electrolytic pumps. (A)** Schematic diagram of forces on the membrane relevant to these simulations. **(B)** Volume vs. time curves for 1 mL pumps varying the modulus of elastic membrane ($h$ = 1 µm). **(C)** Volume vs. time curves for 1 mL pumps varying the membrane thickness ($E$ = 1 MPa). **(D)** Effect of pump radius on delivery time for 1 mL drug volume, ($E$ = 4.3 MPa and $h$ = 70 µm). Calculations in (A – D) run with constants: $i$ = 15 mA, T = 310 K. **(E)** Evaluation of the effects of membrane thickness for 3 mL pumps using parameters measured in actual devices, $i$ = 120 mA and $E$ = 4.3 MPa.

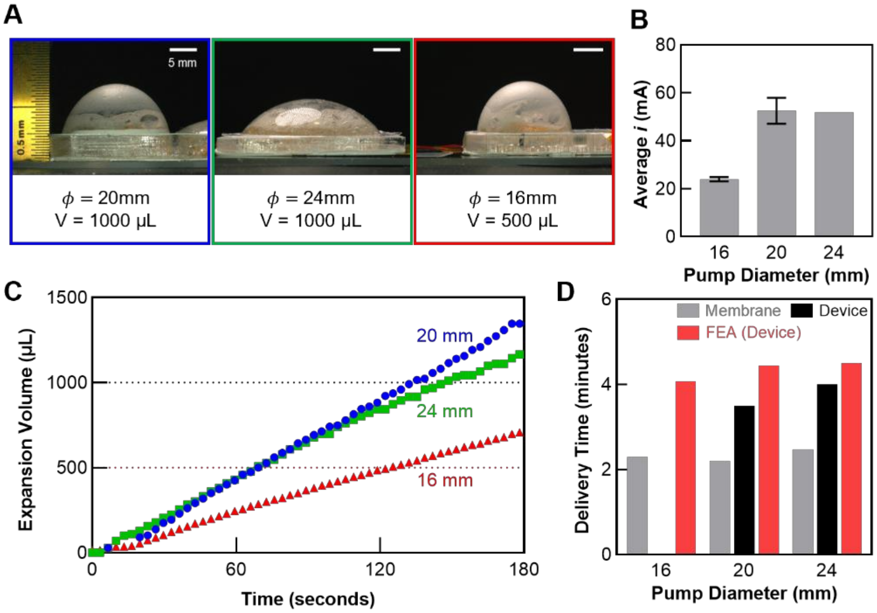


**Fig. S9. Benchtop studies of the effects of pump geometry on the profile of membrane deformation. (A)** Photograph of three electrolytic pumps with different geometries at inflated states. Scale bars, 5 mm. **(B)** Average current consumption for pumps with different diameters, recorded during electrolysis with the device PCB and battery. **(C)** Expansion volume calculated by image analysis as the membrane deforms during electrolysis. **(D)** Estimated time to deliver the total drug reservoir content according to benchtop studies of the membrane deformation, full device testing, and FEA modeling ($E$ = 4.3 MPa, $h$ = 70 µm, $T$ = 310 K, and $i$ from experiment).

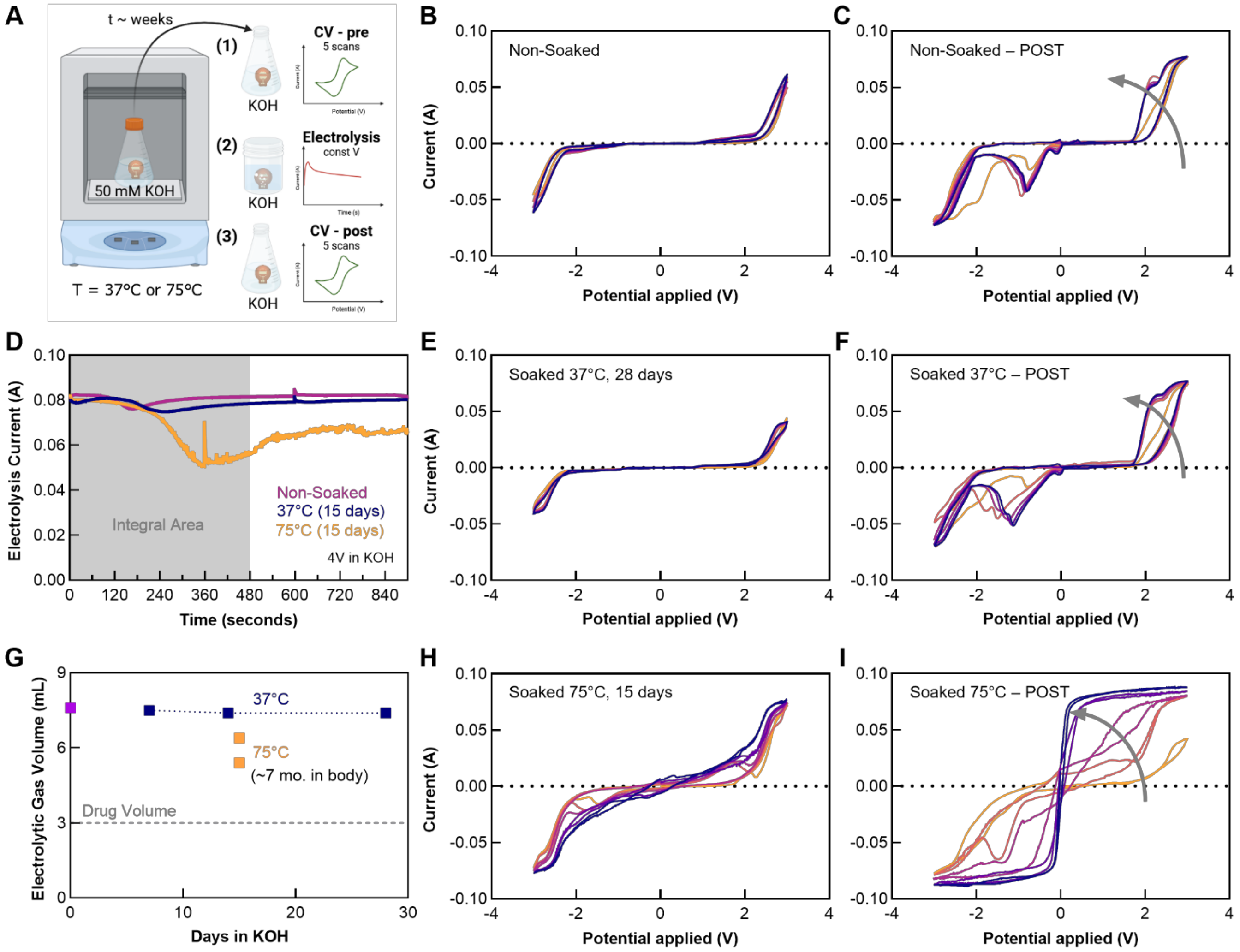


**Fig. S10. Benchtop studies of electrode degradation. (A)** Schematic illustration of the experimental setup and study design. The experiments consisted of soaking in potassium hydroxide electrolyte (50 mM KOH) at elevated temperature (37°C or 75°C) for several weeks followed by *(1)* cyclic voltammetry (CV), *(2)* constant voltage electrolysis, and *(3)* post-electrolysis CV. All CV tests involved scans of voltage between +/- 3 V at a rate of 50 mV/sec for 5 cycles. Constant volage electrolysis testing at an applied potential of 4 V. All electrochemical characterization was carried out in the electrolyte solution, 50 mM KOH. Cyclic voltammetry plots for a fresh non-soaked control electrode **(B)** before and **(C)** after electrolysis. (D) Electrolysis current at constant voltage for control and aged electrodes. Cyclic voltammetry plots for an electrode aged at 37°C for 28 days (E) before and **(F)** after electrolysis. **(G)** Volume of gas generated by electrolysis, as calculated with Faraday's Law and measured current (panel D) plotted according to aging time and temperature. Cyclic voltammetry plots for an electrode aged at 75°C for 15 days – equivalent to 7 months at body temperature, **(H)** before and **(I)** after electrolysis.

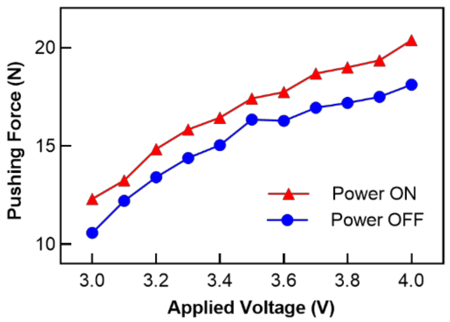


**Fig. S11. Force supplied by the plunger and DC motor as a function of applied voltage.** Each data point corresponds to the saturated force with the maximum load on the motor.

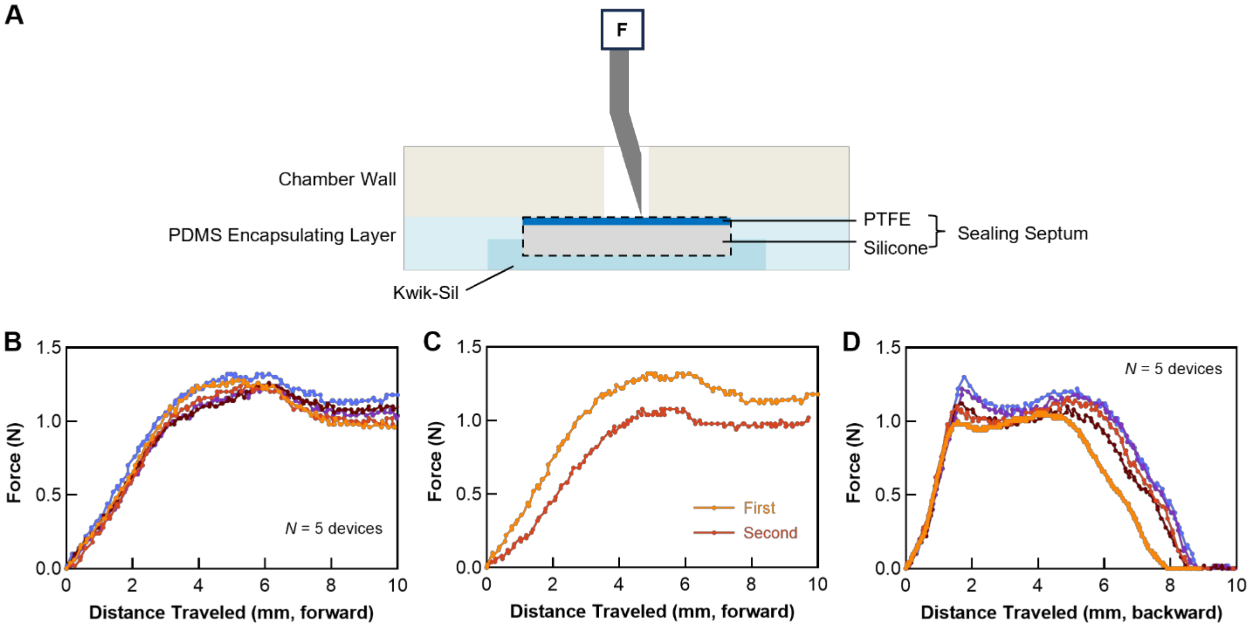


**Fig. S12. Force to deploy and retract the Huber needle through the sealing septum. (A)** Schematic illustration of the setup to measure the force to deploy and retract the needle through the sealing septum. **(B)** Required force for the needle to pierce the front sealing septum. **(C)** Required force for the needle to pierce the front sealing septum for the first and second time. **(D)** Required force to retract the deployed needle through the sealing septum.

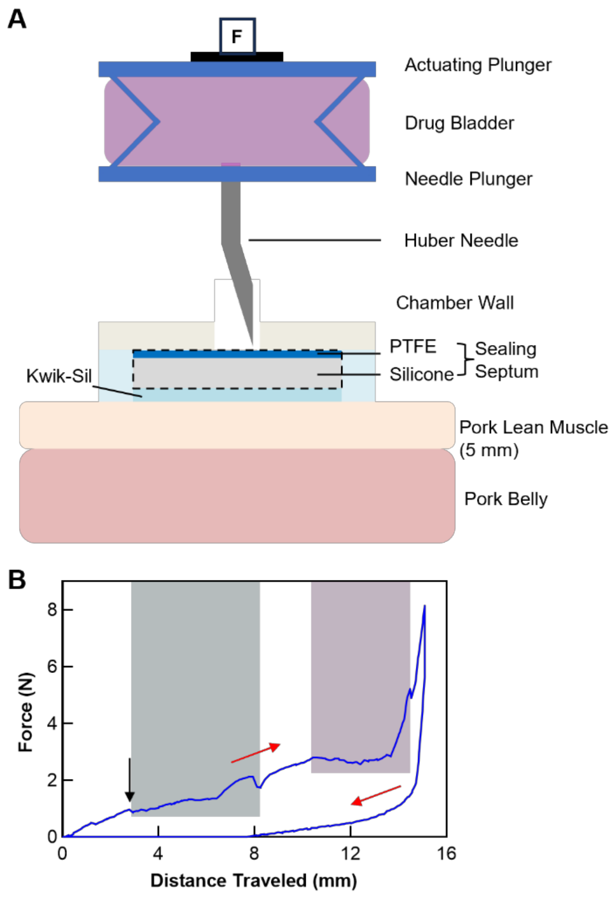


**Fig. S13. Force during a successive sequence of deploying the needle, injecting the solution, and retracting the needle. (A)** Schematic image of the experimental setup. The bladder is filled with 1.5 mL of solution for this characterization. **(B)** Plot of force during a successive sequence of deploying the needle into a two-layer pork tissue model, injecting the solution, and retracting the needle. Red arrows indicate the progression of time. The black arrow indicates completion of the process of piercing the sealing septum. The blue area corresponds to the regime for piercing two layers of tissue. The pink area corresponds to the regime for injecting the solution from the bladder into the pork belly.

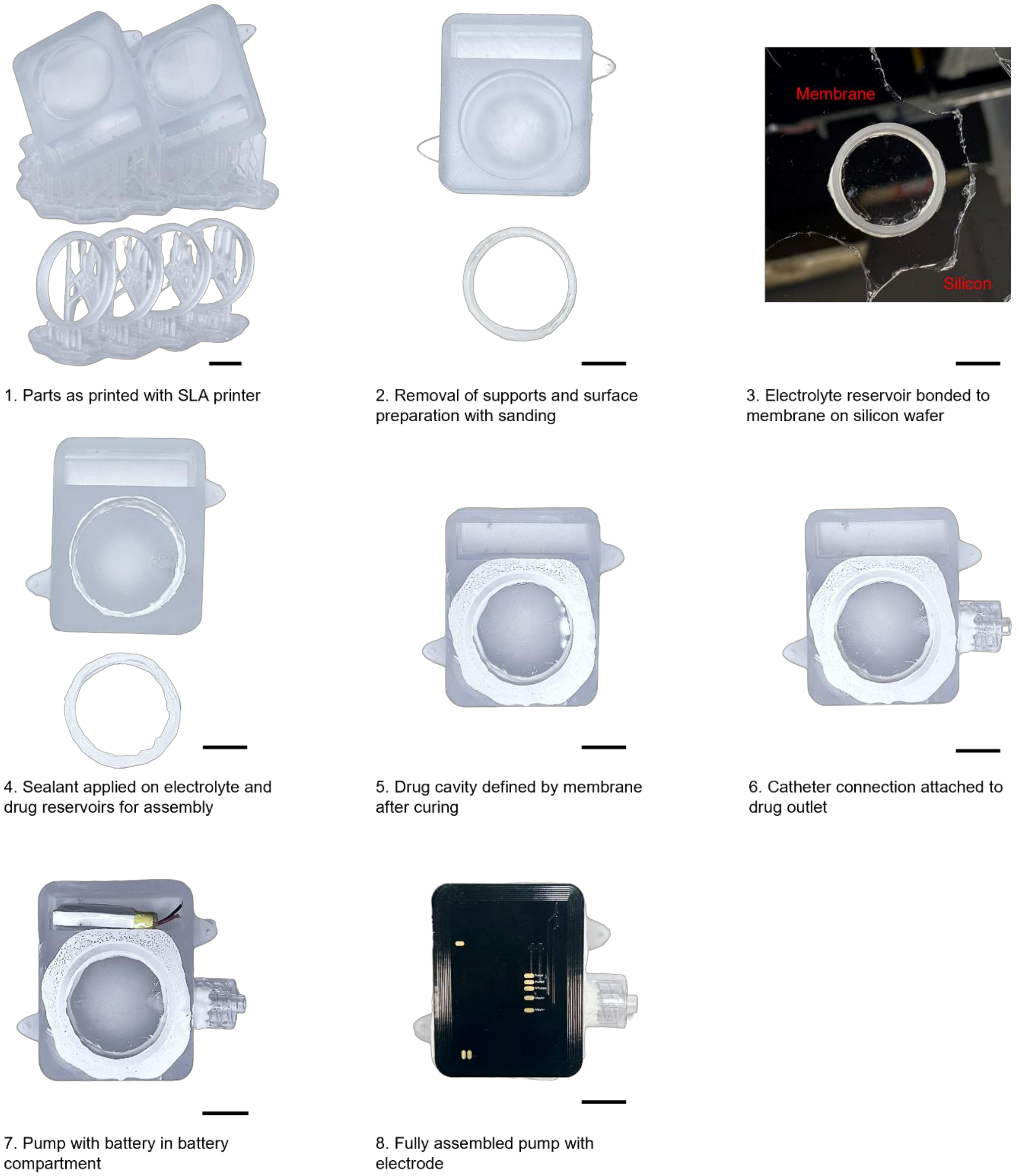


**Fig. S14. Steps for fabricating intravenous Naloximeter devices.** Scale bars, 1 cm.

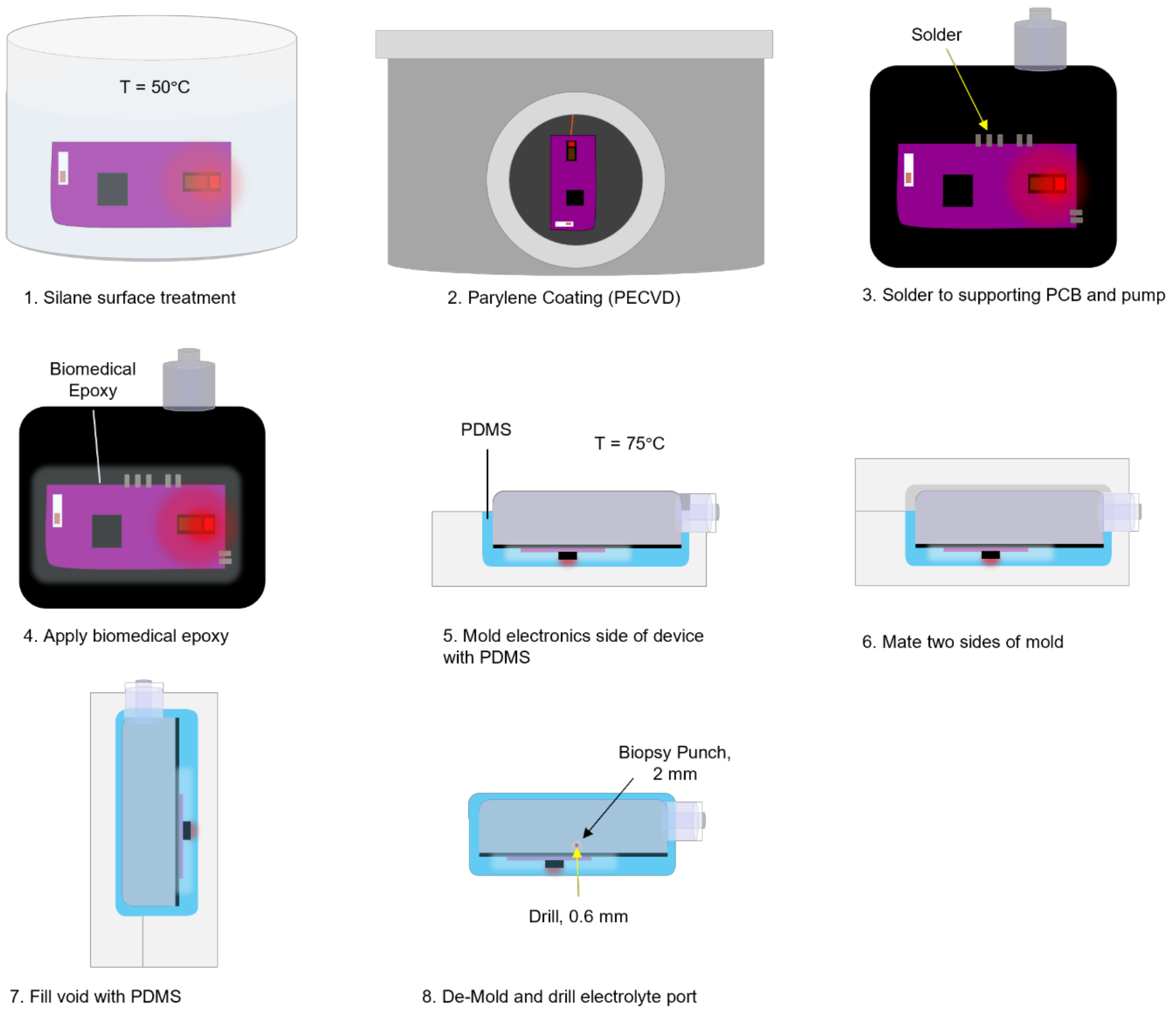


**Fig. S15. Steps for encapsulating Naloximeter devices, illustrated with the intravenous device.**

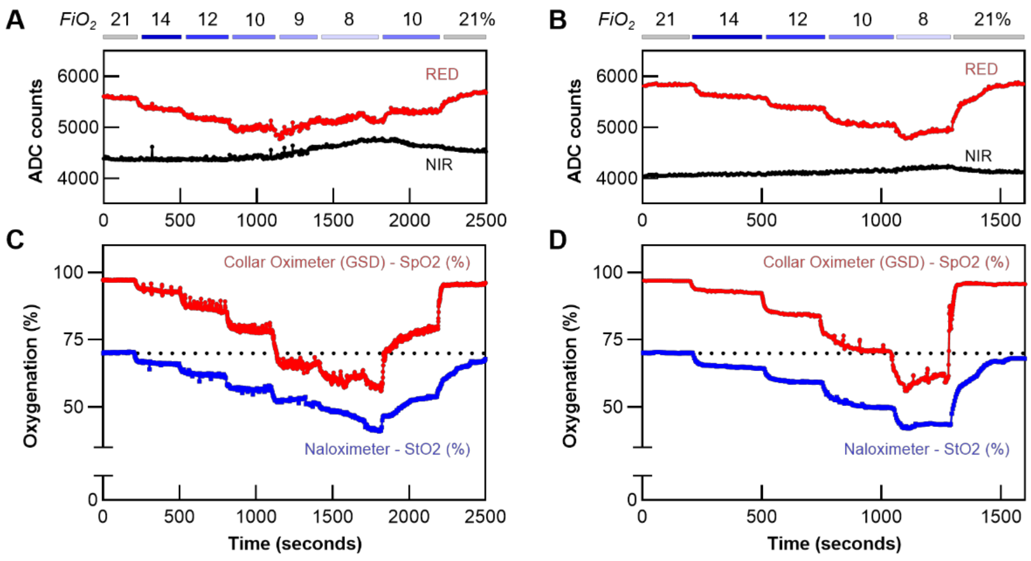


**Fig. S16. Hypoxia studies in rodent models. (A, B)** Raw red and near-infrared (NIR) optical signals from an NFC Naloximeter during hypoxia induced by modulation of the fraction of inspired oxygen ($FiO_2$). **(C, D)** Corresponding calculated $StO_2$ from the Naloximeter and $SpO_2$ measured by a gold standard device (GSD, collar oximeter). Each column represents one animal.

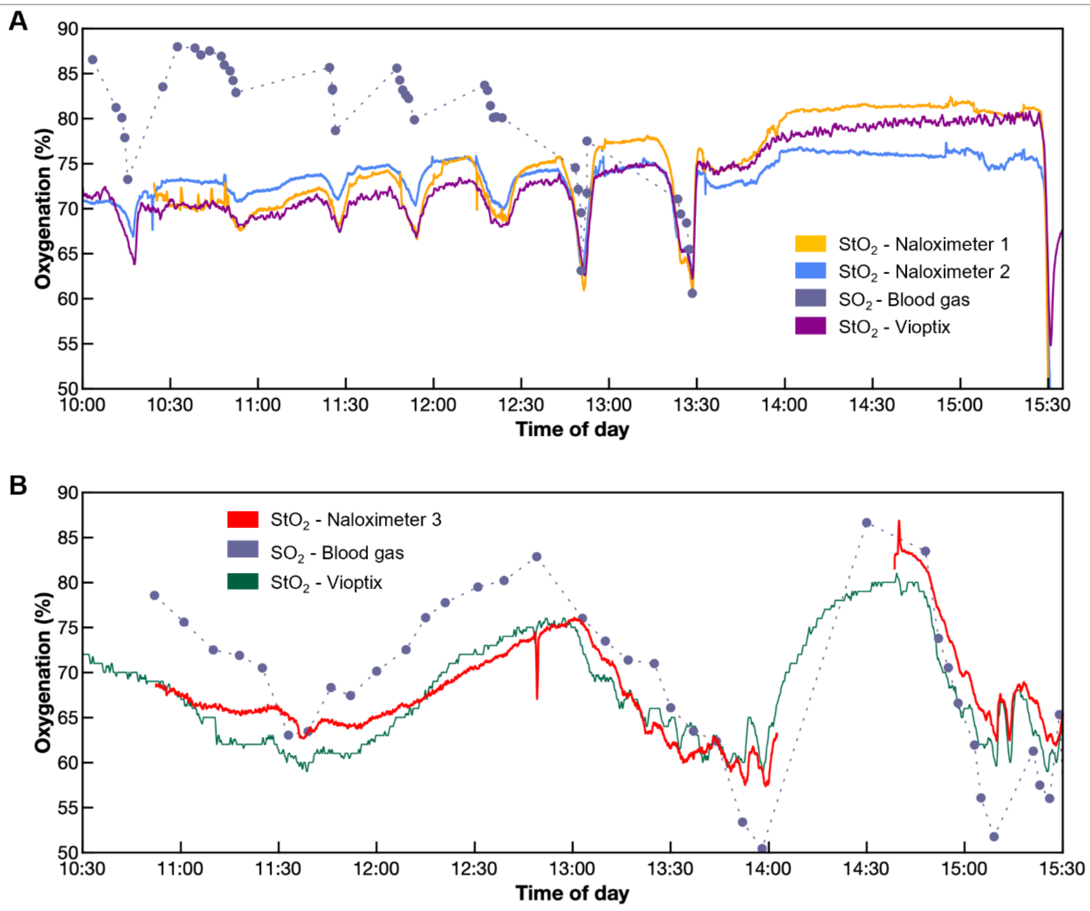


**Fig. S17. Hypoxia studies in a porcine model with comparison to standard clinical devices and measurements. (A, B)** Blood gas saturation is calculated as a mixture of 70% venous and 30% arterial blood.

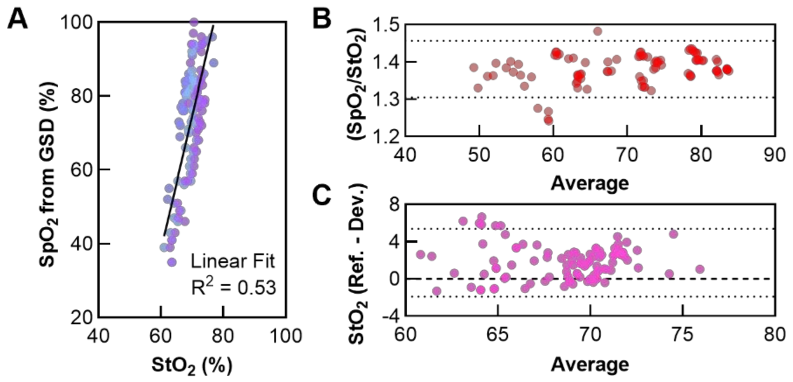


**Fig. S18. Extended analyses of hypoxia studies in swine and rodent models. (A)** Comparison of $SpO_2$ measured with gold standard device (GSD, clip-on pulse oximeter) and $StO_2$ measured with a Naloximeter during hypoxia in a porcine model. **(B)** Bland-Altman plot to compare (ratio versus average) $SpO_2$ measured with gold standard device and $StO_2$ measured with the Naloximeter. Dashed lines depict the 95% confidence interval. **(C)** Bland-Altman plot to compare (difference versus average) $StO_2$ measured with a clinical device (Vioptix) and $StO_2$ measured with the Naloximeter in porcine model. Dashed lines depict the 95% confidence interval.

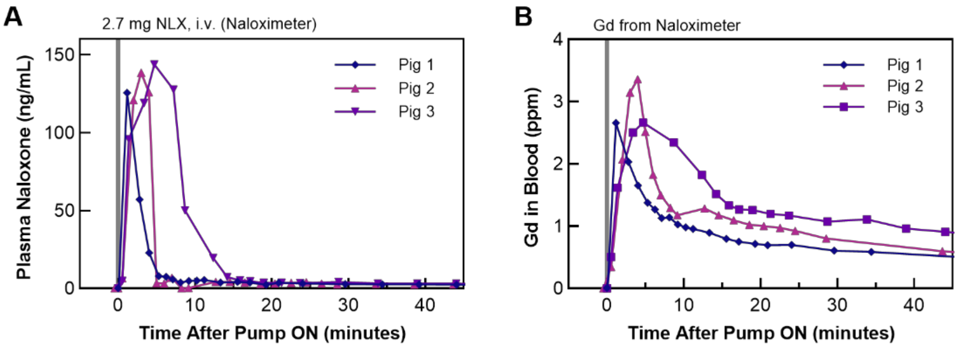


**Fig. S19. Gadolinium (Gd) and Naloxone pharmacokinetics in porcine models with intravenous Naloximeter devices. (A)** Naloxone and **(B)** Gd measured in plasma and blood, respectively, after delivery of 3 mL of drug solution from Naloximeter device. $N = 3$ animals, the same blood draw was split into samples for NLX and Gd quantification to facilitate direct comparison between the methods.

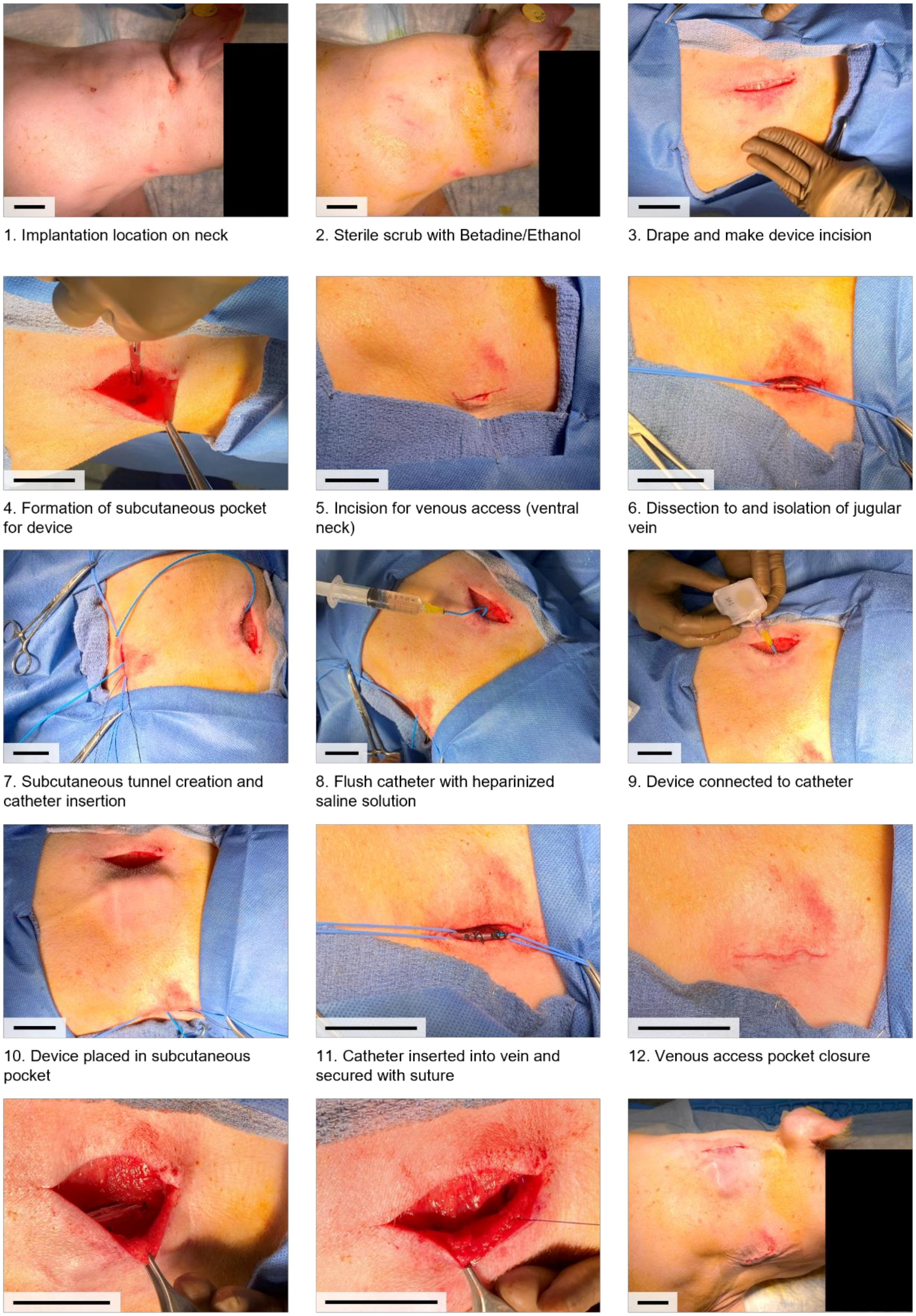

1. Implantation location on neck
2. Sterile scrub with Betadine/Ethanol
3. Drape and make device incision
4. Formation of subcutaneous pocket for device
5. Incision for venous access (ventral neck)
6. Dissection to and isolation of jugular vein
7. Subcutaneous tunnel creation and catheter insertion
8. Flush catheter with heparinized saline solution
9. Device connected to catheter
10. Device placed in subcutaneous pocket
11. Catheter inserted into vein and secured with suture
12. Venous access pocket closure
13. Device secured in place using its suture fixation wing
14. Device pocket closure in multiple layers with suture
15. Post-operative incision care: Film dressing (Tegaderm)

**Fig. S20. Surgical approach for implanting an intravenous Naloximeter device in the external jugular vein of a pig.** Scale bars, 5 cm.

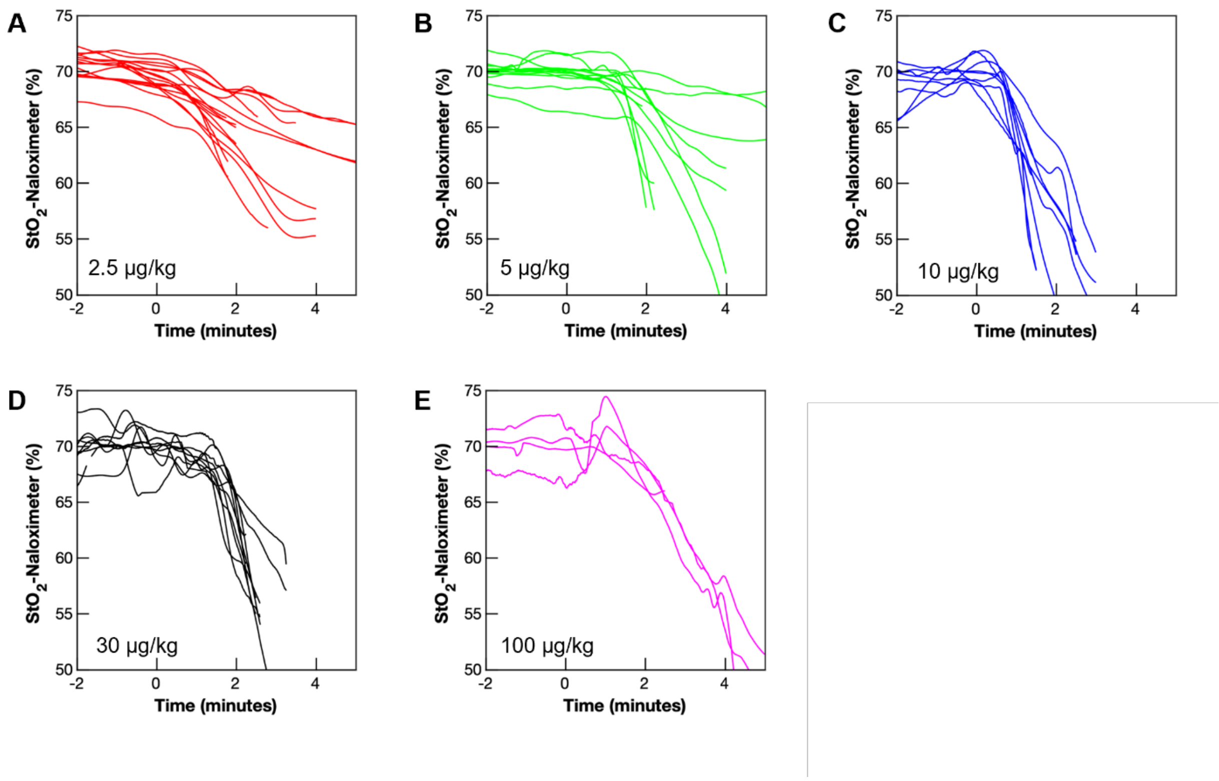


**Fig. S21. Tissue oxygenation ($StO_2$) recorded with Naloximeters in porcine models at different fentanyl doses. (A)** Fentanyl dose of 2.5 µg/kg, $N = 6$ subjects and $n = 17$ devices. **(B)** Fentanyl dose of 5 µg/kg, $N = 4$ subjects and $n = 12$ devices. **(C)** Fentanyl dose of 10 µg/kg, $N = 9$ subjects and $n = 9$ devices. **(D)** Fentanyl dose of 30 µg/kg, $N = 6$ subjects and $n = 10$ devices. **(E)** Fentanyl dose of 100 µg/kg, $N = 4$ subjects and $n = 4$ devices. Fentanyl was administered at time = 0. Overall $N = 29$ subjects, $n = 52$ devices.

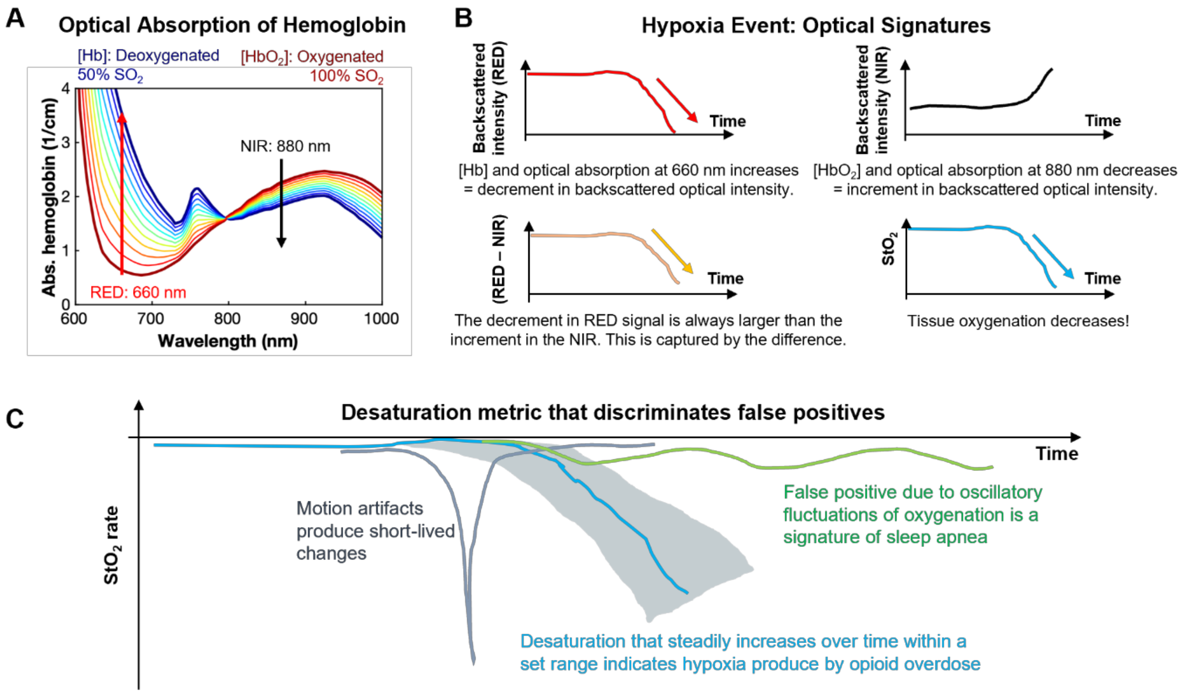


**Fig. S22. Physiological dynamics during hypoxia produce measurable signatures that contrast with false positives produced by motion artifacts and sleep apneas. (A)** Optical absorption of hemoglobin depends on the concentration of its oxygenated ($HbO_2$) and deoxygenated (Hb) states. Based on data from ref *(50)*. Graph shows the change in the optical absorption as hemoglobin oxygenation changes from 100% $SO_2$ (dark red) to 50% $SO_2$ (dark blue). Differential spectroscopy, at the red = 660 nm ($\boldsymbol{\lambda_1}$) and near-infrared (NIR) = 880 nm ($\boldsymbol{\lambda_2}$) wavelengths, is used to calculate the tissue oxygenation ($\boldsymbol{StO_2}$). These signals capture the physiological dynamics. **(B)** Hypoxia is accompanied by clear changes in the optical signal and the calculated $StO_2$. These data produce a set of variables to detect desaturation. **(C)** Typical confounding events follow from motion artifacts and behaviors associated with sleep apnea. Motion artifacts can lead to false positives that are typically eliminated based on criteria associated with the rate of change of the optical signal. Episodes of desaturation due to sleep apnea can be eliminated by their characteristic oscillatory patterns and desaturation rate.

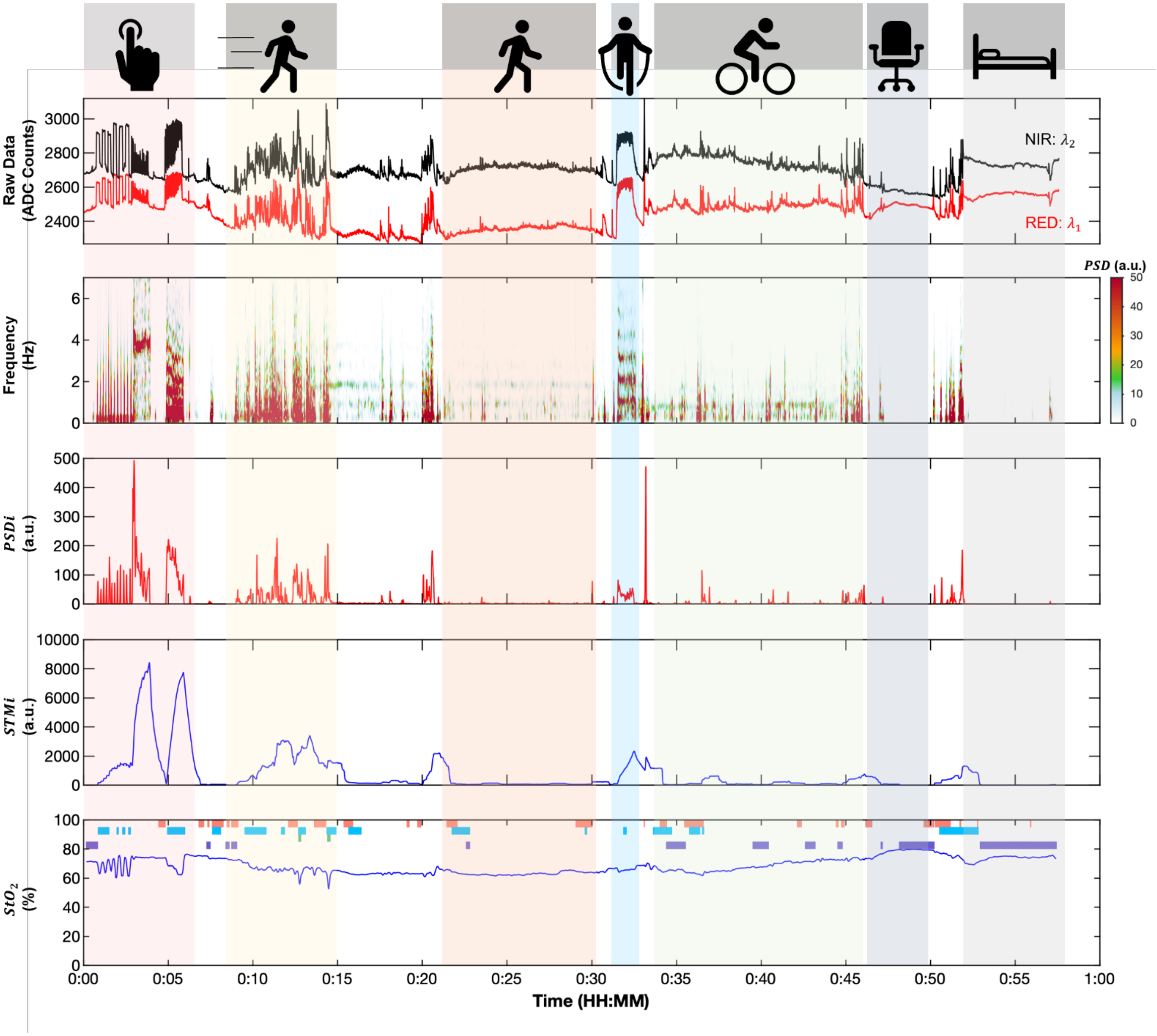


**Fig. S23. Data from cutaneous optical sensors in humans associated with different types of physical activity.** Data recorded with dual-wavelength optical sensor mounted on the skin in the arm of a human subject volunteer performing a series of movements. Movement sequence: Applying pressure to the skin (red), running (yellow), walking (orange), aerobic jumping (blue), cycling (green), resting while seated (blue-gray), resting while supinated (gray). Plots from top-down: red and near-infrared (NIR) optical signals; spectrogram; power spectral density index ($\boldsymbol{PSDi}$) in the 1-5 Hz frequency band; short-term motion index ($\boldsymbol{STMi}$, 60 s) and calculated tissue oxygenation ($\boldsymbol{StO_2}$).

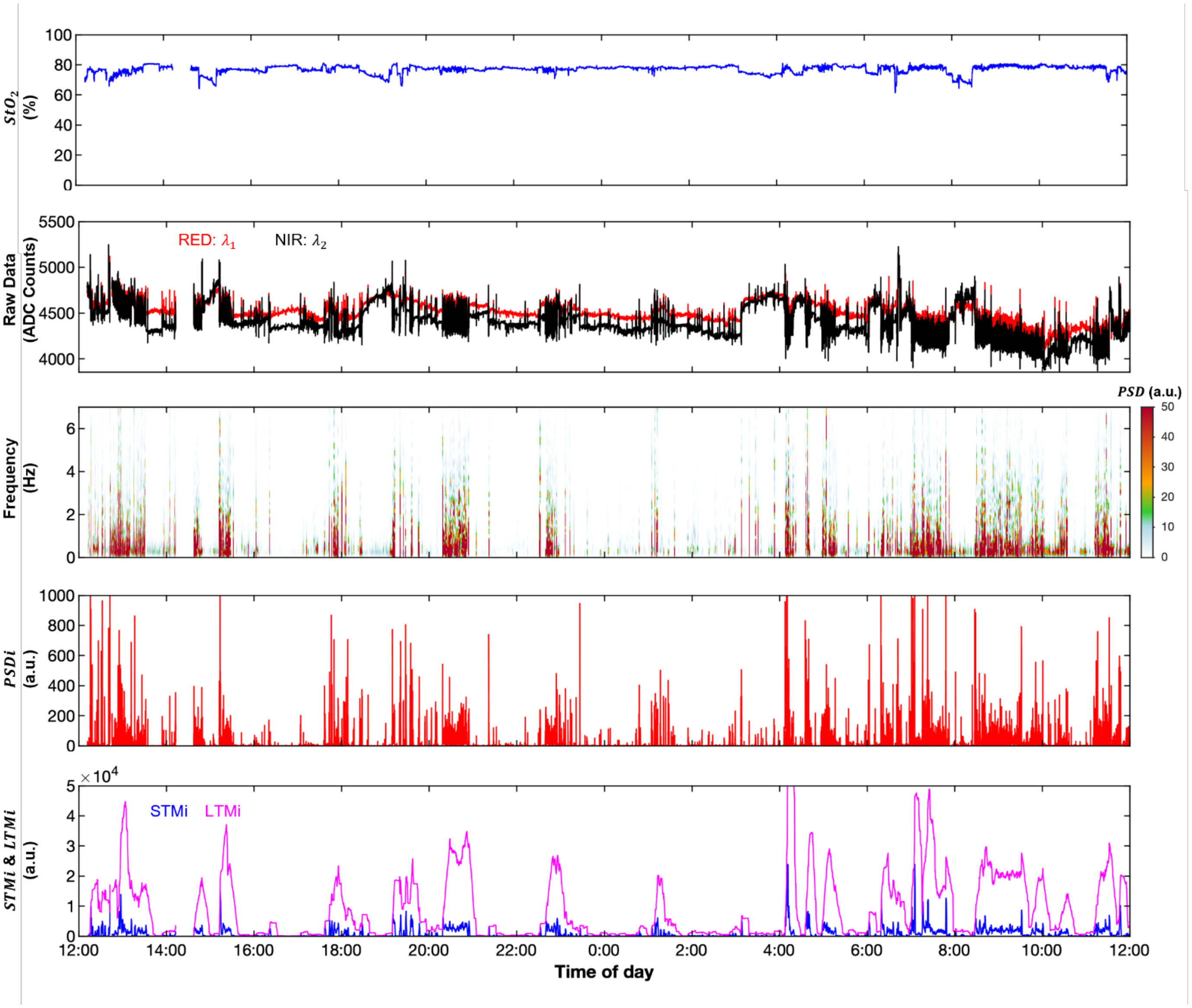


**Fig. S24. Cumulative power spectral density index for defining episodes of physical activity.** Data recorded with dual-wavelength optical sensor in a freely moving porcine model over 24 hours. From top-down: calculated tissue oxygenation ($\boldsymbol{StO_2}$); red and near-infrared (NIR) raw optical signals; spectrogram; power spectral density index ($\boldsymbol{PSDi}$) in the 1-5 Hz frequency band; short-term motion index ($\boldsymbol{STMi}$, 60 s) and long-term motion index ($\boldsymbol{LTMi}$, 10 min). Episodes of high motion index, apparent in the raw data, corresponds with the episode of increased $\boldsymbol{LTMi}$, power spectral density and $\boldsymbol{PSDi}$.

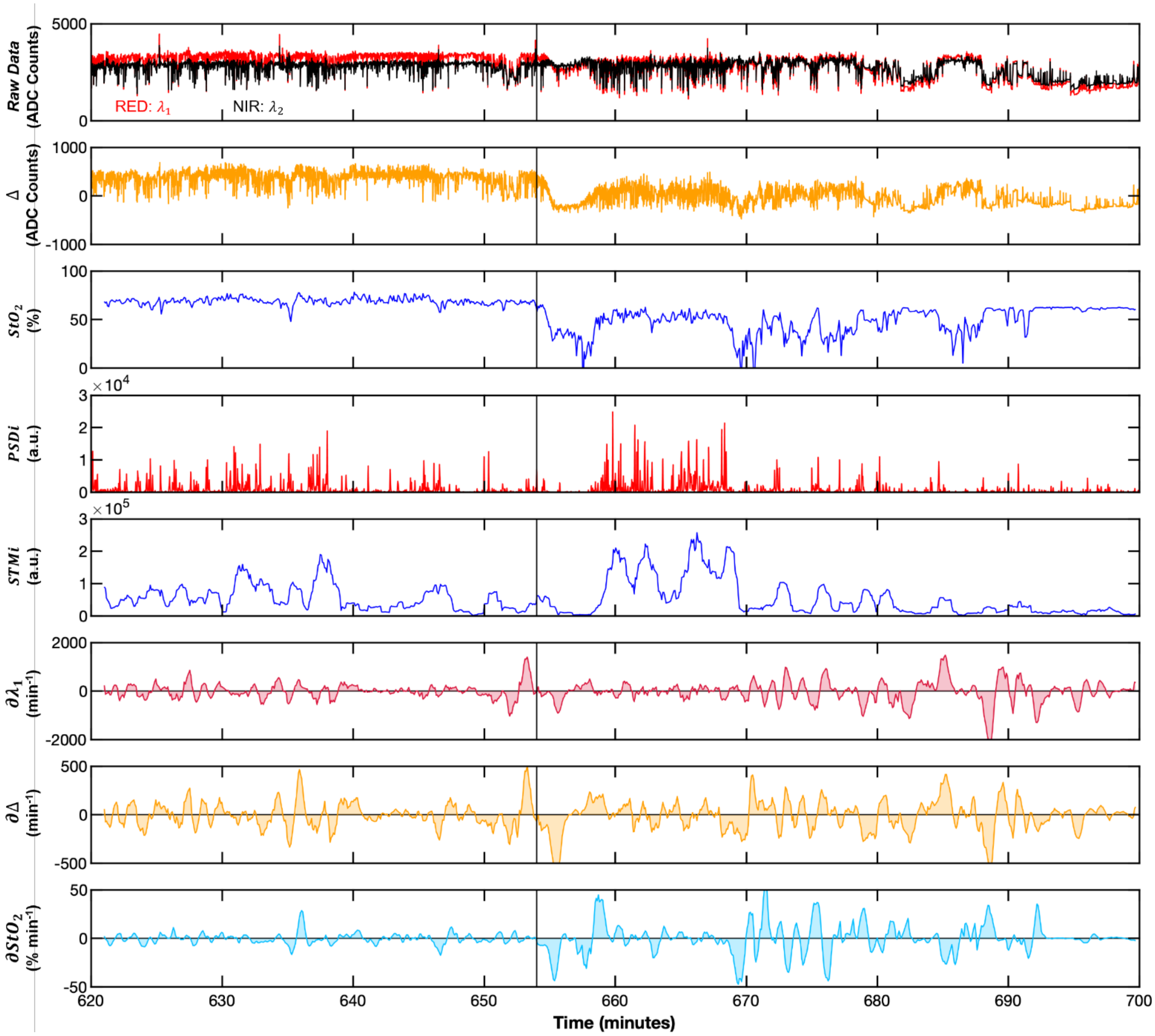


**Fig. S25. Optical sensor data, and its multivariable derivatives, during fentanyl overdose in a freely moving pig.** From top-down: raw data of the red (RED, $\boldsymbol{\lambda_1}$) and near-infrared (NIR, $\boldsymbol{\lambda_2}$) optical signals; difference between these signals ($\Delta = \boldsymbol{\lambda_1} - \boldsymbol{\lambda_2}$); calculated tissue oxygenation ($\boldsymbol{StO_2}$); power spectral density index ($\boldsymbol{PSDi}$) in the 1–5 Hz frequency band; short-term motion index ($\boldsymbol{STMi}$); rate of change ($\boldsymbol{\partial}$) of red ($\boldsymbol{\partial\lambda_1}$); difference ($\boldsymbol{\partial}\Delta$); and tissue oxygenation ($\boldsymbol{\partial StO_2}$). The time of fentanyl administration is indicated with a vertical line.

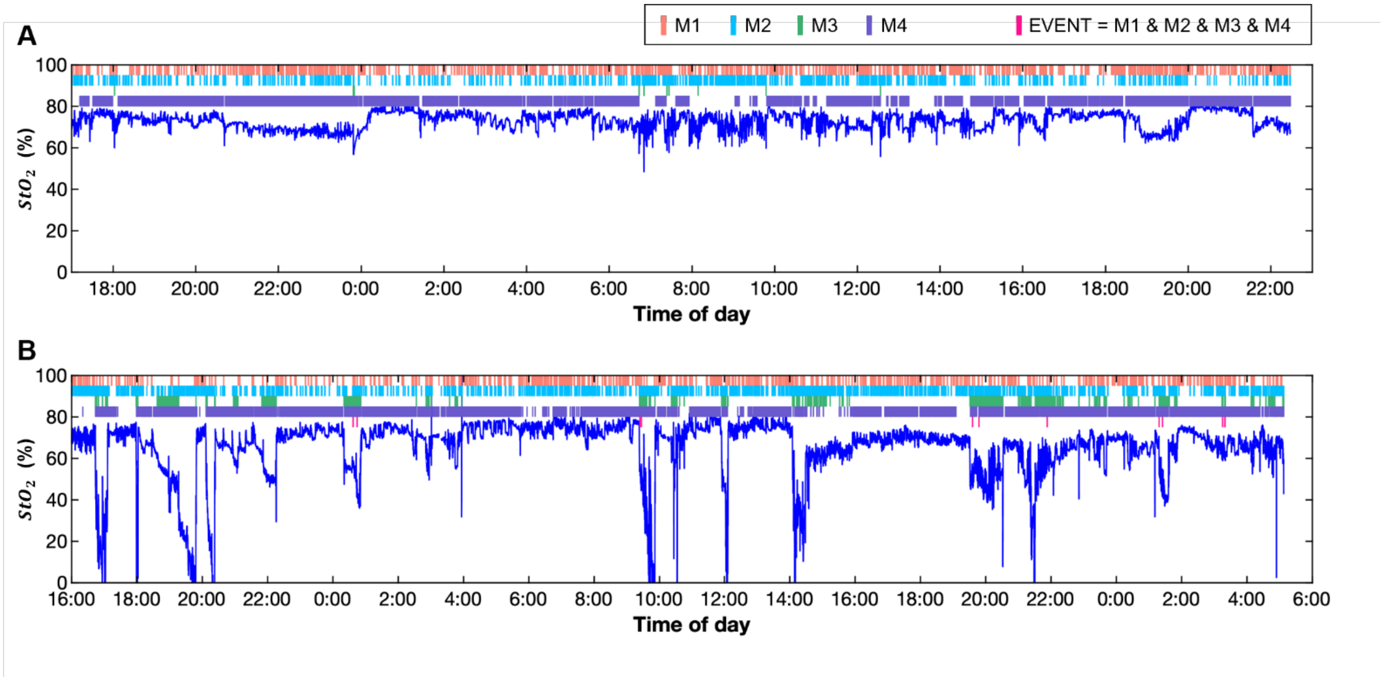


**Fig. S26. Extended recordings of oxygenation ($StO_2$) with Naloximeters in freely moving pigs.** **(A, B)** Colored bars represent instances where the logical metrics (M1 – M4) are true. EVENTS (M1 & M2 & M3 & M4) are registered in (B) but without satisfying the sequential condition to quality for an overdose. $N$ = 2 animals with Naloximeters implanted in **(A)** the neck (jugular approach) for 15 days and **(B)** the flank (mammary approach) for 32 days.

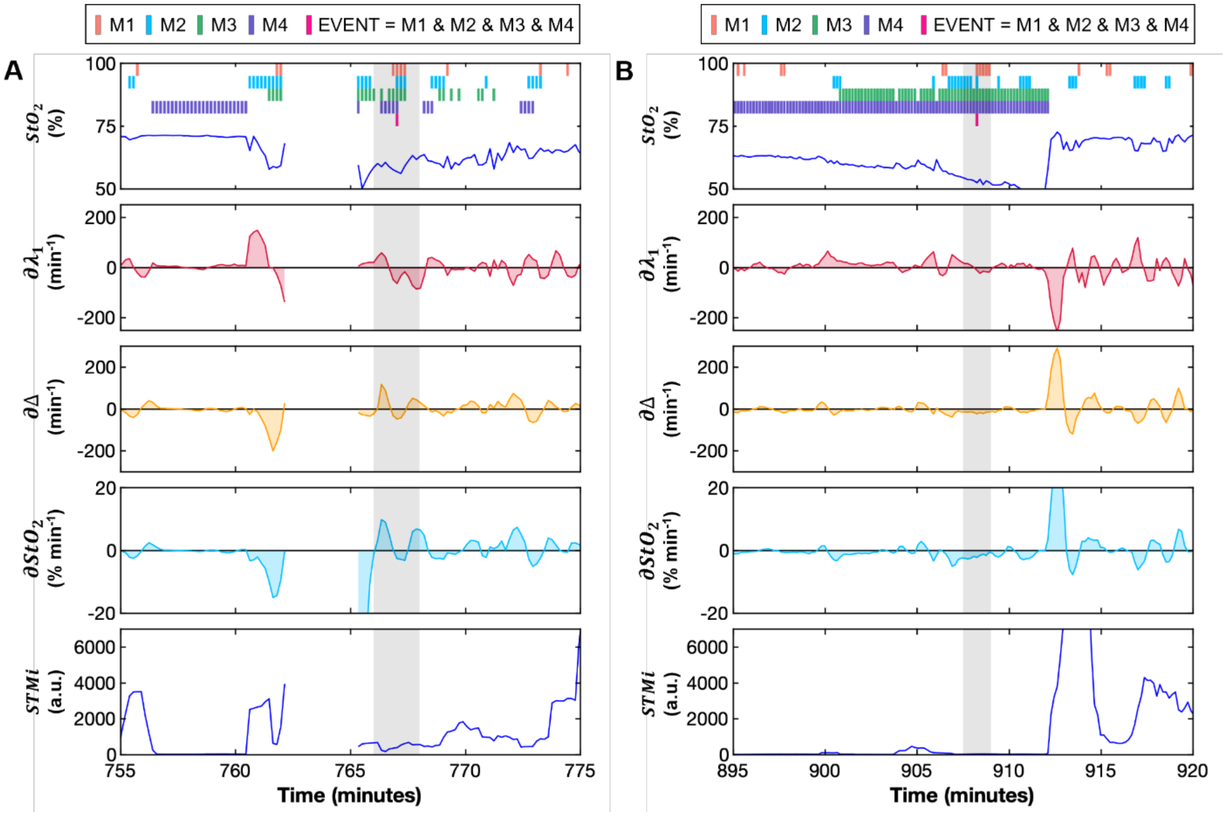


**Fig. S27. Metrics associated with algorithm validation over 24 hours at selected times where an event was detected. (A, B)** From top-down: calculated tissue oxygenation ($\boldsymbol{StO_2}$) showing when the logical metric conditions (M1, M2, M3, and M4) are true; rate of change ($\boldsymbol{\partial}$) of red ($\boldsymbol{\partial\lambda_1}$); difference ($\boldsymbol{\partial}\Delta$); and tissue oxygenation ($\boldsymbol{\partial StO_2}$); short-term motion index ($\boldsymbol{STMi}$). The instances of EVENTs (M1 & M2 & M3 & M4) are isolated and did not produce a WARNING (three consecutive occurrences). Each column represents a distinct episode.

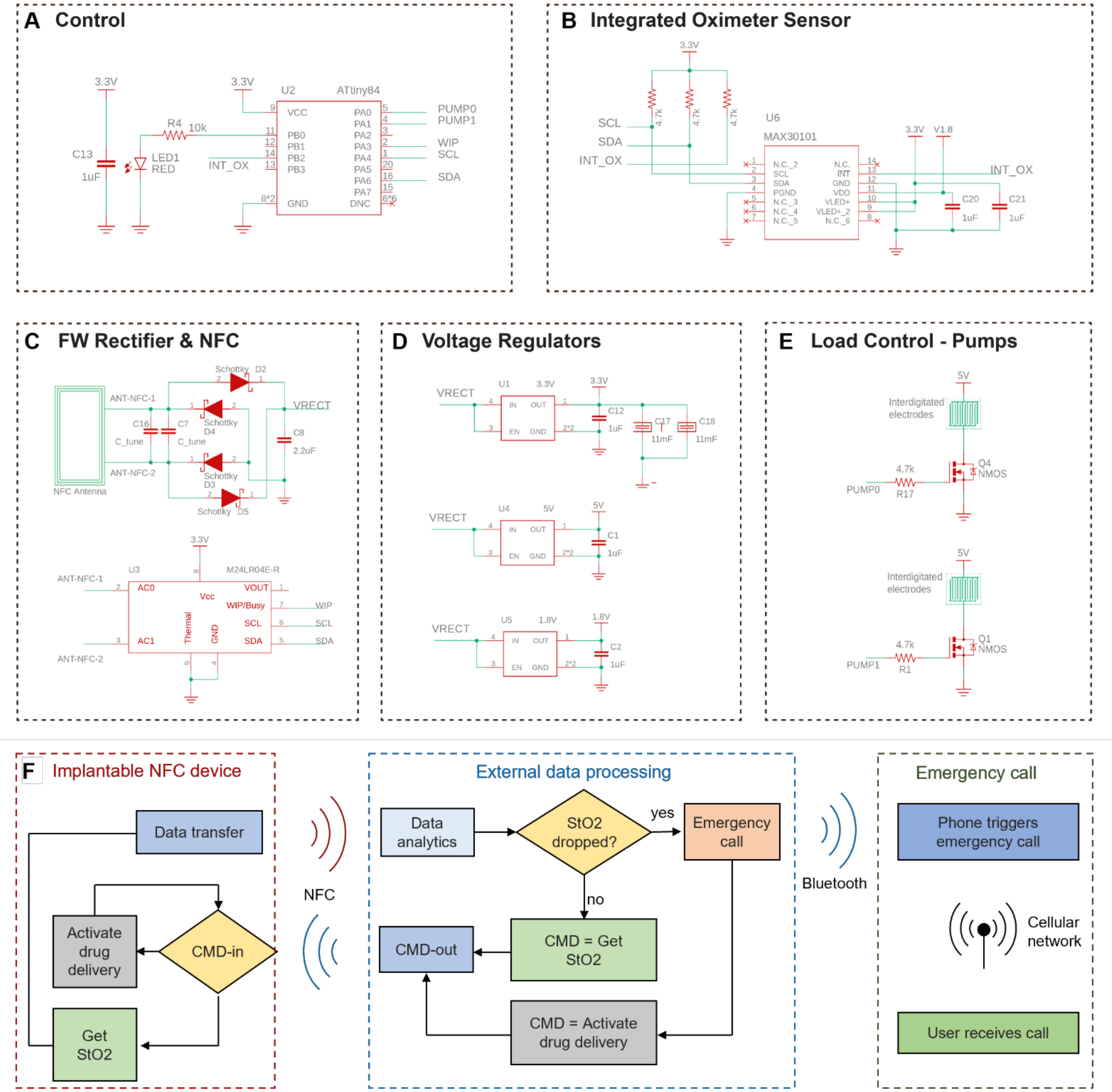


**Fig. S28. Electronic circuit and functional block diagrams for the battery-free, NFC Naloximeter for rodents. (A)** The ATTiny84 microcontroller controls the operation of the device and the manages the wireless communication. **(B)** The integrated optical sensor MAX30101 connects with the MCU using I$^2$C communication. **(C)** RF front end containing a full-wave rectifier and an RF random access memory used for bidirectional NFC communication. **(D)** A voltage regulator supplies 3.3 V to supercapacitor bank (top). A 5 V regulator provides voltage to the electrolytic pumps (center) and a 1.8 V regulator supplies voltage to the dual-wavelength optical sensor (bottom). **(E)** Pump control system with MOSFETs and interdigitated gold electroplated electrodes. Each pump is controlled independently. **(F)** Functional block diagram of the NFC Naloximeter that demonstrates the close-loop implementation in rodents. From left to right: Implanted NFC device; external data processing unit implemented in MATLAB; emergency call management system.

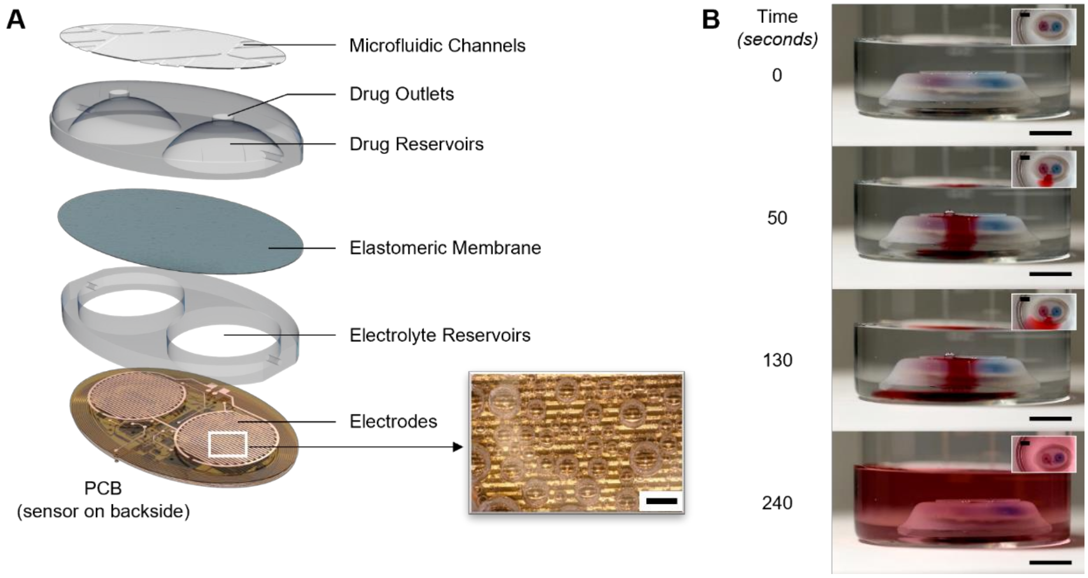


**Fig. S29. Battery-free, NFC Naloximeter for rodents. (A)** Exploded view schematic illustration of the device with inset image of electrolytic gas generation by interdigitated electrodes. Scale bar, 1 mm. **(B)** Photographs that show drug delivery, using food dye as drug proxy for purposes of visualization. Scale bars, 1 cm.

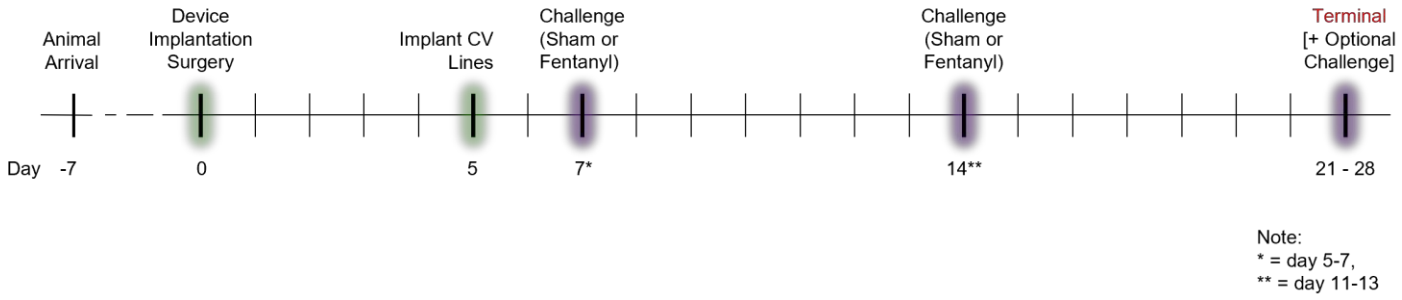


**Fig. S30. General timeline for large animal survival studies.**

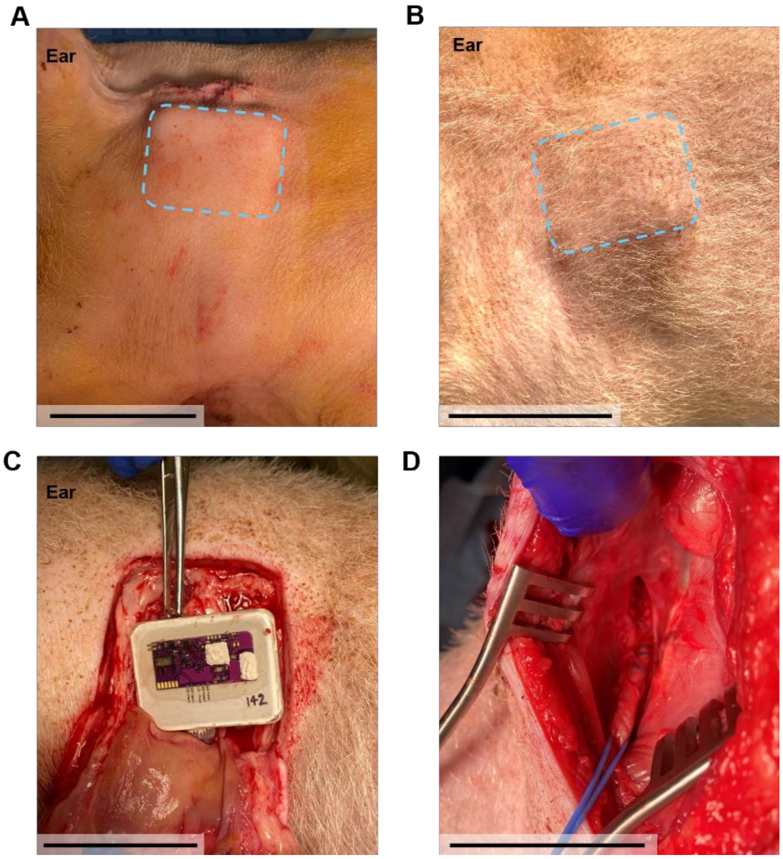


**Fig. S31. Healing progression for an intravenous Naloximeter implanted in the jugular vein of a pig model.** Photographs near the area **(A)** immediately after implantation and **(B)** on day 44 post-implantation. Dashed boxes denote the device outline. Photographs of the **(C)** implanted Naloximeter and **(D)** catheter intact in the jugular vein at day 44 post-implantation. Scale bars, 5 cm.

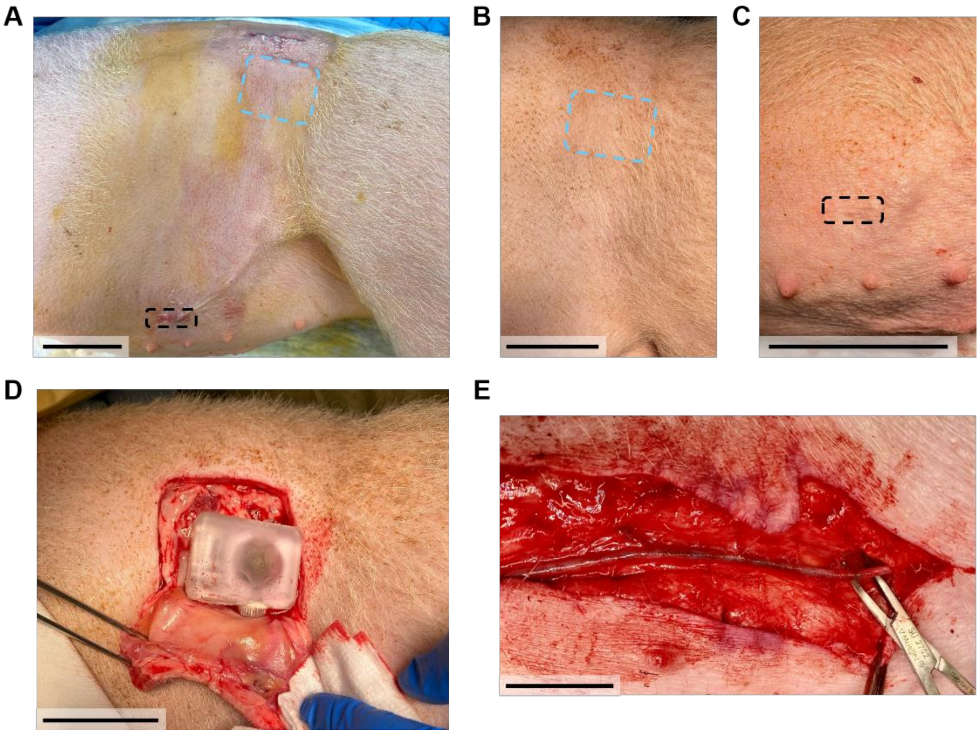


**Fig. S32. Healing progression for an intravenous Naloximeter implanted in the mammary vein.** Photographs near the area **(A)** immediately after implantation and **(B, C)** on day 44 post-implantation at the device and catheter locations, respectively. Dashed blue boxes denote the device outline. The dashed black boxes denote the vascular access incision. Photographs of the **(D)** implanted device and **(E)** catheter intact in the mammary vein at day 44 post-implantation. Scale bars, 5 cm.

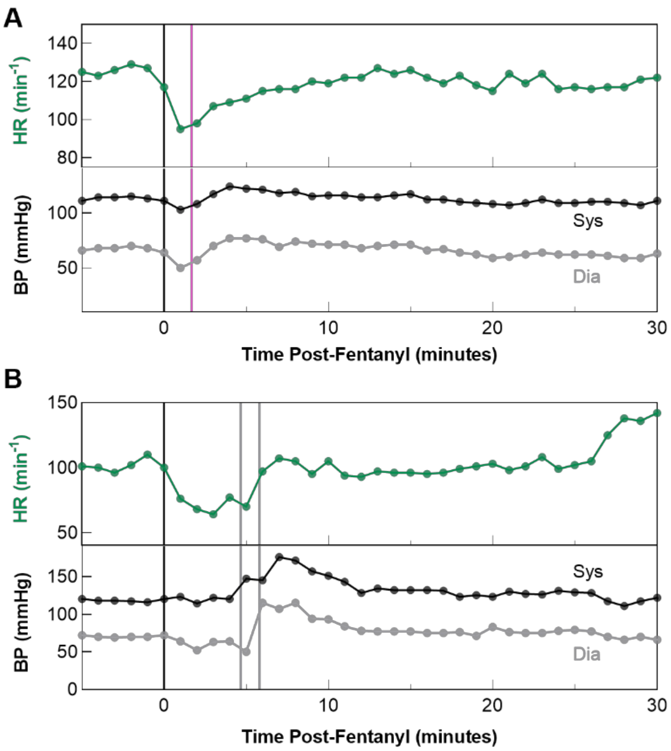


**Fig. S33. Additional vital signs from (A) closed-loop rescue with Naloximeter and (B) the control case without Naloximeter in anesthetized pig model at 2.5 µg/kg fentanyl dose.** Top: Heart rate from electrocardiogram (ECG), bottom: invasive blood pressure. Black line denotes time of fentanyl (i.v.) dose, pink line denotes time of closed-loop NLX (i.v.), gray line denotes time of manual rescue NLX (i.v.). Data in A, B correspond to overdoses in Fig. 4E, 4F, respectively.

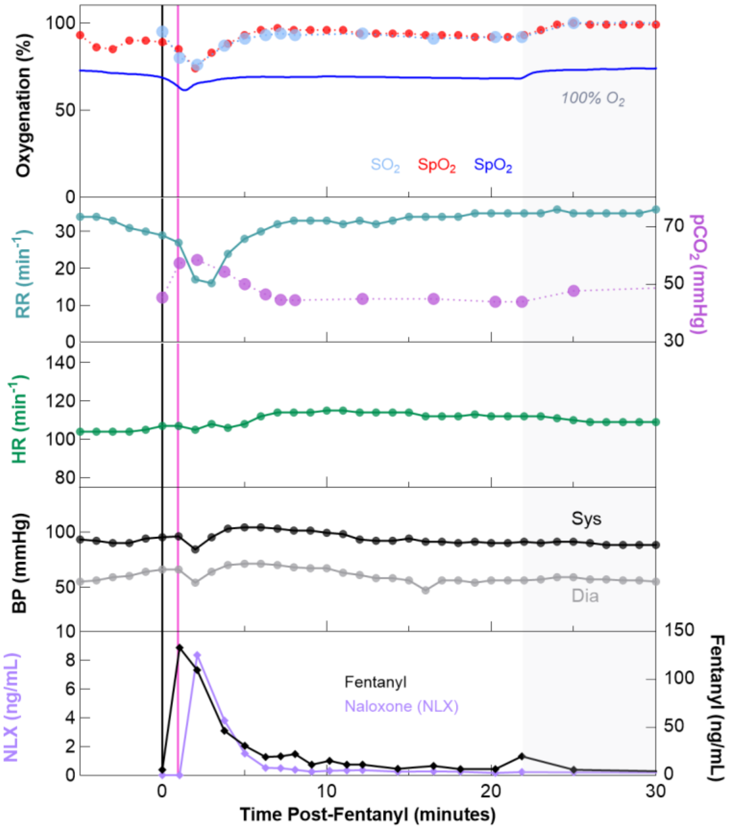


**Fig. S34. Closed-loop rescue demonstration with intravenous Naloximeter in anesthetized pig model at 2.5 µg/kg fentanyl dose.** From top-down: Oxygenation ($SO_2$: Arterial Blood Gas, $SpO_2$: commercial pulse oximeter, $StO_2$: Naloximeter), respiratory vitals, heart rate from electrocardiogram (ECG), invasive blood pressure, and pharmacokinetics of naloxone (NLX) and fentanyl. Black line denotes time of fentanyl (i.v.) dose, pink line denotes time of closed-loop NLX.

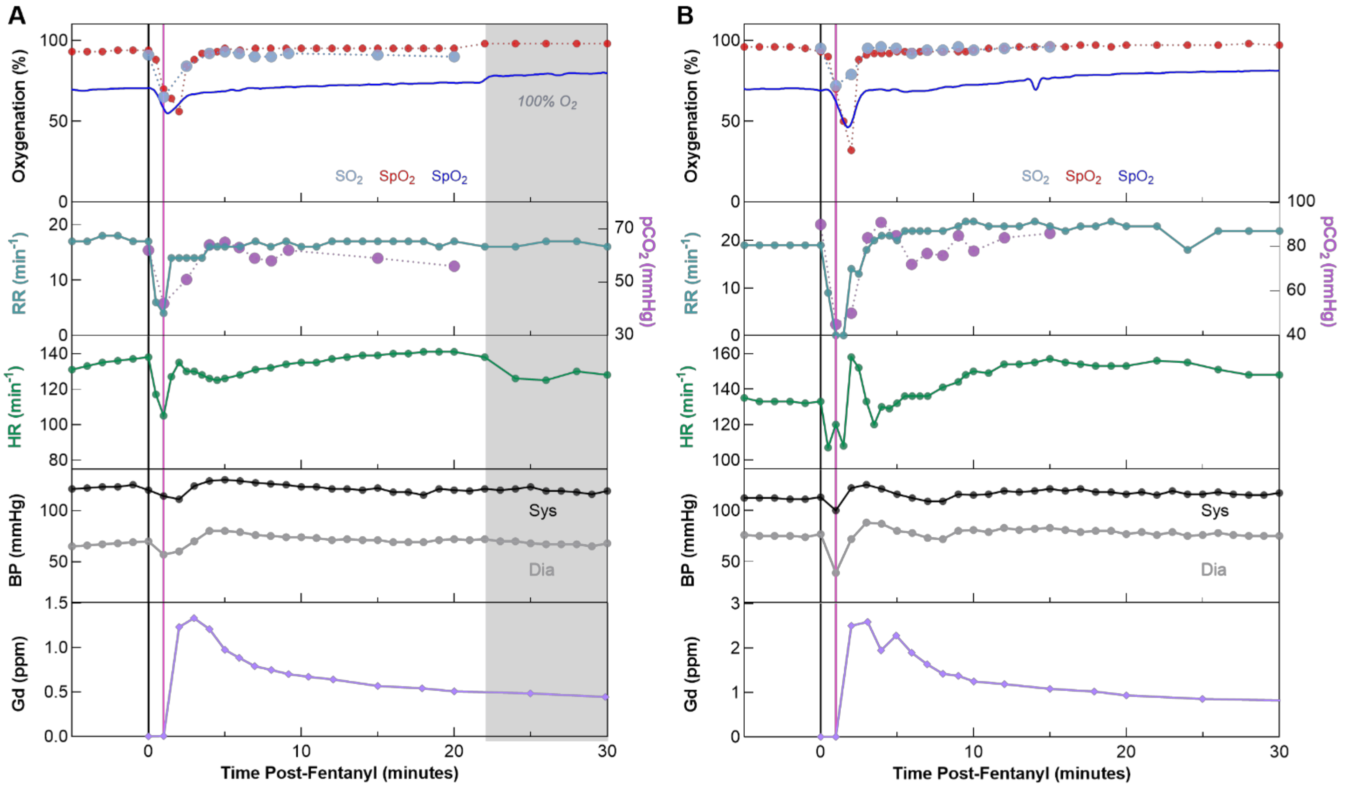


**Fig. S35. Closed-loop rescue demonstrations with intravenous Naloximeter in anesthetized pig model at 5 µg/kg fentanyl dose. (A, B)** From top-down: Oxygenation ($SO_2$: Arterial Blood Gas, $SpO_2$: commercial pulse oximeter, $StO_2$: Naloximeter), respiratory vitals, heart rate from electrocardiogram (ECG), invasive blood pressure, and gadolinium pharmacokinetics. Black line denotes time of fentanyl (i.v.) dose, pink line denotes time of closed-loop NLX (i.v.).

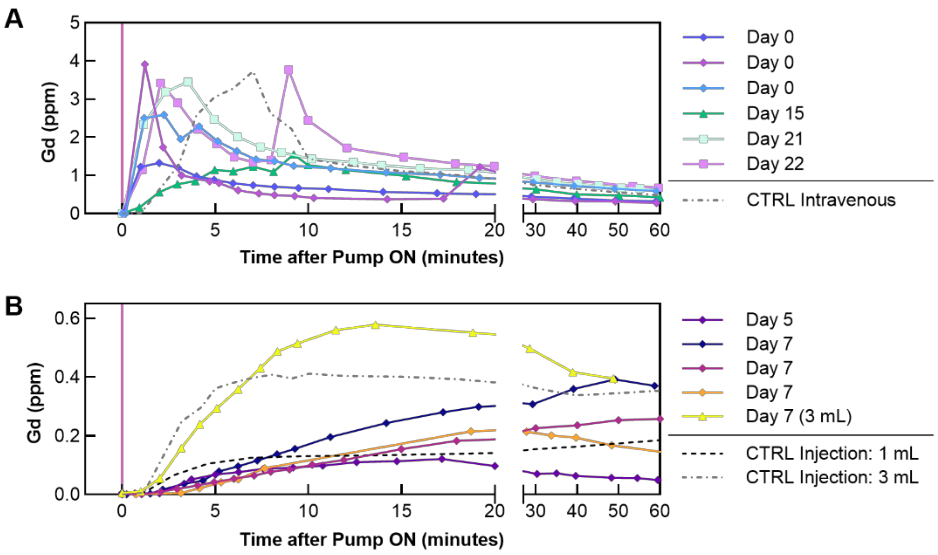


**Fig. S36. Pharmacokinetics of NLX proxy, Gadolinium (Gd), when delivered by the intravenous Naloximeter after various implantation durations. (A)** Gadolinium pharmacokinetics when delivered intravenously by the Naloximeter (mammary and jugular vein) between 0 and 22 days after implantation, or a medical infusion pump in the auricular vein (CTRL Intravenous). Dose is 3 mL. **(B)** Gadolinium pharmacokinetics when delivered subcutaneously by the Naloximeter (without a catheter) between 5 and 7 days after implantation, or by manual subcutaneous injection in tissue surrounding healed devices at 21 and 71 days after implantation (CTRL Injection). The subcutaneous dose is 1.5 mL unless otherwise specified.

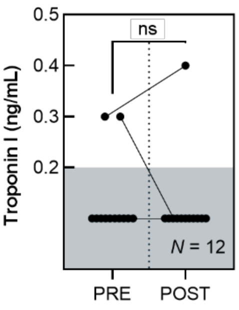


**Fig. S37. Blood biomarkers for cardiac function before (PRE) and after (POST) intravenous Naloximeter drug delivery.** Troponin I quantified from blood serum with a veterinary Troponin I test (IDEXX Laboratories). Values above 0.2 ng/mL are considered elevated, values less than this are plotted within threshold box (lower reporting limit 0.2 ng/mL). Difference is not significant between groups (paired t-test, $P = 0.674$), though one case showed elevated Troponin levels in both PRE and POST samples.

| State | Average current (mA) | Average power (mW) |
|---|---|---|
| Advertising | 0.175 | 0.664 |
| Connection | 1.017 | 3.863 |
| Connected (idle) | 0.200 | 0.758 |
| Connected + Sending Status Updates | 0.231 | 0.877 |
| Connected + Optical Sensor ON | 1.461 | 5.553 |

**Table S1. Summary of power consumption of the Naloximeter for large animals during different operational states.** Average current and power consumption calculated over a 5 s window.

**Movie S1.**

**Demonstration of drug delivery from intravenous Naloximeter device.** Pink dye (Rhodamine B) represents NLX solution, clear container of water represents vasculature. Video speed: 1× and 8× (noted in video).

**Movie S2.**

**Demonstration of drug delivery with injector Naloximeter device, showing dual-injection.** Blue solution indicates NLX solution and yellow solution represents biofluid in the body. Speed: 1×. Scale bar, 1 cm.
